# An accurate genetic clock

**DOI:** 10.1101/020933

**Authors:** David Hamilton

## Abstract

Our method for “Time to most recent common ancestor” TMRCA of genetic trees for the first time deals with natural selection by apriori mathematics and not as a random factor. Bioprocesses such as “kin selection” generate a few overrepresented “singular lineages” while almost all other lineages terminate. This non-uniform branching gives greatly exaggerated TMRCA with current methods. Thus we introduce an inhomogenous stochastic process which will detect singular lineages by asymmetries, whose “reduction” then gives true TMRCA. Reduction implies younger TMRCA, with smaller errors. This gives a new phylogenetic method for computing mutation rates, with results similar to “pedigree” (meiosis) data. Despite these low rates, reduction implies younger TMRCA, with smaller errors. We establish accuracy by a comparison across a wide range of time, indeed this is only y-clock giving consistent results for 500-15,000 ybp. In particular we show that the dominant European y-haplotypes R1a1a & R1b1a2, expand from c3700BC, not reaching Anatolia before c3300BC. This contradicts current clocks dating R1b1a2 to either the Neolithic Near East or Paleo-Europe. However our dates match R1a1a & R1b1a2 found in Yamnaya cemetaries of c3300BC by Svante Pääbo et al, together proving R1a1a & R1b1a2 originates in the Russian Steppes.

## Introduction

**T**he genetic clock, computing *TMRCA* by genetic mutations, was conceived by Emile Zuckerkandl and Linus Pauling[30] on empirical grounds. However work on genetic drift by Motto Kimura[15] gave a theoretical basis and formula. Soon after pioneering work by L.L. Cavalli-Sforza [6], correlated genetic drift to age of lineages for human populations. Suppose at position *j* on the genome is distinguished by number *x* which in the next generation has mutation *x → x±*1 occurring at rate *μ_j_*. Measuring total variance *V* from the mode [22] one finds that the TMRCA = *V/*(∑_*j*_ *μ*_*j*_). This method and variations (denoted as KAPZ) is used to estimate the TMRCA of y(chromosome) haplotypes defined by a SNP (single nucleotide polymorphism) mutation.

In practice sample sizes were too small to compute accurate mutation rates from “meiosis”, i.e. father-son pairs[4]. Alternatively, estimating rates from genetic lineages of known age gave rates with significant discrepancies between different lineages. Indeed for the y-clock these “phylogenetic” rates are often 2 times larger than those from meiosis, while the opposite may be true for other clocks [2], [9], [14], [16].

For the Y-chromosome we show that the mutation rates are essentially constant, at least for the time scale 500- 15,000 ybp, and over different lineages. However KAPZ cannot give accurate TMRCA, i.e. one needs deeper mathematics to deal with non-uniform branching. Also there is a paradox: we can accurately estimate the mutation rates of “short tandem repeat” (STR) at different **D**NA **Y**-chromosome **S**egments (DYS). But we find they can differ by more than a factor of 100, so over a very long time scale we expect their rates to vary as the genomes geometry changes. Also we find knowing the average mutation rate does not give accurate TMRCA.

Of course it was noticed that the mathematics underlying KAPZ is most accurate for large populations, indeed continuous distributions, whereas actual populations are small. In this case the same stochastic model generates many discrete distributions, indicating a need for Bayesian methods. These use Monte-Carlo simulations of all possible genealogical trees giving the present sample data, then find TMRCA by a maximum likehood estimate (MLE). An example of this for the y-clock is BATWING[30]. However we shall see that Bayesian methods exaggerate the TMRCA even more than KAPZ. Also MLE is known for large confidence intervals. So our approach is different.

In particular for the y(chromosome)-clock the results have not been reliable. (Similar discrepancies occur for the mitrochondrial clock for “out of Africa”, or for the allele clock for human-chimpanzee divergence [9], birds [15], bacteria[19].) A KAPZ due to Zhivotovsky [29] was applied to the y-haplotype R1b1a2 by Myres [18] giving 9000BC, standard deviation *σ* = 2000. Now for BATWING the *TMRCA* is often greater than KAPZ, e.g. for the Cinnioglu[7] study of Anatolian DNA both methods were applied to the same data and mutation rates. For R1b1a2 the KAPZ has TMRCA 9800BC compared with 18,000BC for BATWING. Balaresque [3] used BATWING to give an origin for R1b1a2 in Neolithic Anatolia c6000BC, but their statistics was disputed by Busby [5]. In verifying the accuracy of our method we simultaneously resolve the problem of the expansion of European y-haplotypes, for example R1b1a2.

## Singular Lineages

A fundamental problem is that present populations have highly overrepresented branches we call *singular lineages*. A well known example is the SNP L21 which is a branch of R1b1a2. Individuals identified as L21 are often excluded from R1b1a2 analysis because they skew the results. Such a singular lineage causes the variance to be much greater, even though the original *TMRCA* remains unchanged, see figure 1.

**Fig. 1.**
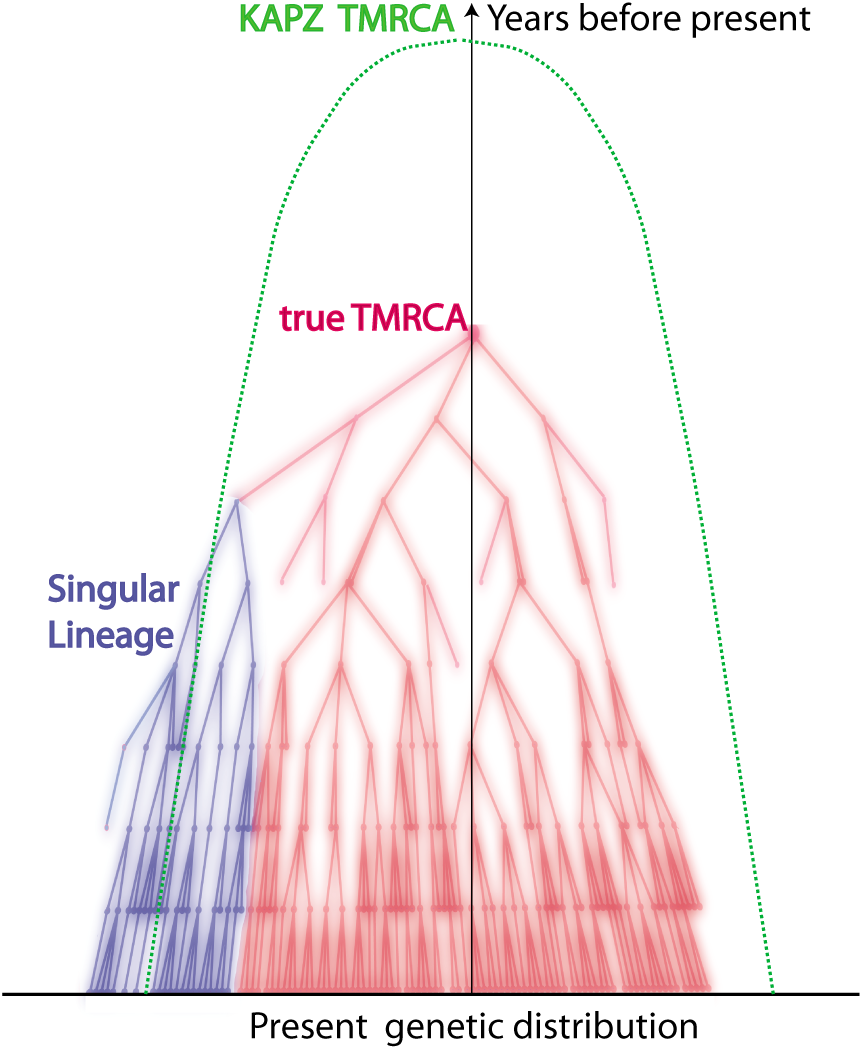
Random tree

For Bayesian methods such lineages are very unlikely giving an even greater apparent *TMRCA*. However one cannot deal with singular branches by excluding them. For one thing, our method will show that 50% of markers show evidence of singular side branches, i.e. more than a SD from expected. Excluding them would also remove some of the oldest branches and produce a *TMRCA* which is too young. Now these singular lineages are very (mathematically) unlikely to arise from the stochastic system which is the mathematical basis of KAPZ (or the equivalent Monte-Carlo process modeling BATWING). We believe that the standard stochastic process is perturbed by “improbable” biological processes.

First, the Watson-Galton process[18] implies lineages almost certainly die out. Conversely, natural selection causes some branches to flourish, e.g. the “kin selection” of W.D. Hamilton[13], shows kin co-operation gives genetic advantages. Consider three examples with well developed DNA projects. Group A of the Hamiltons has approximately 100, 000 descended from a Walter Fitzgilbert c 1300AD. Group A of the Macdonalds has about 700, 000 descendants from Somerfeld c1100AD, and Group A of the O’Niall has over 6 million descendants from Niall of the Seven Hostages, c300AD. These are elite groups with all the social advantages. One sees lines of chieftains, often polygamous. We emphasize kin selection because it seems dominant over natural selection for recent branching, certainly we do not think the O’Niall are genetically superior! Natural selection would cause similar branching over longer time scales. Our model has many extinct twigs with a few successful branches, whereas current models assume a uniform “star radiation”.

## Reduction of Singular Lineages

Although our method is for general molecular clocks to be specific we focus on the y-clock. Consider **D**NA **Y**-chromosome **S**egments (DYS) counting the “short tandem repeat” (STR) number of nucleotides. One uses many of these DYS micro-satellites, marked by *j* = 1*, …N*, each individual *i,* 1 = 1*,…n,* has STR number *x*_*i,j*_. The Y-chromosome is passed unchanged from father to son, except for mutations *x*_*i,j*_ → *x*_*i,j*_±1 occurring at rate *μ*_*j*_.

Modelling singular lineages requires a new stochastic system where instead of a single patriarch we imagine many “virtual patriarchs each originating at a different time and giving a fixed proportion of the present population. Solving for these times and proportions is an inversion problem. But inversion is unstable for such systems, also there is no unique solution. However it turns out that, up to a standard deviation, most DYS markers show at most one singular branch which is found from asymmetries in the distribution. These singular branches are then *reduced* revealing the original lineage. We then compute a branching time *t*_*j*_ for each marker *j*. Now the nonuniform branching process causes the *t*_*j*_ to be randomly distributed so their mean is not the TMRCA see figure 2. Large errors in mutation rates means one cannot simply take the max *t*_*j*_ to be the *TMRCA*. Instead stochastic simulations of the branching process, using robust statistics to avoid outliers, find the most likely *TMRCA*. The effect of reduction is dramatic, e.g. the TMRCA for R1b1a2 changes from 5500BC(KAPZ) to 3700BC after singular reduction, using the same markers and mutation rates, see Figure 3 and Table 1.

**Fig. 2.**
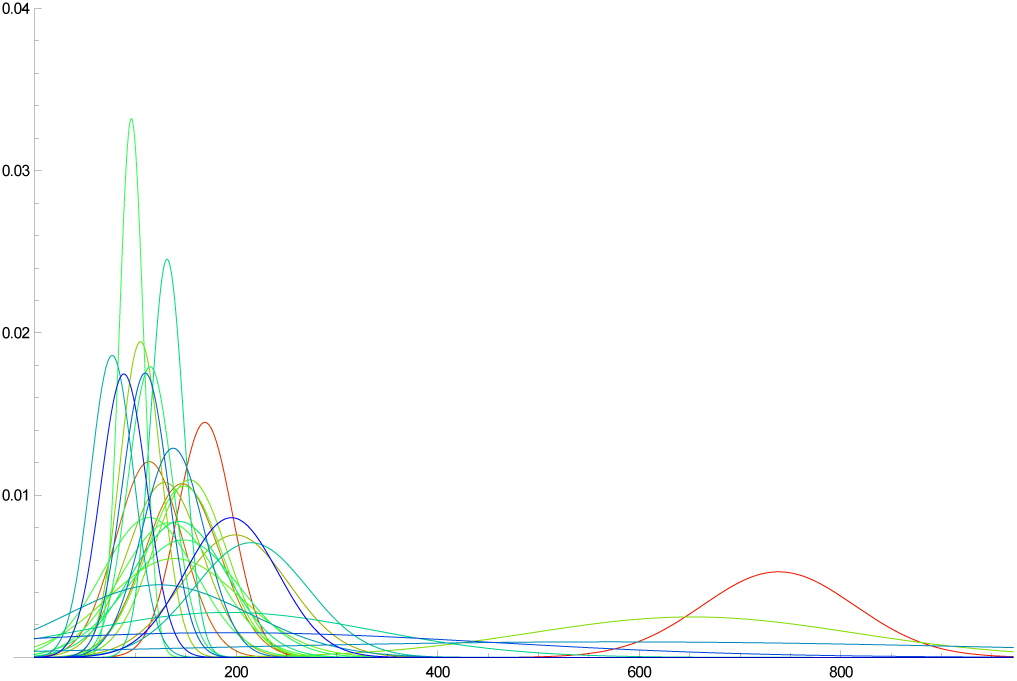
Branching times *t*_*j*_ times(with errors) for R1b1a2 after reduction

**Fig. 3.**
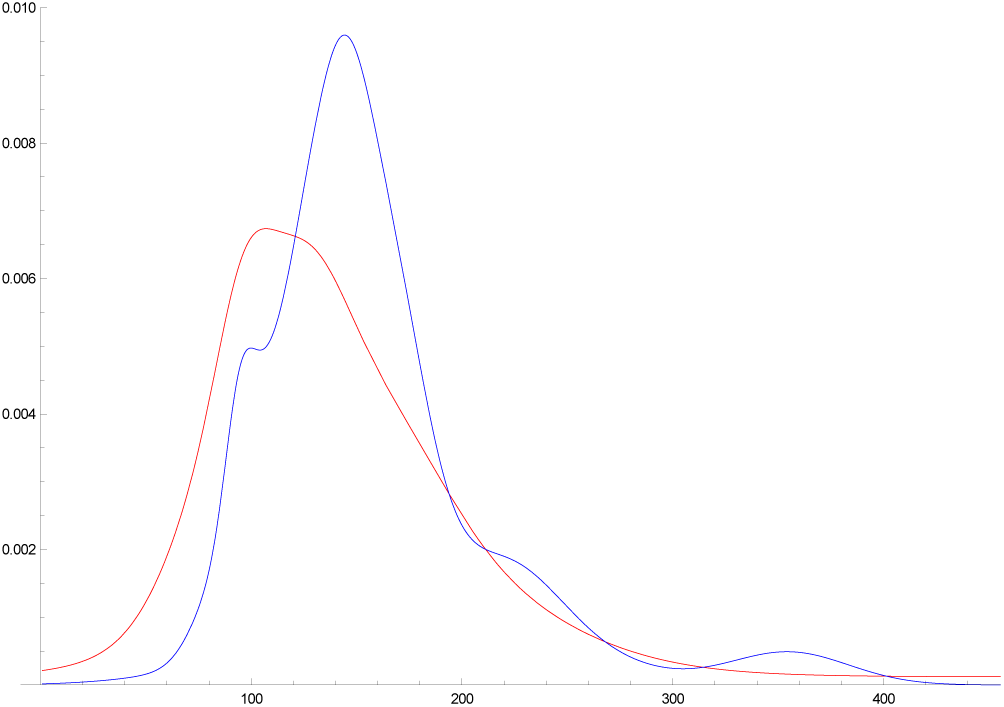
R1b1a2 branching times before(blue) and after (red) reduction

## Accurate Mutation rates

By relying on asymmetries of the distribution to find singular lineages we have to be aware that the mutation process itself might not be symmetric. Indeed if ignored we might be just detecting these asymmetries. So the symmetric model has to be changed so the probability of a mutation is

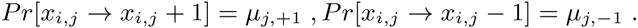

If this marker is free from singular lineages we find that the ratio of the frequencies to the left and right of the mode is

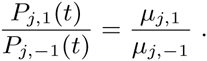

which is time independent. So using eight very large SNP projects we find enough markers free of singularities to compute these ratios and their standard deviations. See Supplementary Information (SI) where Figure 5 shows results. In particular about half the markers show asymmetric ratios are significant, i.e more than two SD from ratio 1. These asymmetric ratios play a very important role, for this ratio is all you need to detect a singular lineage and reduce it. Of course not knowing the exact asymmetric ratio means that bootstrap methods are used extensively for singular reduction, both to compute values and SD.

These methods also imply a new way of computing mutation rates. Previously, there were methods based on meiosos data or phylogenetic studies of family DNA projects (which gave quite different rates). We begin with 8 very large SNP projects from FTDNA using 37 markers, of course with unknown *TMRCA*. We first reduce singular lineages. Then taking asymmetry into account we find mutation rates are the fixed points of an iterative process. This takes about 3 iterates to converge. These mutation rates are normally distributed with mean and SD. Discarding markers with mutation *SD >* 33% leaves us with 29 markers. We find this advanced phylogenetic method gives mutation rates close to those obtained from meiosis and nearly 1*/*2 the values obtained from the usual phylogenetic method. Further validation comes from finding that the equivalence of our rates with meiosos implies *apriori* a human generation of c27 years.

## Results

Accuracy is verified by checking for consistency over the whole range of European history beginning with the medieval:

| Group | *TMRCA* | *[SD]* | Origins |
| --- | --- | --- | --- |
| *Hamilton* | 1358AD | [140] | Fitzgilbert d1330AD |
| *Macdonald* | 900AD | [250] | Norse c800 – 1000AD |
| *O'Niall* | 200AD | [225] | Niall 300AD/Conn100AD |

Archeological finds convinced Marija Gimbutas to attribute Proto Indo-European (PIE) to the Yamnaya Culture c 3500BC of the Russian Steppes, see [12]. This is consistent with mainstream linguistic theory, some even wrote of linguistic DNA. But actual genetics was ignored because current genetic clocks for R1b1a2 pointed to the Renfrew Hypothesis that PIE spread from Neolithic Anatolia, c 6000BC [34]. Or Mesolithic or Paleolithic, depending on the genetic clock. However noone checked if their clock worked over the whole range of time for different lineages.

The next table shows the expansion times of the dominant European y-haplotypes R1b1a2 & R1a1a. These are very close to c3700BC, only Scandinavia is significantly later. This data is from FTDNA projects for region X only using individuals with named ancestor from X. These independent results agree within the standard deviation, with dates matching the Corded Ware Culture, a semi-nomadic people with wagons and horses who expanded west from the Urkraine c3500BC. This is consistent with the oldest R1b1a2, R1a1a skeletons being from the Yamnaya Culture, c 3300BC, see S. Pääbo et al [24].

| Region | *R1b1a2* | *[SD]* | *n* | *R1a1a* | *[SD]* | *n* |
| --- | --- | --- | --- | --- | --- | --- |
| All | 3700BC | [625] | 460 | 3800BC | [700] | 1270 |
| Russia | NA |  |  | 3750BC | [700] | 337 |
| Poland | 3960BC | [950] | 65 | 4600BC | [820] | 876 |
| Germany | 2780BC | [500] | 438 | 3750BC | [800] | 190 |
| Scandinavia | 2550BC | [500] | 153 | 4500BC | [1000] | 140 |

An interesting intermediate step occurs between the medieval and eneolithic. The mythical Irish Chronicles relate that the O’Niall descend directly from the first Gaelic High Kings, which tradition dated c1300-1600BC. The O’Niall have the unique mutation M222 which is a branch of the haplotype L21. For L21, *n* = 1029, we compute *TMRCA* = 1600*BC* and *SD* = 320. These are dates for proto Celtic, i.e. what archeologists call the pre Urnfelder Cultures, c. 1300-1600BC. Furthermore L21 is in turn a branch of haplotype P312 which we date to 2300BC. This date suggests the Bell Beaker Culture of Western Europe. Indeed the only known[24] Bell Beaker genome was found to be P312 with ^14^*C* date 2300BC.

| data | Haplotype | *n* | *TMRCA* | *[SD]* |
| --- | --- | --- | --- | --- |
| Underhill | R1a1a1 | 974 | 2550BC | [400] |
| Rootsi | G2a2 | 536 | 18500BC | [3500] |

Our method requires large data sets and many markers which means we have to rely on data from FTDNA, finding 29 useable markers out of standard 37 they use. In fact many researchers[3] have used FTDNA data. We think our method of reduction with robust statistics solves any problems with this data. To test this we compared our results with R1a1a1 data obtained from Underhill[27] with *n* = 974(which involved excluding his four M420 individuals and others with missing markers), and 15 useable markers. The result was 2550*BC* SD = 400, within the CI of our R1a1a results. Table 1 shows the results of extensive simulations using random subsets of our FTDNA data, for 29, 15 and 7 markers. For the same 15 markers as the Underhill[27] the different FTDNA data gives very similar 3300BC SD = 840 for R1a1a, verifying the correctness of using FTDNA data. However once you get down to 7 markers the confidence interval becomes large, e.g. R1a1a gives 3400*BC* SD = 1500. Also it becomes difficult to deal with outliers.

**Table 1.**
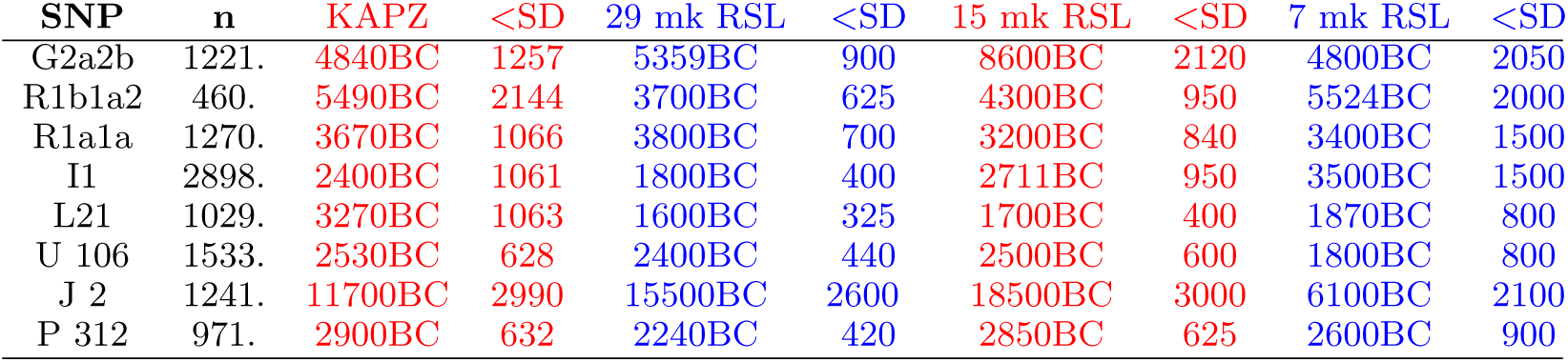
Major European SNP: Comparing Singular Reduction for 7, 15, 29 markers with KAPZ. Notice similar TMRCA for KAPZ and Singular Reduction, if there is little branching.

| SNP | *n* | KAPZ | <SD | 29 mk RSL | <SD | 15 mk RSL | <SD | 7 mk RSL | <SD |
| --- | --- | --- | --- | --- | --- | --- | --- | --- | --- |
| G2a2b | 1221. | 4840BC | 1257 | 5359BC | 900 | 8600BC | 2120 | 4800BC | 2050 |
| R1b1a2 | 460. | 5490BC | 2144 | 3700BC | 625 | 4300BC | 950 | 5524BC | 2000 |
| R1a1a | 1270. | 3670BC | 1066 | 3800BC | 700 | 3200BC | 840 | 3400BC | 1500 |
| I1 | 2898. | 2400BC | 1061 | 1800BC | 400 | 2711BC | 950 | 3500BC | 1500 |
| L21 | 1029. | 3270BC | 1063 | 1600BC | 325 | 1700BC | 400 | 1870BC | 800 |
| U 106 | 1533. | 2530BC | 628 | 2400BC | 440 | 2500BC | 600 | 1800BC | 800 |
| J 2 | 1241. | 11700BC | 2990 | 15500BC | 2600 | 18500BC | 3000 | 6100BC | 2100 |
| P 312 | 971. | 2900BC | 632 | 2240BC | 420 | 2850BC | 625 | 2600BC | 900 |

An example with few markers is the R1b1a2 data of Balaresque[3]. Our method (this time with 7 useable markers) gave SD *>* 30%. Now Balaresque used the Bayesian method BATWING[30] to suggest a Neolithic origin in Anatolia. With the same Cinnioglu[7] data our method gives for Turkish R1b1a2 (*n* = 75) a TMRCA = 5300BC, SD = 3100, i.e. anytime from the Ice Age to the Iron Age as seen in

| *R1b1a2* | *n* | *TMRCA* | *[SD]* |
| --- | --- | --- | --- |
| *Eire* | 75 | 1750BC | [1250] |
| *England* | 74 | 1844BC | [1250] |
| *Spain* | 207 | 4600BC | [1900] |
| *France* | 62 | 4300BC | [2400] |
| *Germany* | 147 | 5650BC | [2300] |
| *Turkey* | 69 | 5300BC | [3100] |

Fortunately, once again, we find good data from FTDNA: the Armenian DNA project, see below. By tradition the Armenians entered Anatolia from the Balkans c1000BC so they might not seem a good example of ancient Anatolian DNA. But some 100 generations of genetic diffusion has resulted in an Armenian distribution of Haplotypes J, G, R1b1a2 closely matching that of all Anatolians, therefore representive of typical Anatolian DNA. We see that Anatolian R1b1a2 arrived after c3300BC, ruling out the Neolithic expansion c6000BC. When dealing with regional haplotypes, e.g. R1b1a2 in Anatolia, the *TMRCA* is only a upper bound for the arrival times, for the genetic spread may be carried by movements of whole peoples from some other region. This means one has to be careful interpreting regional data, e.g. the TMRCA for the R1b1a2(USA) is c3700BC but nobody thinks it arrived then.

| *Armenian* | *n* | *TMRCA* | *[SD]* |
| --- | --- | --- | --- |
| *R1b1a2* | 99 | 3300BC | [800] |
| *G2a2b* | 46 | 9300BC | [2000] |
| *J2* | 97 | 12100BC | [2200] |

Observe that our *TMRCA* for Armenian G2a2b (formerly G2a3) and J2 show them to be the first Neolithic farmers from Anatolia, i.e. older than 7000*BC*. From Table 1 we see J2, G2a2b for all of Western Europe (non-Armenian data). Our dates show J2 was expanding at the end of the Ice Age. Modern J2 is still concentrated in the fertile crescent, but also in disconnected regions across the Mediterranean. The old genetic model predicted a continuous wave of Neolithic farmers settling Europe [8]. But you cannot have a continuous maritime settlement: it must be *leap-frog*. Also repeated resettlement from the Eastern Mediterranean has mixed ancient J2 populations, and our method gives the oldest date. On the other hand G2a2b shows exactly the dates expected from a continuous wave of Neolithic farmers across Central Europe. Our dates are consistent with recent findings that the majority of early Neolithic skeletons found in Western Europe are G2a2, c 5000BC see[33], whereas the oldest R1b1a2 found so far is Bellbeaker c2300BC, [24], [25].

## Discussion

Archeology, evolutionary biology, not to mention epidemiology, forensics and genealogy are just some of the applications of molecular clocks. Unfortunately current clocks have been found to give only “ballpark” estimates. Our method is the only one giving accurate time, at least for the human ychromosome verified over the period 500 *-* 15, 000*ybp*. There should be many applications for this y-clock, not to mention generalizations to mitochondrial and allele clocks.

Some geneticists thought natural selection makes mutation rates too variable to be useful. The problem is confusion between the actual biochemistry giving mutations and superimposed processes like kin selection producing apparently greater rates. Notice that the SD for our mutation rates is on average 14% which is much smaller than the actual previous rates. We believe this proves the reality of neutral mutation rates.

Many applications to genetics, forensics, genealogy require the TMRCA between just two individuals, or between two species, a classic method was given by Walsh[28]. While we are accurate for “big data”, for this “two-body problem” one cannot determine what singular lineages the branching has been through. Just using our new asymmetric mutation rates will not work. So it would be important to find an accurate method.

Pääbo et al[24], [25] observed all 6 skeletons from Yamnaya sites, c 3300BC by ^14^*C* dating, are either R1a1b1 and R1a1a. This and other work [33] involve very difficult genetic analysis of specimens which may not always be available. Also such analysis cannot date the origin of R1a1b1 and R1a1a. Our *TMRCA* shows both these haplotypes expanding at essentially the same time c3700BC. This and our later date for Anatolia, combined with Pääbo et al, implies that R1b1a2 and R1a1a must have originated in the Yamnaya Culture.

In checking accuracy we ran into the question of the origins of PIE. Although there are genes for language there is certainly none for any Indo-European language. Thus inferences have to be indirect. Marija Gimbutas saw patterns in symbolism and burial rituals suggesting the Yamnaya Culture was the cradle of Proto Indo-European. Also their physiology was robustly Europeanoid unlike the gracile skeletons of Neolithic Europe, but this could be nutrition and not genetic. From the above we conclude that the spread of this robust type into Western Europe in the late Neolithic marked an influx of Steppe nomads. Now if R1b1a2 had been shown to spread from Anatolia c6000BC it would have been taken as strong evidence for “out of Anatolia” because of the association of R1b1a2, R1a1 with Indo-European languages. But our accuracy check showed that it was G2a, J2 that spread with the Neolithic Expansion from Anatolia. Now these have been associated with Caucasian languages or Semitic, but never with Indo-European.

## Materials and Methods

This work is biomathematical theory validified by data from publshed sources, see Supplementary Information SI for full mathematical development, data, algorithms and detailed MATHEMATICA worksheets. To verify the theory and compute mutation rates we use diverse data, from FTDNA y-haplotype projects for G2a2b, R1b1a2, R1a, I1, L21, U106, J2, P312. Also we used regional projects for Germany, Scandinavia, Poland and Russia for their R1b1a2, R1a1a data. The Armenian DNA project was important for its R1b1a2, J2 and G2a2b data. We also used DNA projects M222 (O’Niall), Macdonald (Group A which is R1a1a), Hamilton (group A which is I1). This was compared with non FTDNA data from Balaresque, Underhill and Rootsi.

## ONLINE TEXT: Supplementary Information

1. Biomathematical theory
2. Mutation rates tables
3. Mathematica worksheets
4. Data

## Biomathematical theory

We emphasize the role of extraneous forces like kin-selection which operates on too big a scale and rarely enough with results that cannot be subsumed into the mutation rates. So we return to basic principles.

### Fundamental Solutions

The Y-chromosome has DYS marked by *j* = 1, …*N*, where one can count the STR number *x*_*j*_. Consider the probability *P*_*j,k*_ (at time *t* generations) that at marker *j* we have *x*_*j*_ = *k*. This satisfies the homogenous stochastic system

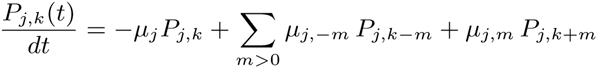

This homogenous system gives a uniform expansion from a single patriarch.

The system is essentially the model of Wehrhahn[29] who had *μ*_*j*,-1_ = *μ*_*j*,1_. We introduce asymmetric mutations with total rate

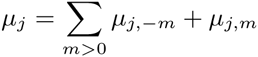

About 50% of DYS markers show asymmetric mutations, i.e. 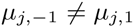.

The fundamental solution comes from the generator function

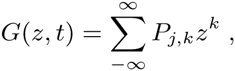

with complex variable *z*, and normalized initial condition *x*_*j*_ = 0 or *P*_*j*,0_(0) = 1:

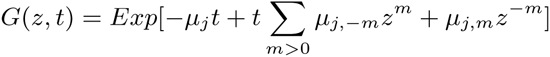

Then *G* can be expanded in powers of *z* to give *P*_*j,k*_(*t*). Now for the simplest asymmetric case, with only one step mutations, we have 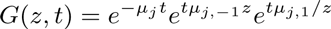 =

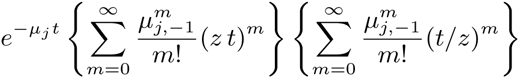

so using the Hyperbolic Bessel Function of Order 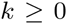, see Olver^7^

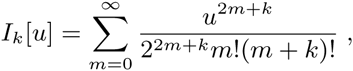

we see that the homogenous system has fundamental solution

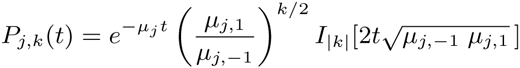

From this we obtain the second moment:

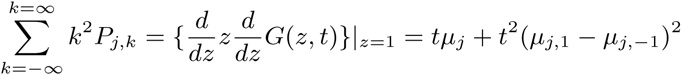

Also from the fundamental solution we find, independently of time

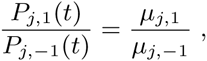

which we call the *asymmetric ratio*. It will be repeatedly used.

Of course the actual initial value is not *x*_*j*_ = 0 but was usually taken to be the mode *m*_*j*_ which was assumed to be the value for original patriarch. Assuming symmetry, i.e. *μ*_*j*,-1_ = *μ*_*j*,1_, the TMRCA is:

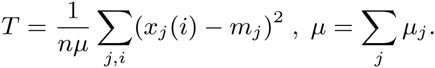

From the present distribution of data we use the frequency

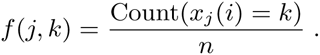

One problem with the KAPZ formula is that higher frequencies *f*(*j, k*)*, |k|* = 2, 3*…* are overrepresented in the actual data. This is because the probability of a spontaneous two step mutation is much higher then the product of two one step mutations. So instead we use the frequency to solve the transcendental equation for the unknown *t*

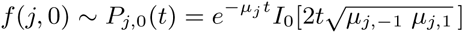

This nonlinear equation is easily solved via mathematical software such as MATHEMATICA (I used version 9 running on a boosted 2014 iMac which has accurate hyperbolic Bessel functions. Earlier versions on older iMacs gave inaccuracies so one had to compile one’s own functions). Using this formula resolves some other problems with the KAPZ method, e.g. 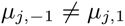 gives an extra quadratic term which if ignored causes large errors.

### Heterogeneous diffusion equation

However the main problem is singularities in the stochastic process. For a uniform stochastic process, 1−*P*_*j*,0_(*t*) ∼ 1−*f*(*j*, 0) is the probability of some mutation. So the expected variance is *f*(*j*, 0)(1-*f*(*j*, 0)). Thus if the actual data variance *V*_*j*_ >> *f*(*j*, 0)(1 − *f*(*j*, 0)) we are not uniform. Now a sublineage of very high fertility increases variance, giving apparently greater *TMRCA* although it is unchanged. One finds similar results for Bayesian methods.

The correct approach to nonuniformity assumes at times *t*_*i*_ (generations ago) a certain proportion 0 ≤ *ρ*_*i*_ ≤ 1 of the present population originated from a “virtual patriarch” with an initial STR value *m*_*i*_. The resulting system :

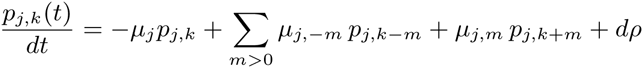

i.e. *dρ* are atoms of weight *ρ*_*i*_ with STR value *m*_*i*_ occurring at time *t*_*i*_. As the system is linear and isotropic the solution is a combination of fundamental solutions *P* of the homogenous system. Thus the present distribution *f*(*j, k*) is

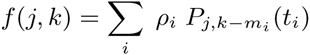

This allows us to consider populations mixed by having singular lineages from overfertile patriarchs, or by actual immigration from the outside. The inverse problem seeks to find singularities from present data. Unfortunately inversion is ill posed for such systems like the heat equation. This instability produces poor accuracy. Furthermore there is no unique solution, e.g.the present distribution could have been created yesterday.

However we find that *∼* 50% of the DYS markers show no significant difference from the uniform expansion of a single patriarch, i.e. the data variance *V*_*j*_ is close to the expected variance *f*(*j,* 0)(1 *- f*(*j,* 0)). The other markers show at most one significant side branch, i.e. there is an original branch starting at time *t*_*j,*0_ with STR *m*_0_ and a second one with STR *m*_1_ = *m*_0_ *±* 1 at time *t*_*j*,1_ < *t*_*j*,0_ with significant 0 *< ρ*_1_ *< ρ*_0_.

### Reduction

We locate these singular lineages by looking for asymmetries in the distribution. For a uniform flow from a single patriarch the frequency of STR value *k* is given by *f*(*j, k*) ∼ *P*_*j,k*_(*t*). The asymmetric ratio:

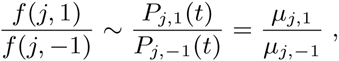

is completely independent of time *t*. Therefore if say

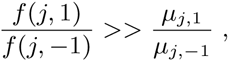

we have a singular lineage at *k* = +1. Thus the excess at *k* = +1 is

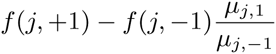

To first order approximation then frequency *f*(*j,* +2) is due to this singularity at *j* = +1 which therefore gave a contribution

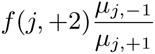

to *k* = 0. Thus removing the effect of the singularity at *k* = +1 leads to new frequencies

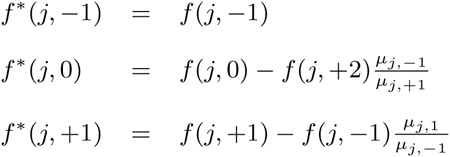

These of course are no longer normalized so we rescale to obtain the renormalized frequency *F* (*j, k*), e.g.

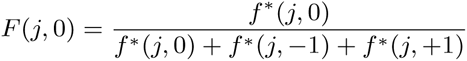

which will be used to compute the expansion time for marker *j*. There are similar formulae if the singularity was at *k* = *-*1.

However there is sampling error both in the frequencies and the *μ*_*j*,1_,μ_*j*,-1_. So we bootstrap taking into account these uncertainties, running the computation thousands of times. Generally we find the branch singularity is always one of *k* = 0, +1*, -*1 with no SD. In a few cases the singularity may seem to wander between *k* = 0, +1*, -*1. So in the case of a wandering singularity we obtain a distribution over *k* = 0, +1*, -*1 with a mean and SD. In these cases we find the singularity is relatively small and does not make much difference to the final result. However to have a stable method we do not throw out these wandering singularities but in the algorithm use the mean to average between *k* = 0 and *k* = *±*1, e.g. if the mean is *k* = 0 then we use the original unreduced frequency.

Notice that we assume at most one side branch. In theory there could be many and solving for these produce even better approximations to the present data. In fact you could get perfect matching but find the atoms were created yesterday! The thing is that while many markers show significant deviation from a uniform flow from a single patriarch, after we have carried out reduction for one possible side branch we find no significant difference from a uniform flow, i.e. the difference is within the SD. This is of course an approximation, the next level beyond Zuckerkandl and Pauling, but given the noise in the data perhaps the best we can do. Later we further reduce the effect of outliers by using robust statistics.

Reducing the singular lineages increases the frequency *f*(*j,* 0) of the mode and decreases the computed *TMRCA*. But as the method of reducing singularities does not respect higher frequencies *f*(*j, k*) it follows the KAPZ formula cannot be used and instead we use the probability of no mutations, i.e. solve

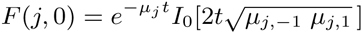

This is done for each DYS marker *j*, giving expansion times *t*_1_*, …t_N_* for each marker, with computed CI. (An extra fixed source of error is the uncertainty in the mutation rates which we deal with later). We find the reduction of singularities makes striking difference to the *t*_*j*_ of the effected markers, often a reduction of *∼* 50% for *TMRCA*.

Now the existence of side branches implies that the main branch could itself have been the side branch for an earlier branch that did not survive. Thus we do not expect the expansion times *t*_1_*, …t_N_* for each marker to be essentially equal., i.e they are not within the SD of each other. Indeed we see that the distribution of the times *t*_*j*_ for different markers are almost certainly not randomly arranged about a single *TRMCA T* but distributed from *T* to the present. This is seen whether you use reduction or not, or our mutation rates or not. (For a given population one could scale mutation rates to get equal *t*_*j*_, but then applying these adhoc mutation rates to other populations does not yield the same values). The spread out distribution of surviving branches is another verification of our theory of many extinctions, few survivors. The distribution of the times *t*_*j*_ for different markers we call the branching distribution, which is now discussed.

### The Branching Distribution

The times *t*_*j*_ for different markers are sorted from the youngest to the oldest, forming a sequence 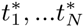. The generation of these branches is by an unknown probability distribution *dτ*_0_ over [0*,T*]. We model *dτ*_0_ by assuming a surviving lineage is generated at random with probability *β*Δ*t* in time period [*t, t* + Δ*t*], multiplied by the probability that the branching hasn’t already occurred. The constant *β* averages fertility and extinction rates, the chance of a new lineage surviving. As *β → ∞* we get current theory where all lineages originate from a single patriarch at time *T*. Simulations with the data show that *β* varies in the range 1 to *∞*. We make no a priori estimate of *β*, unlike Bayesian methods where an overall fertility rate is a predetermined parameter. Instead our stochastic simulation will find the most likely *β, T* in each case. Assuming independence, then the generation of branches follows the well known exponential distribution:

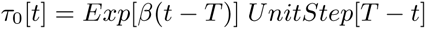

Notice this implies a finite probability that some markers have essentially zero mutations. This is actually seen in examples. Both the Hamilton Gp A and Macdonald Gp A have number of individuals *n >* 100. For the time scale of *>* 700 years we do not expect there is more than one marker out of 33 which shows absolutely no mutations from the mode. In fact in both cases there are 8 markers where all *n* individuals have exactly the same STR value.

Estimating the parameter *T* for an exponential distribution is a well known problem of statistics. Kendall proved the best estimate for *T* would be max *t*_*j*_. Unfortunately there is also considerable error *λ_j_*% for the mutation rates *μ_j_*. Later we give a method for reducing this error, even so we find the SD in the range 10%*-*30% which gives corresponding range in error for each *t*_*j*_. We understand that the *t*_*j*_ are being generated by the distribution *dτ*_0_ but superimposed on this is a further uncertainty due to mutation rates etc. In particular the largest *t*_*j*_ may be wildly inaccurate. Also we found that simply taking the average consistently underestimates the *TMRCA* by a wide margin.

Assuming the mutation rates have normal distribution with mean *μ_j_* and variance *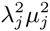*, the *t*_*j*_ have SD *t*_*j*_λ_*j*_. Thus the actual data for 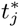 has probability density function for *s >* 0

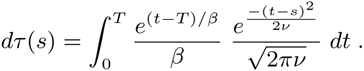

The variance *v* depends on two sources. First from the uncertainty in mutation rates, for each marker we get variance 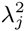, giving total

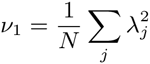

However a small sample also has inherent error from sampling. We are measuring the probability that there is a mutation. This is binomial with probability *H*_*j*_ = *H*_*j*_(*t*) =

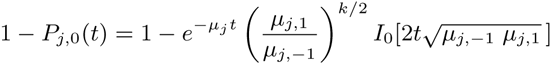

Hence for sample size *n* there is variance *H*_*j*_(1 *- H_j_*)*/n*, so the variance in time due to this is scaled by the derivative giving:

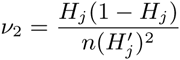

The function 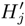 has actually to be computed as an inverse function depending on *H*_*j*_. Therefore the total variance averaged over all *N* markers is *ν* = *ν*_1_ + *ν*_2_. Although for large samples (*n* > 1000) the second term is insignificant it does effect the results once you get to *n* = 100. In our algorithm the branching distribution is used to generate large numbers of random branching times so as to bootstrap error estimates. In turns out much faster to compile the distribution function as a table which can be repeatedly called on.

### Estimating TMRCA by Robust Statistics

Inaccurate large values of 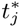 are mitigated by using “robust” statistics with quintiles instead of means/variances. Using FTDNA data we began with 37 markers. However the 4 markers of DYS464 are unordered and cannot be used. Also we find that markers DYS 19/394, 385b, 459b, CDYb have errors *>* 33% in mutation rates so are not used. (These are some of the most popular ones in the literature!). So usually we have *N* = 29 markers and take “quintiles” 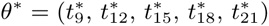. This means that tail end data is not discarded but kept as the information there are 8 values of 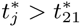, which effectively deals with outliers. Bootstrap methods give the confidence interval CI for each quintile.

Thus we wish to find the best estimate of *T* given *θ*^*^ (and CI). This well known statistical problem was investigated by Stochastic Simulations (SS). We also tried Maximum Like-hood Methods which gave similar results but with larger CI. Monte-Carlo Methods are used to produce very large numbers (*∼* 10^7^) of *T, β* with corresponding Distribution. These randomly generate ordered times (*s*_1_*…s*_29_) for which we take the quintiles *θ* = (*s*_9_*, s*_12_*, s*_15_*,s*_18_*, s*_21_). We filter by requiring that *θ* close to the data *θ^*^*, i.e. *‖θ^*^ - θ‖ <* 2*SD*. This gives a stochastic neighborhood *U* of *θ^*^* typically containing *>* 10^5^ sets of data but with *T* is known for each *θ ∈ U*. Thus we can construct a quasilinear estimator:

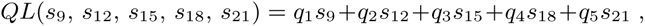

and use least squares over *U* to find constants (*q*_1_*, q*_2_*, q*_3_*, q*_4_*, q*_5_) minimizing

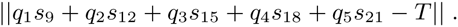

The (*q*_1_*, q*_2_*, q*_3_*, q*_4_*, q*_5_) are computed in MATHEMATICA. We test this by applying the QL to all of *U*, unsurprisingly

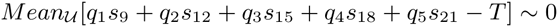

What is important is that we find the uncertainty in the SS itself. Actually this depends on the data and is calculated in each case but for our examples we find

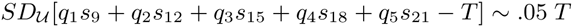

Finally the quasilinear estimator is applied to the experimental data

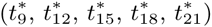

to obtain our best estimate of *T*. Application of *QL* computes the SD for our data, giving part of the overall SD. This must be combined with the SD coming from the uncertainty in the SS. Overall we find that our method has SD *∼* 12%, this includes variances from our data, mutation rates and uncertainty in the SS. We also tested with 15 and 7 markers. Here one must use “quintiles” 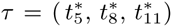, 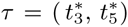, respectively with all the loss of accuracy that implies. See Table 1 for comparisons using 29, 15, 7 markers on same data.

### Accurate Mutation rates

**Any genetic clock depends on reasonably accurate mutation rates. The meiosos method looks for mutations in father-son studies. However typical rates of** *μ* = .002 **would require nearly** 50, 000 **pairs to get an SD of** 10%**. Small samples have meant large errors. The phylogenetic approach studies large family groups with well developed DNA/genealogy data. So inverting the KAPZ formula would yield accurate rates. However,** *singular lineages* **makes this problematic. Genealogical data might give mutation rates much greater than the biochemical rates because kin selection etc tend to exaggerate the apparent mutation rate. An inspection of** 10 **different sources finds mutation rates claiming SD** *∼* 10% **yet they differ from each other by up to** 100%**. We describe a new method.**

To compute our rates we apply our theory to the large DNA projects for the SNP M222, L21, P312, U106, R1b1a2, I1, R1a1a. This avoids dealing with populations such as family DNA projects which are self selecting, i.e only those with the correct surname which neglects distant branches. Also we have very large samples, our average *n >* 1000. Greater accuracy should come from more generations and individuals. The problem is that we do not know their *TMRCA*.

### Asymmetric Mutation

However before computing mutation rates we must consider asymmetric mutations, i.e. the left and right mutation rates 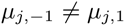. For a uniform stochastic process we again use the asymmetric ratio

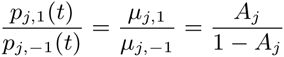

to define the *asymmetric constant A_j_* ∈ [0, 1] for marker *j*. For example *A*_*j*_ = 0.5 is complete symmetry. Of course singularities will effect this ratio, however these only occur *<* 50% of markers. Thus for each marker, SNP we compute this ratio. We find the SD for each SNP is relatively small while the difference between SNP can be large. However for each marker, using 8 SNP enables outliers to be easily removed leaving allowing us to use simple linear regression: i.e. average of the *A*_*j*_ over the remaining SNP groups. We see that asymmetry is a real effect: 50% of the *A*_*j*_ are more than two SD from symmetry *A*_*j*_ = 0.5.

Observe this is significant. The total second moment is

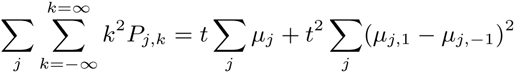

So using all our 33 DYS markers with our *μ*_*j*_, we compute constants

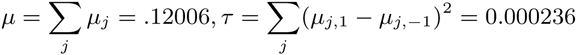

The KAPZ formula gives variance *V* = *μt* compared to the corrected formula *μt* + *τt*^2^. The uncorrected KAPZ gives an overestimate *>* 400% for *>* 200 generations. This effect can be nullified by using the mean instead of the mode, variance instead of the second moment, however failing to do so gives a large error. Furthermore other methods which assume symmetric mutations will also be inaccurate. Having estimates on the asymmetry is essential to our method because we find singular lineages by looking for asymmetry in the data. Any such anomaly needs to be significantly greater than the natural asymmetry.

### Mutation Rates as a fixed Point

Next we compute mutation rates using 8 very large SNP groups. First, using the asymmetric constants we find singular lineages and reduce their effect. We take account of the error in the *A*_*j*_ by a bootstrap technique, which gives the variance for each frequency *f*(*j,* 0). For a given SNP *k* if markers *j* started their expansion at the same time TMRCA *T*_*k*_ we could calculate mutation rates *μ*_*j*_ via

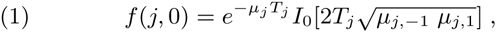

or rather average the 8 different *μ_j_* we would obtain. However because of branching caused by extinction of lineages the different markers do not originate at the same time but at different times *t*_*j*_. In this case we expect these *t*_*j*_ to be randomly distributed about the log mean over a middle set of times *t*_*j*_. So, for each SNP group *k* = 1*,‥*8 define mean time *T*_*k*_, not the TMRCA but the mean log mean over a middle set of markers, which is less. We find that this is very stable. So for a fixed marker *j* the data *τ_k,j_* = *t*_*j*_ - *T*_*k*_ should be randomly distributed about zero over the different SNP *k* = 1*, ‥,* 8. However the wrong choose of *μ_j_* would give a bias. In fact this is what we see if the mutation rates *μ_j_* = .002 were chosen. In appendix graphs show the *τ_k,j_, k* = 1*,‥*8 bunched around a nonzero point. Thus we try to find *μ_j_* so that the *τ_k,j_, k* = 1, 2*,‥*8 has mean zero. However the *τ_k,j_, k* = 1, 2*,‥*8 depend nonlinearly on the rates *μ_j_*, as does the mean *T*_*k*_,*k* = 1*,‥*8. We find this nonlinear regression problem is solved by an iterative scheme which starts with any reasonable set of DNA rates, finding any reasonable choice iterates to the same final answer. So choose *μ_j_* = .002 to begin. Suppose at some stage we have apparent mutation rates *μ_j_*. Then, for each SNP, and each marker we solve equation (1) to obtain the apparent *t*_*j*_. For each SNP *k* = 1*,‥*8 we compute the mean log time *T*_*k*_. At the next step we get new rates *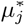* from

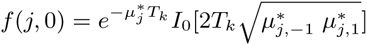

Averaging 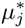, *k* = 1, ‥8 we get our next set of *μ_j_* of mutation rates. However this method would be effected by a marker showing a singular lineage. Fortunately these are few in number and by comparison between the different SNP we remove the outliers. We then repeat the process, computing *T*_*k*_ again with the new rates, and another set of mutation rates. So we have an iterative process.

One problem is that the iterates could tend to decrease to zero or increase to *∞*, as we are only calculating relative rates. To prevent this we renormalize after each iteration so the total ∑*μ_j_* is constant. We found the iterative scheme quickly converges to a fixed set of mutation rates, unique up to a constant factor. The CI is computed by bootstrap parametrized by the uncertainties in data and the asymmetric constants.

### The generation factor *γ*

This method does not give absolute mutation rates but *relative* mutation rates *μ_j_γ*,where *γ* is universal time scale constant. To find *γ* we apply our method to compute the *T* = *TMRCA* of three famous DNA projects and choose *γ* so the scaled *T/γ* best fits the historical record. We choose the DNA projects for the O’Niall(M222), Gp A of Macdonald (R1a1a) and Gp A of the Hamiltons (I1). These are large groups with characteristic DNA and fairly accurate times of origin. Of course finding one constant *γ* from three projects is inherently more accurate than using one project to find 33 different mutation rates. Actually assuming a generation of 27*years* these three projects yield *γ* = 1 with about 5% error, i.e. there is no actual need for this correction. This is a constant error (like uncalibrated ^14^*C* dating).

Thus *γ* is related to the length of a generation. Most researchers use 25*yrs* for *t >* 500*ybp* and 27*yrs* for *t <* 500*ybp*. Balaresque and al used 30*yrs* based on Fenner [11] who sees a 30*yr* generation for modern hunter-gatherers. Our theory allows any nominal generation as it really doesn’t matter, being included in the *γ* factor which we compute in years not generations. However to give actual mutation rates we need an actual generation so we take 27 years. This appears in our worksheet computation. Notice that choosing a 30 year generation results in a 10% increase in the quoted mutation rate. As we find our mutation rates are close to the actual rates from meiosis this means the 27 year generation is also correct.

| # | DYS | Hamilton<sup>4</sup> | SD<sup>3</sup> | Burgella<sup>2</sup> | SD<sup>3</sup> | Chandler<sup>2</sup> | NIST<sup>2</sup> | FTDNA<sup>4</sup> |
| --- | --- | --- | --- | --- | --- | --- | --- | --- |
| 1. | 393 | 0.72 | 0.14 | 1.03 | 0.36 | 0.76 | 0.08 | 1.43 |
| 2. | 390 | 2.52 | 0.18 | 2.12 | 0.22 | 3.11 | 2.4 | 5.32 |
| 3. | 19/394<sup>1</sup> | 1.3 | 0.52 | 2.19 | 0.21 | 1.51 | 2.38 | 1.45 |
| 4. | 391 | 4.98 | 0.2 | 2.72 | 0.18 | 2.65 | 2.88 | 4.15 |
| 5. | 385a | 1.26 | 0.13 |  |  |  | 2.1 | 5.68 |
| 6. | 385b<sup>1</sup> | 3.13 | 0.34 |  |  |  | 2.1 | 5.68 |
| 7. | 426 | 0.07 | 0.24 |  |  | 0.09 |  | 0.26 |
| 8. | 388 | 0.22 | 0.22 | 0.42 | 2.31 | 0.22 |  | 0.25 |
| 9. | 439 | 3.76 | 0.11 | 5.48 | 0.16 | 4.77 |  | 4.95 |
| 10. | 389-I | 1.93 | 0.1 | 2.53 | 0.21 | 1.86 | 1.88 | 2.23 |
| 11. | 392 | 0.36 | 0.27 | 0.43 | 0.59 | 0.52 | 0.58 | 1.59 |
| 12. | 389b | 2.96 | 0.11 | 3.17 | 0.18 | 2.42 | 2.96 | 2.72 |
| 13. | 458 | 7.99 | 0.08 | 6.88 | 0.16 | 8.14 | 10.8 | 6.3 |
| 14. | 459a | 0.39 | 0.18 |  |  |  |  |  |
| 15. | 459b<sup>1</sup> | 2.98 | 0.47 |  |  |  |  |  |
| 16. | 455 | 0.16 | 0.21 |  |  | 0.16 |  | 0.46 |
| 17. | 454 | 0.11 | 0.22 |  |  | 0.16 |  | 0.47 |
| 18. | 447 | 3.8 | 0.15 | 4.56 | 0.96 | 2.64 |  | 4. |
| 19. | 437 | 0.99 | 0.18 |  |  | 0.99 | 1.5 | 2.15 |
| 20. | 448 | 1.16 | 0.21 |  |  | 1.35 | 1.8 | 2.71 |
| 21. | 449 | 11.7 | 0.14 | 18.97 | 0.52 | 8.38 |  | 7.84 |
| 22. | 460 | 2.63 | 0.13 | 3.82 | 0.66 | 4.02 |  |  |
| 23. | GATAH4 | 3.93 | 0.1 | 2.24 | 0.44 | 2.08 | 2.51 |  |
| 24. | YCA IIa | 0.32 | 0.23 |  |  |  |  |  |
| 25. | YCA IIb | 1.4 | 0.18 |  |  |  |  |  |
| 26. | 456 | 8.1 | 0.23 | 4.5 | 0.21 | 7.35 |  |  |
| 27. | 607 | 2.15 | 0.13 |  |  | 4.11 |  | 4.1 |
| 28. | 576 | 10.65 | 0.11 | 16.22 | 0.44 | 10.22 |  | 10.2 |
| 29. | 570 | 4.6 | 0.2 | 12.61 | 0.52 | 7.9 |  | 7.9 |
| 30. | CDYa | 14.71 | 0.09 |  |  |  |  | 35.3 |
| 31. | CDYb<sup>1</sup> | 13.4 | 2. |  |  |  |  | 35.3 |
| 32. | 442 | 2.9 | 0.11 |  |  | 3.24 |  |  |
| 33. | 438 | 0.43 | 0.14 |  |  | 0.55 | 0.7 |  |
| Mean |  | 3.6 |  | 5.3 |  | 3.2 | 2.5 | 6.4 |

| # | DYS | A<sub>j</sub> | SD | SD to 0.5 |
| --- | --- | --- | --- | --- |
| 1. | 393 | 0.675 | 0.087 | 2. |
| 2. | 390 | 0.463 | 0.093 | 0.4 |
| 3. | 19/394 | 0.973 | 0.032 | 14.7 |
| 4. | 391 | 0.029 | 0.008 | 62.8 |
| 5. | 385a | 0.699 | 0.096 | 2.1 |
| 6. | 385b | 0.82 | 0.085 | 3.8 |
| 7. | 426 | 0.37 | 0.232 | 0.6 |
| 8. | 388 | 0.91 | 0.072 | 5.7 |
| 9. | 439 | 0.734 | 0.359 | 0.7 |
| 10. | 389-I | 0.779 | 0.105 | 2.7 |
| 11. | 392 | 0.954 | 0.04 | 11.2 |
| 12. | 389b | 0.703 | 0.325 | 0.6 |
| 13. | 458 | 0.512 | 0.137 | 0.1 |
| 14. | 459a | 0.139 | 0.125 | 2.9 |
| 15. | 459b | 0.003 | 0.001 | 353. |
| 16. | 455 | 0.277 | 0.168 | 1.3 |
| 17. | 454 | 0.962 | 0.03 | 15.4 |
| 18. | 447 | 0.154 | 0.025 | 13.6 |
| 19. | 437 | 0.09 | 0.09 | 4.6 |
| 20. | 448 | 0.216 | 0.172 | 1.6 |
| 21. | 449 | 0.518 | 0.15 | 0.1 |
| 22. | 460 | 0.107 | 0.05 | 7.9 |
| 23. | GATAH4 | 0.17 | 0.198 | 1.7 |
| 24. | YCAIIa | 0.195 | 0.163 | 1.9 |
| 25. | YCAIIb | 0.19 | 0.175 | 1.8 |
| 26. | 456 | 0.671 | 0.416 | 0.4 |
| 27. | 607 | 0.243 | 0.103 | 2.5 |
| 28. | 576 | 0.387 | 0.157 | 0.7 |
| 29. | 570 | 0.448 | 0.077 | 0.7 |
| 30. | CDYa | 0.37 | 0.181 | 0.7 |
| 31. | CDYb | 0.258 | 0.082 | 2.9 |
| 32. | 442 | 0.603 | 0.17 | 0.6 |
| 33. | 438 | 0.715 | 0.215 | 1. |
| Mean* | #1-33 | 0.26* | 0.134 |  |

## Complete worked example for G2a3, R1b1a2, R1a1, I1, L21, U106, J2, P312

We use 29 markers (standard method) for G2a3, R1b1a2,R1a1, I1, L21, U106, J2, P312, requires running compiled functions from 29ComFun and its data file W29ComFun First we enter DNA file δδ

~~~
*δδ;*
~~~

Each file has NN members

~~~
NN = Table[Length [*δδ*[[q1]]], {q1, 1, 8}]
~~~

~~~
{1221, 460, 1270, 2898, 1029, 1533, 1241, 971}
~~~

We use asymptotic rates *α*0, *β*0,LB shown

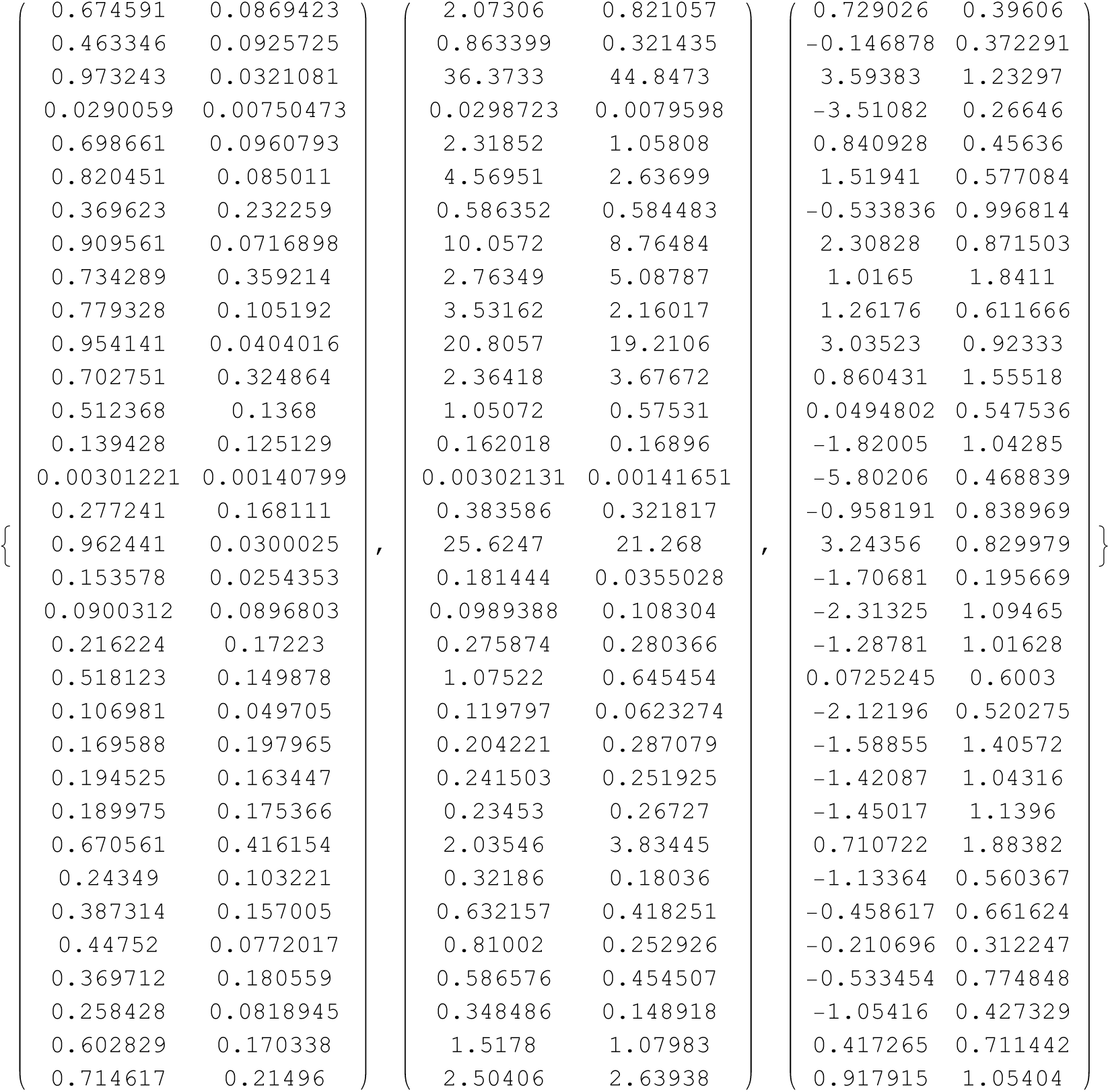

~~~
Table[ 1, {j, 1, 33}]; BB[j_] := B[[j]]; BB0 = Table[ 1, {j, 1, 33}];
~~~

~~~
PP = Table[ { N[C]F[Normal]istribution[0, 1], -4 + 0.01*j]], -4 + 0.01*j}, {j, 0, 800}];
~~~

~~~
P = Interpolation[PP];
~~~

We bootstrap with n02 cycles

~~~
n02 = 1000
~~~

~~~
500
~~~

The method of reduction is applied

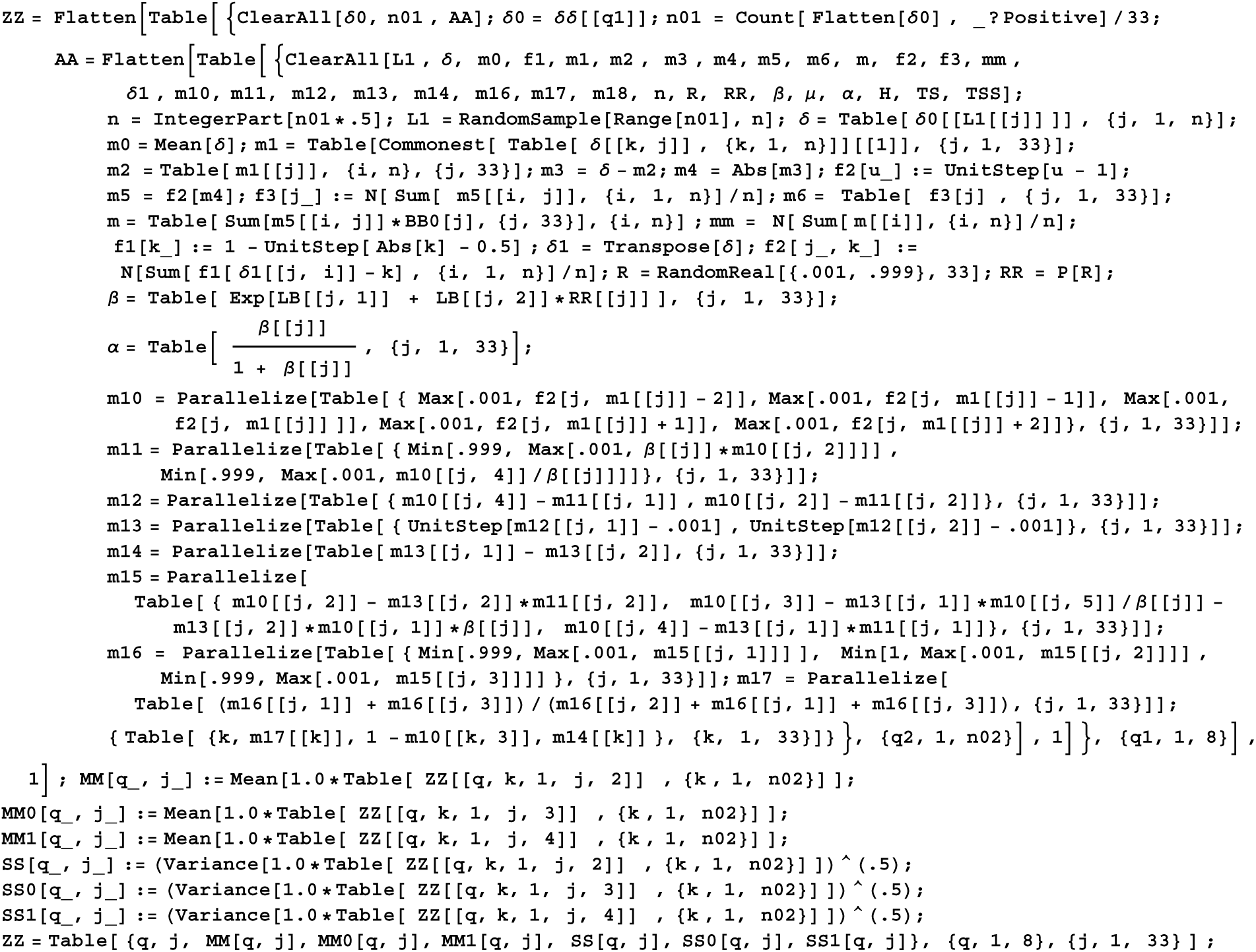

The output is for each file (q), marker (j), reduced frequency f0, unreduced frequency f0, mean ± then SD for each

~~~
MatrixForm[Transpose[ZZ]]
~~~

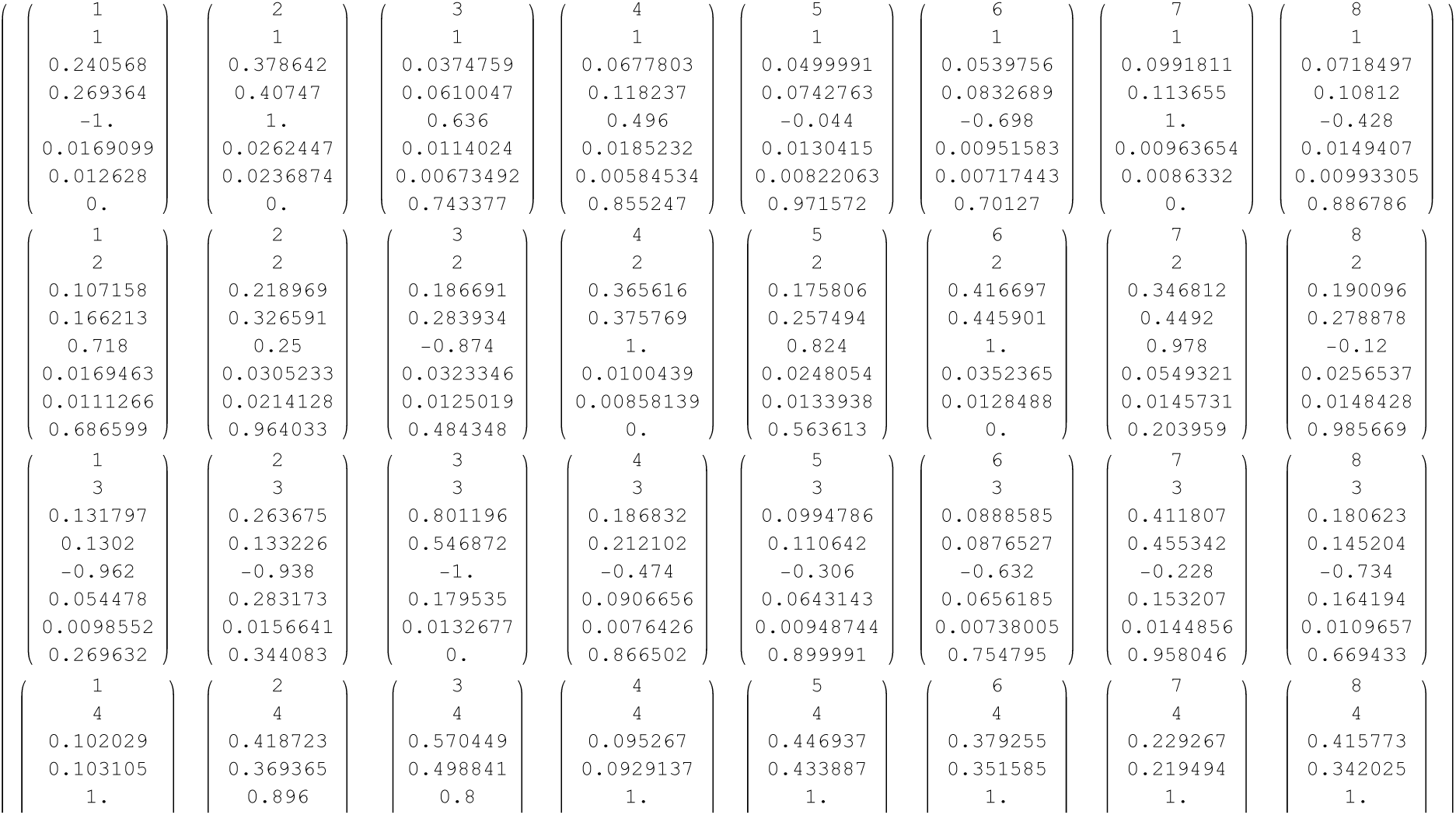
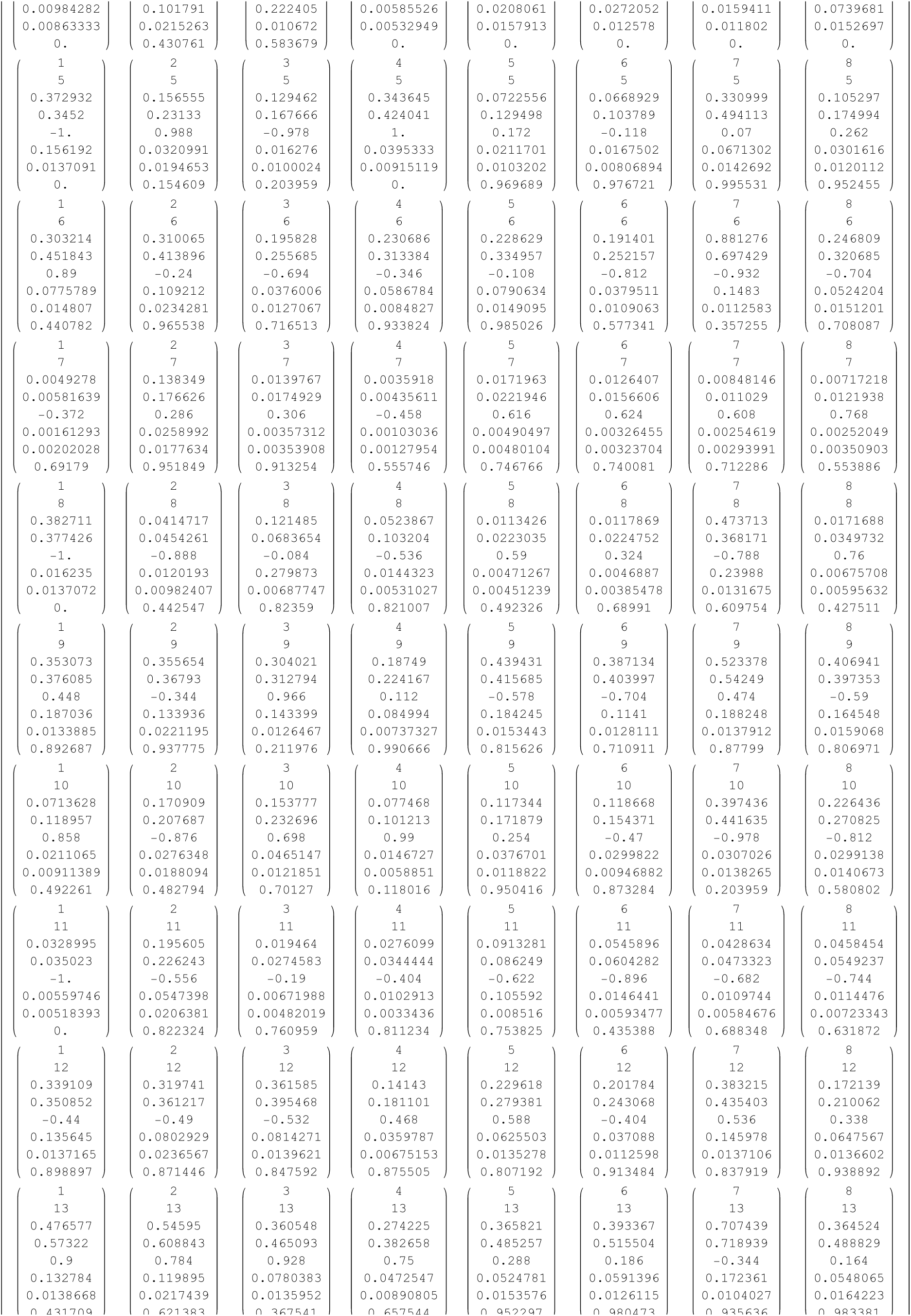
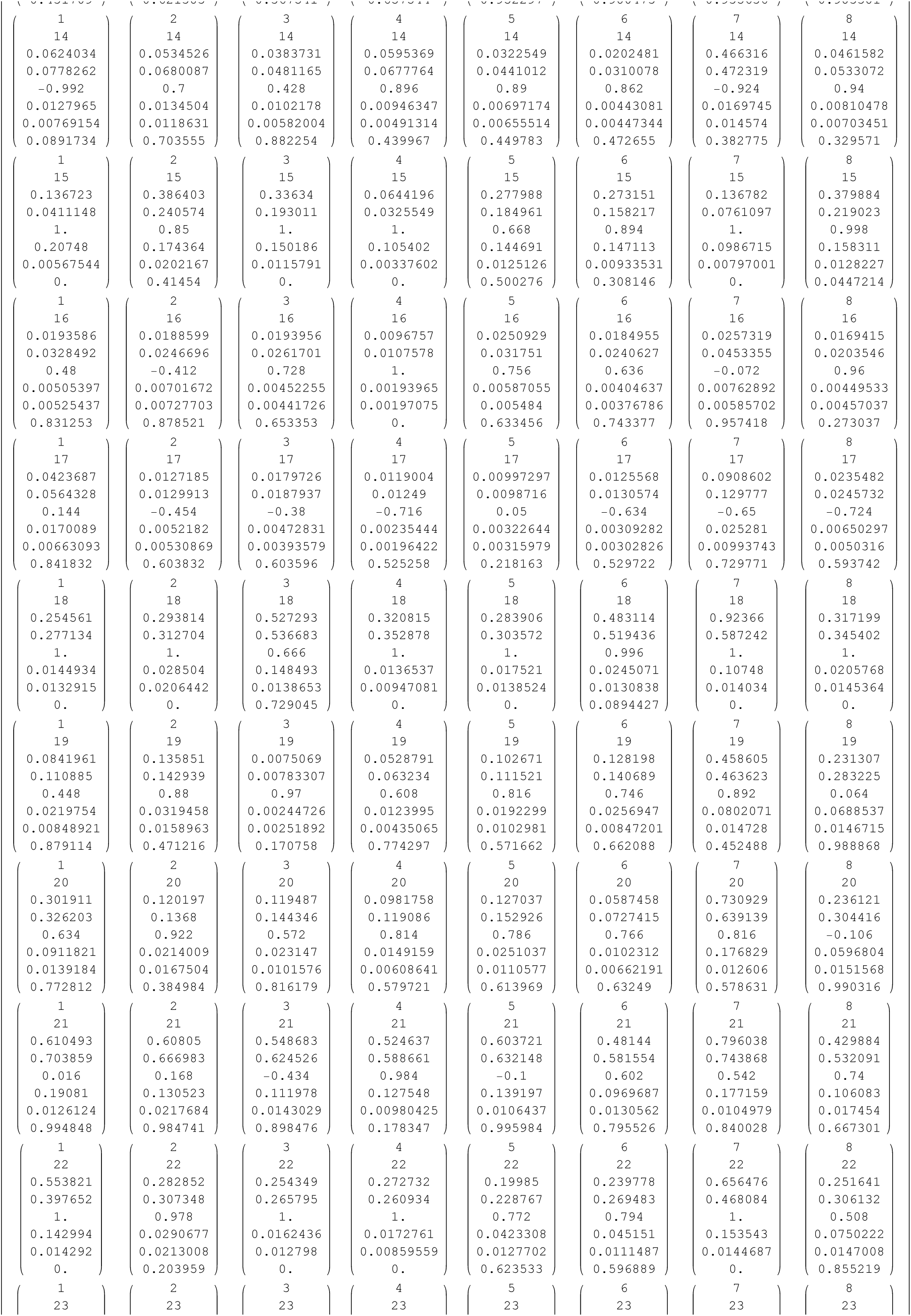
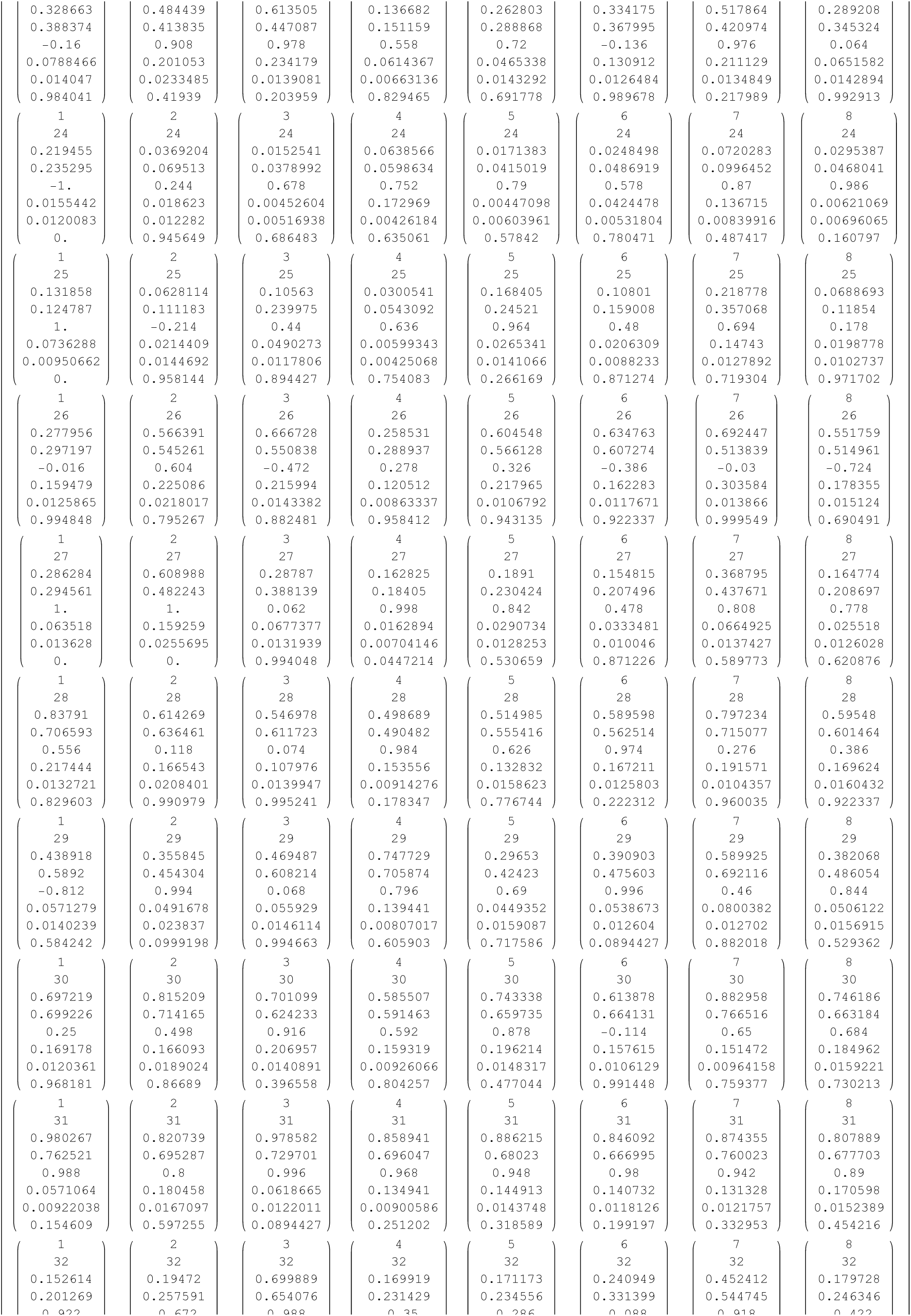
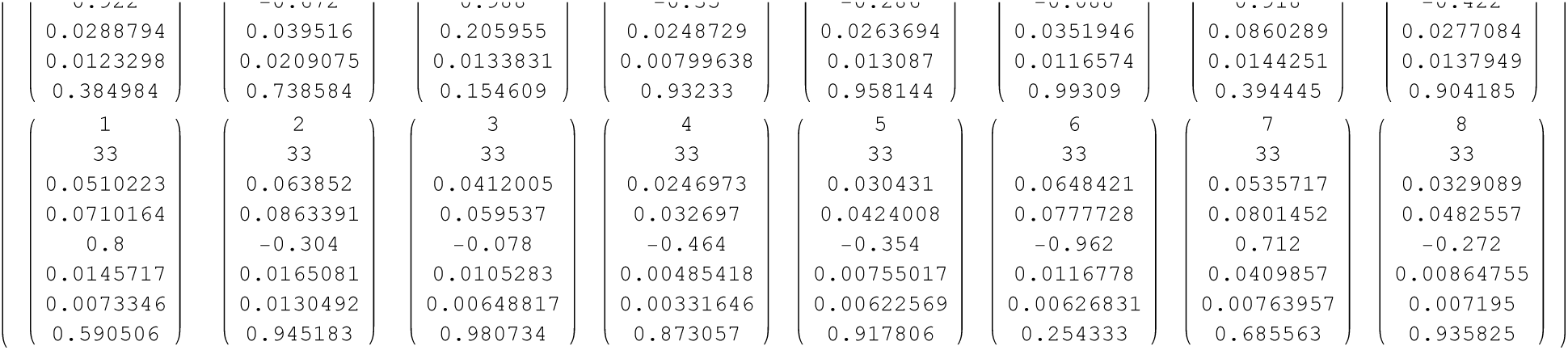

We conservatively weight the reduction by mean ±

~~~
ZZZ = Table[{q1, k, Abs[ZZ[[q1, k, 5]]] * ZZ[[q1, k, 3]] + (1 - Abs[ZZ[[q1, k, 5]]]) * ZZ[[q1, k, 4]] }, {q1, 1, 8}, {k, 1, 33}];
~~~

~~~
MatrixForm[Transpose[ZZZ]]
~~~

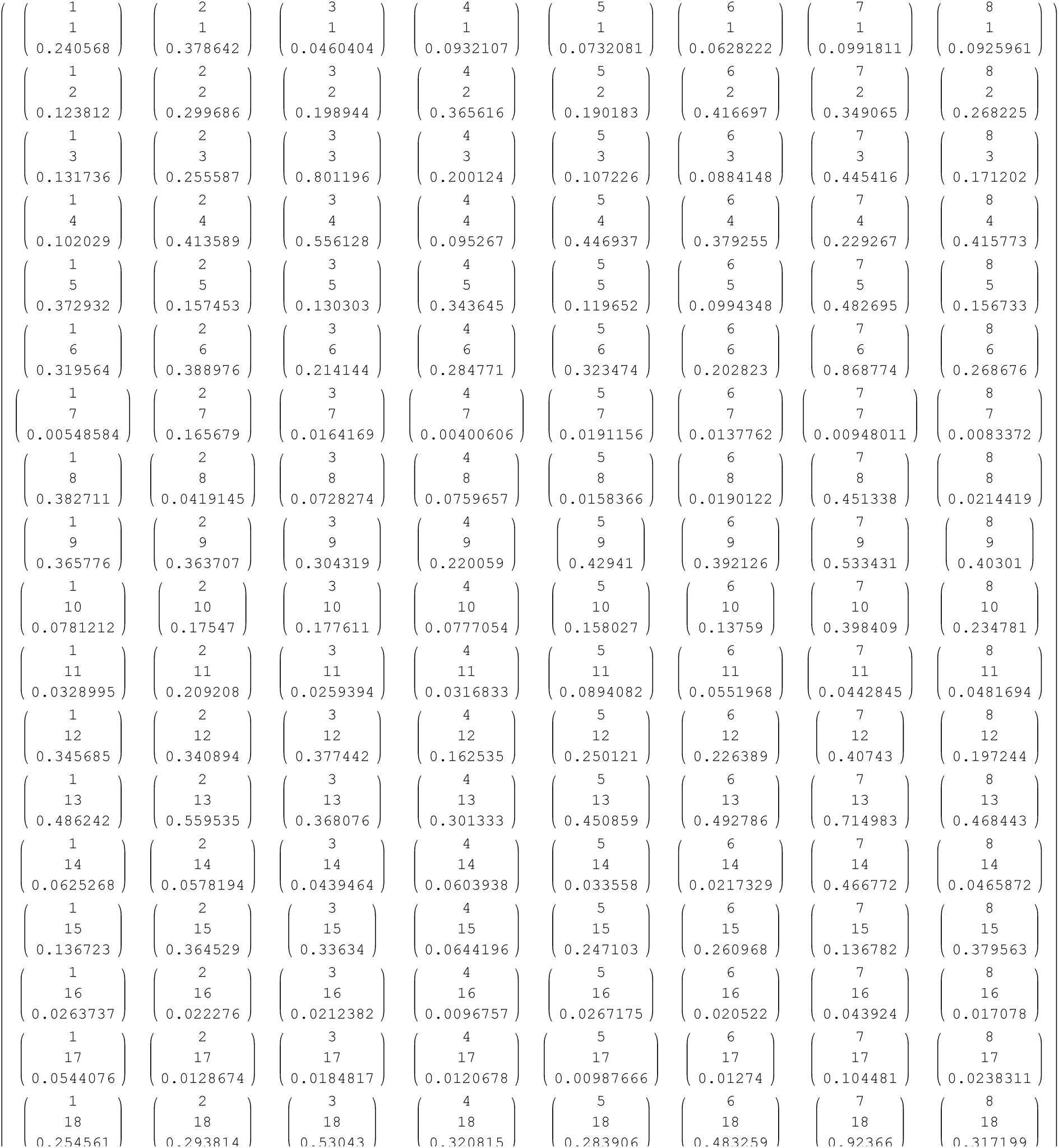
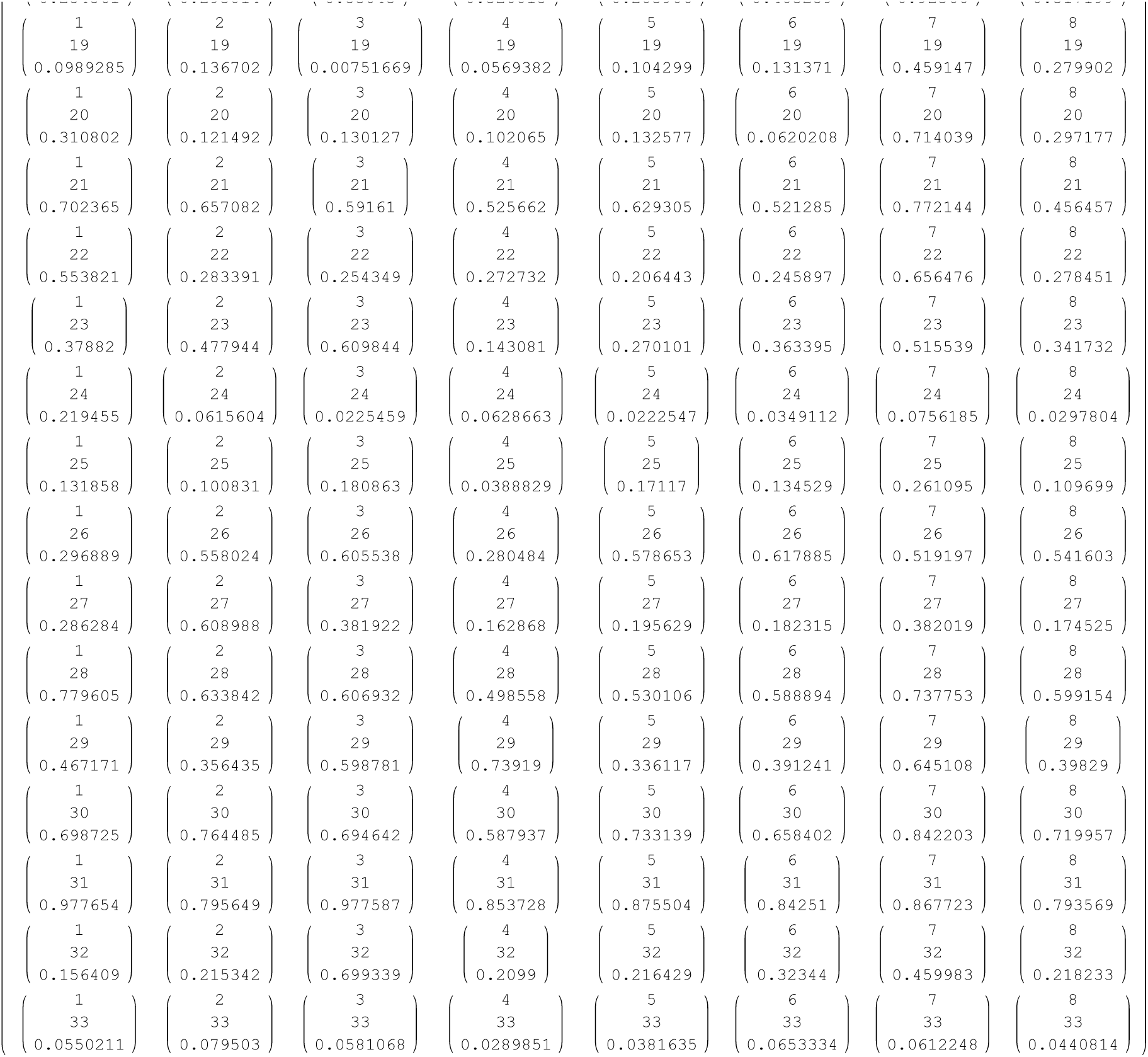

This has SD

~~~
ZZS = Table[(((Abs[ZZ[[q1, k, 5]]]*(ZZ[[q1, k, 6]] ^2 + ZZ[[q1, k, 3]] ^2) + (1 - Abs[ZZ[[q1, k, 5]]])*(ZZ[[q1, k, 7]] ^2 + ZZ[[q1, k, 4]] ^2) Abs[ZZ[[q1, k, 5]]] ZZ[[q1, k, 3]] + (1 - Abs[ZZ[[q1, k, 5]]]) ZZ[[q1, k, 4]]) ^ 2)) ^ (.5)), {q1, 1, 8}, {k, 1, 33}]
~~~

~~~
MatrixForm[ Transpose[ZZS]]
~~~

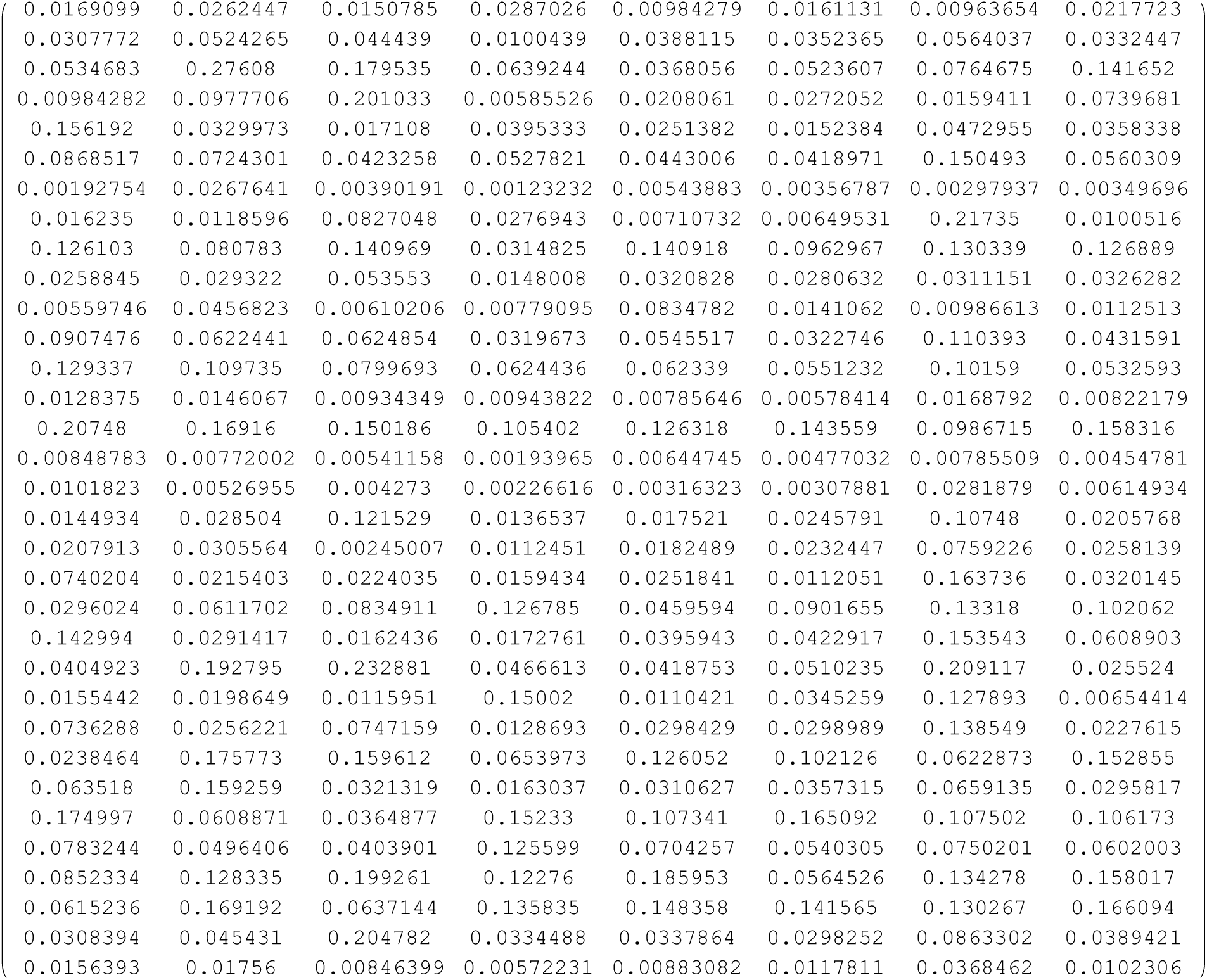

To computes times for each marker need mutation rates

~~~
MatrixForm[ μ00]
~~~

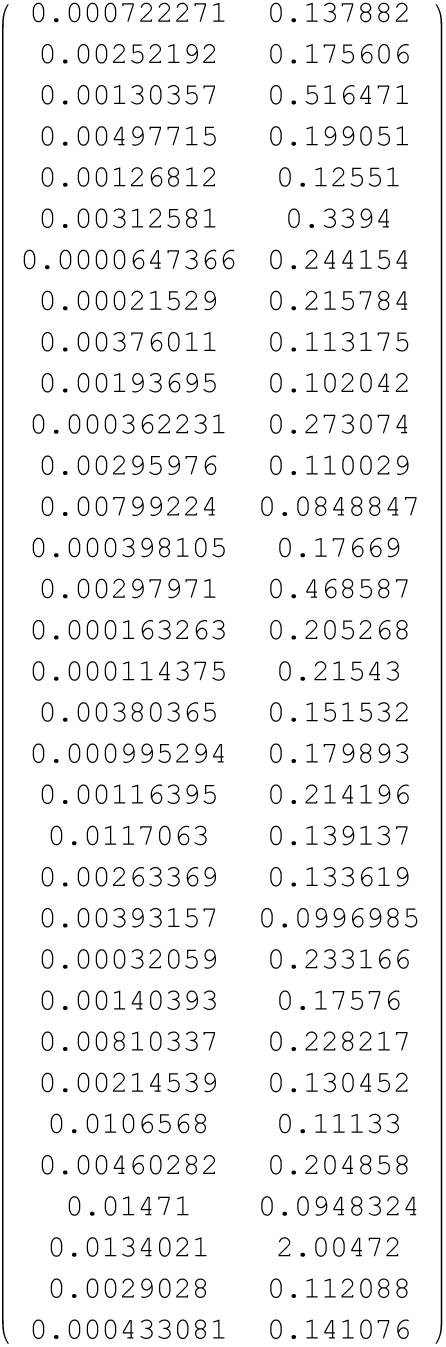

Also need hyperbolic Bessel functions

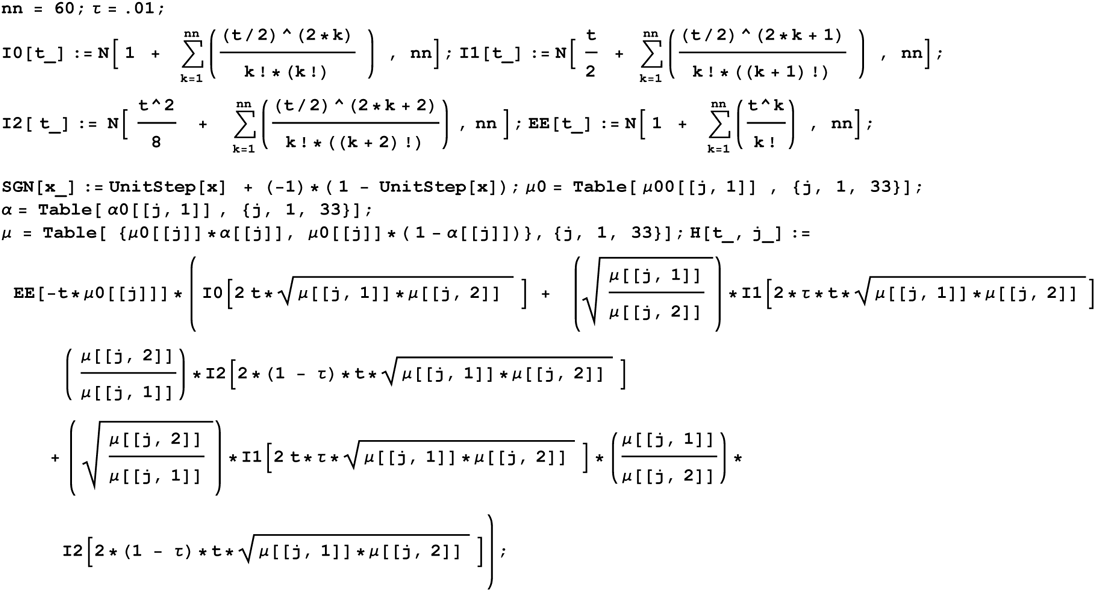

The times T for each marker are obtained by

~~~
TS[k_] := Normal[InverseSeries[Series[ (1 - H[x, k]), {x, 0, 60}] ]]; TSS[t_, k_] := TS[ k] /. x → t ;
~~~

which is applied to give expansion times for each marker

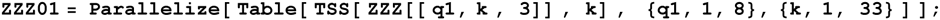

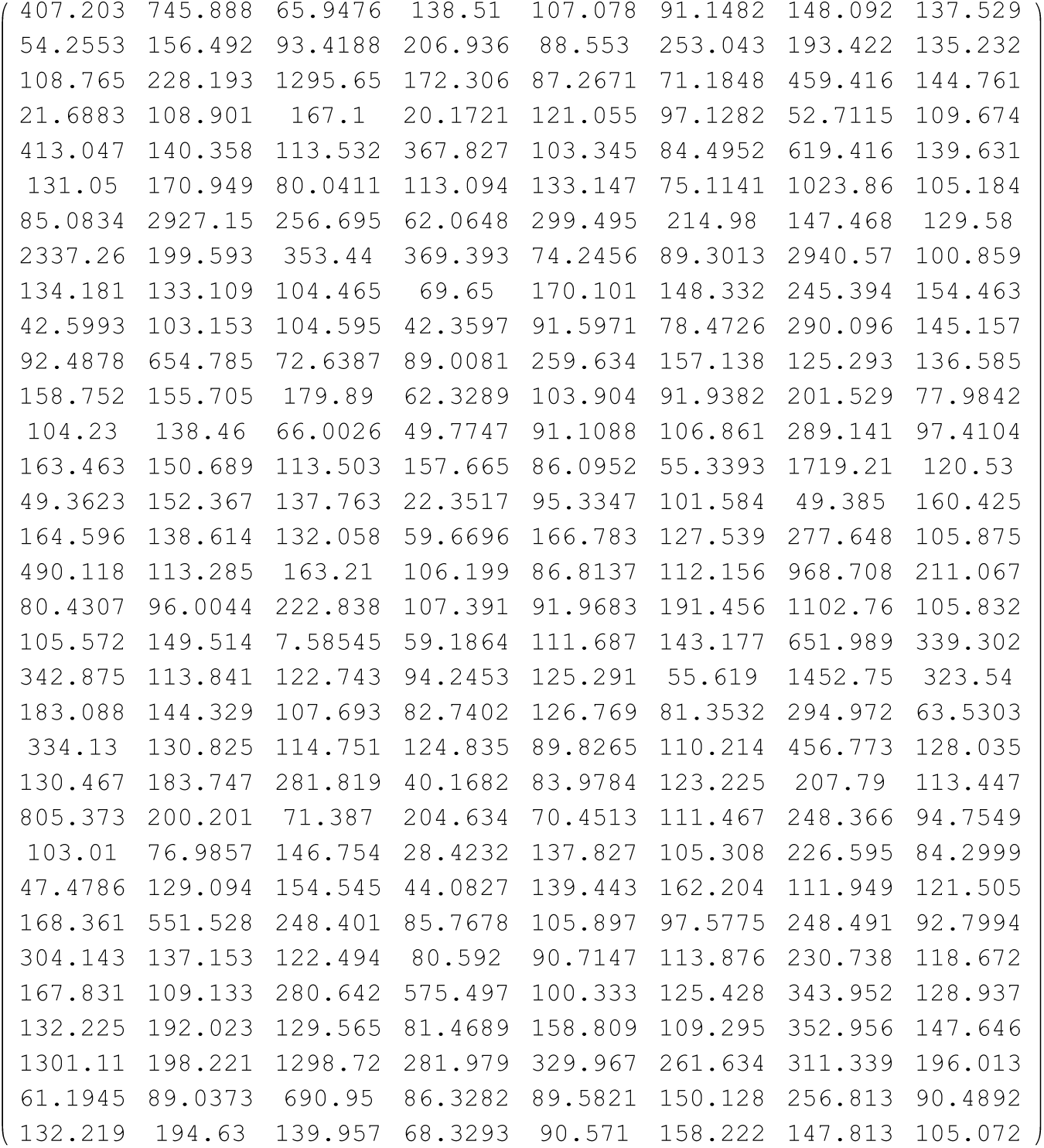

The SD for T are

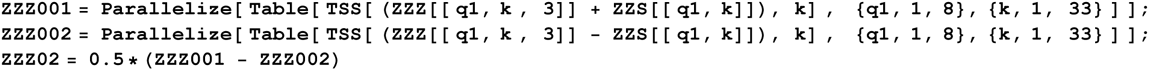

~~~
MatrixForm[ Transpose[ZZZ02]]
~~~

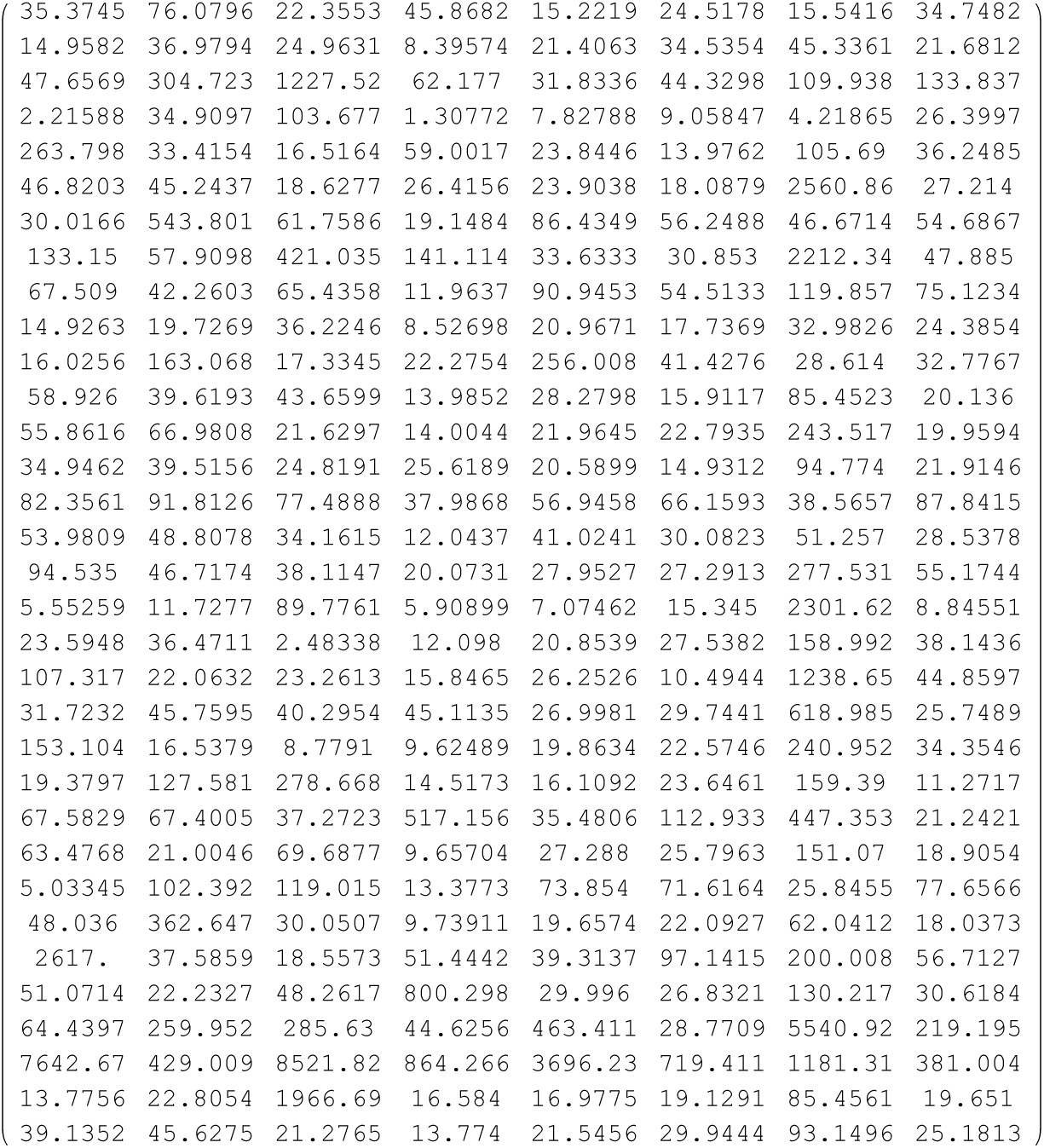

We are only using 29 markers

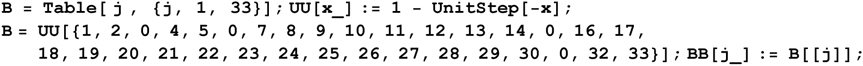

We first sketch the spectrum of times

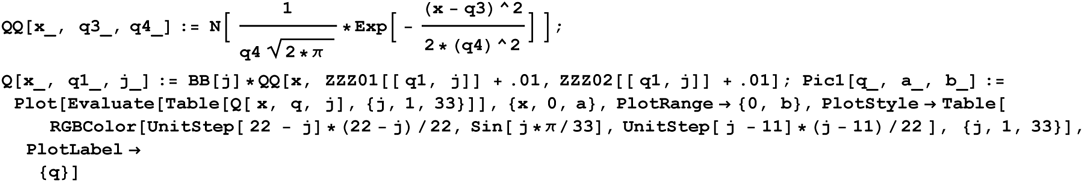

This plots the time distributions for each marker

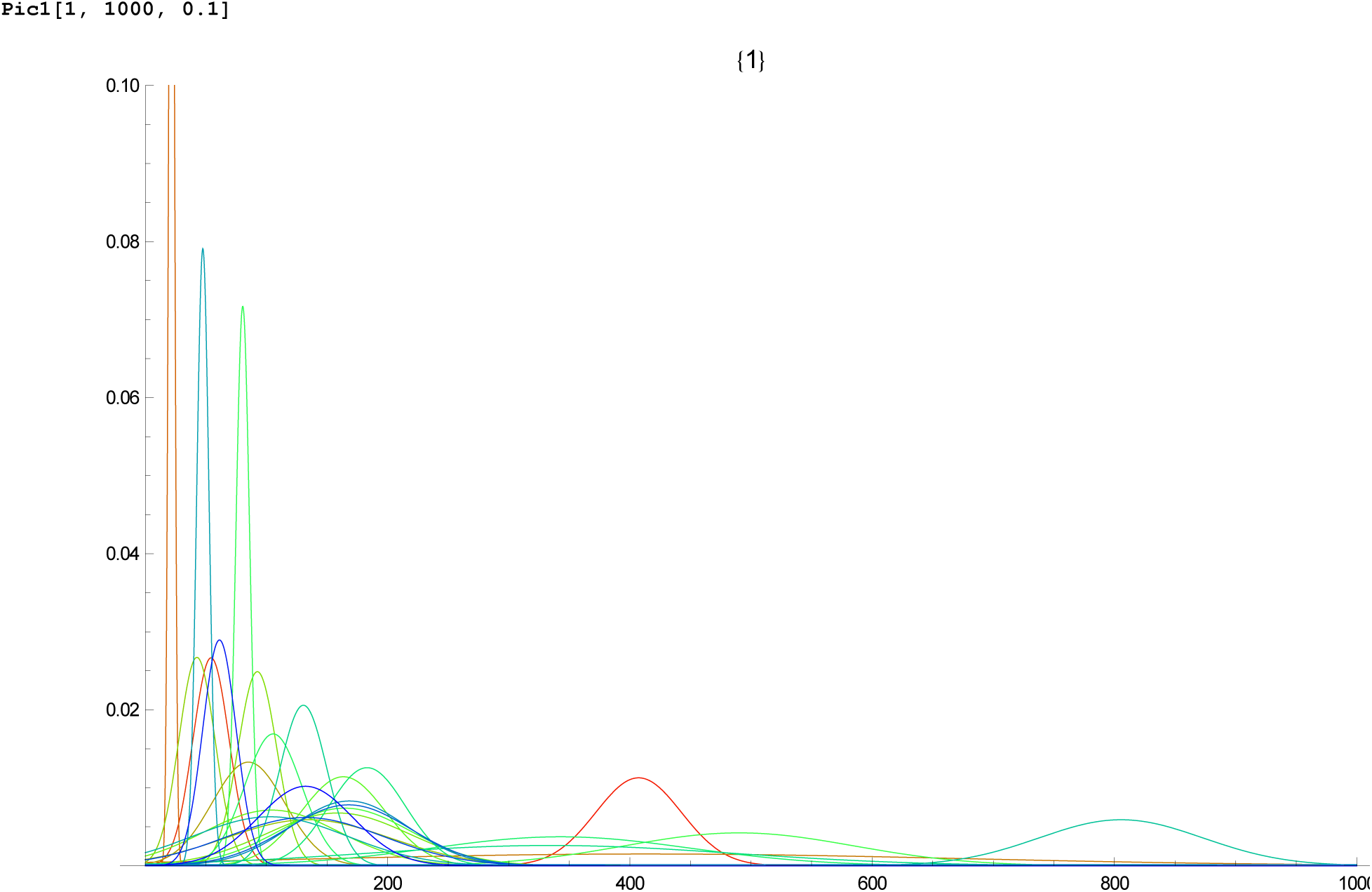

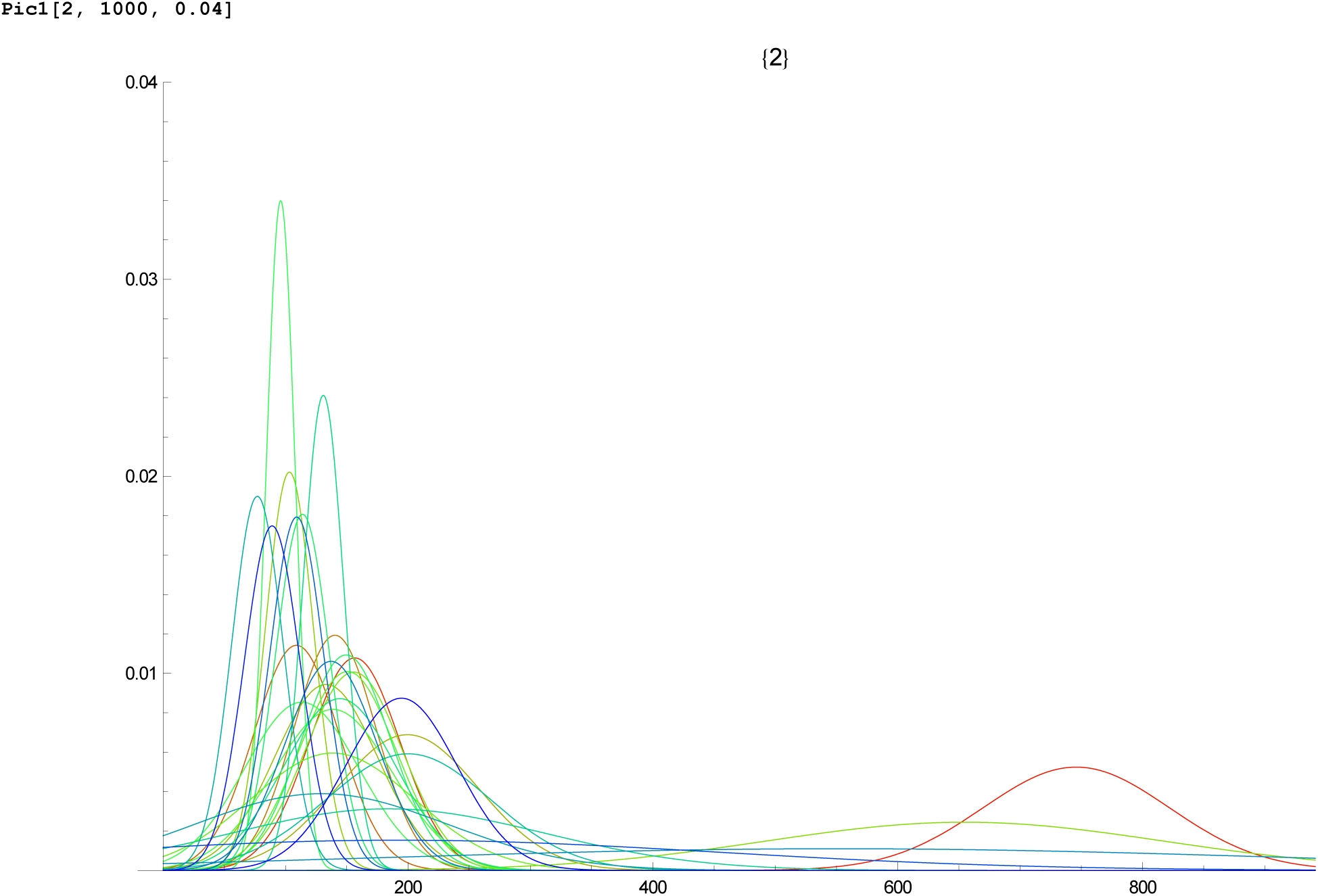

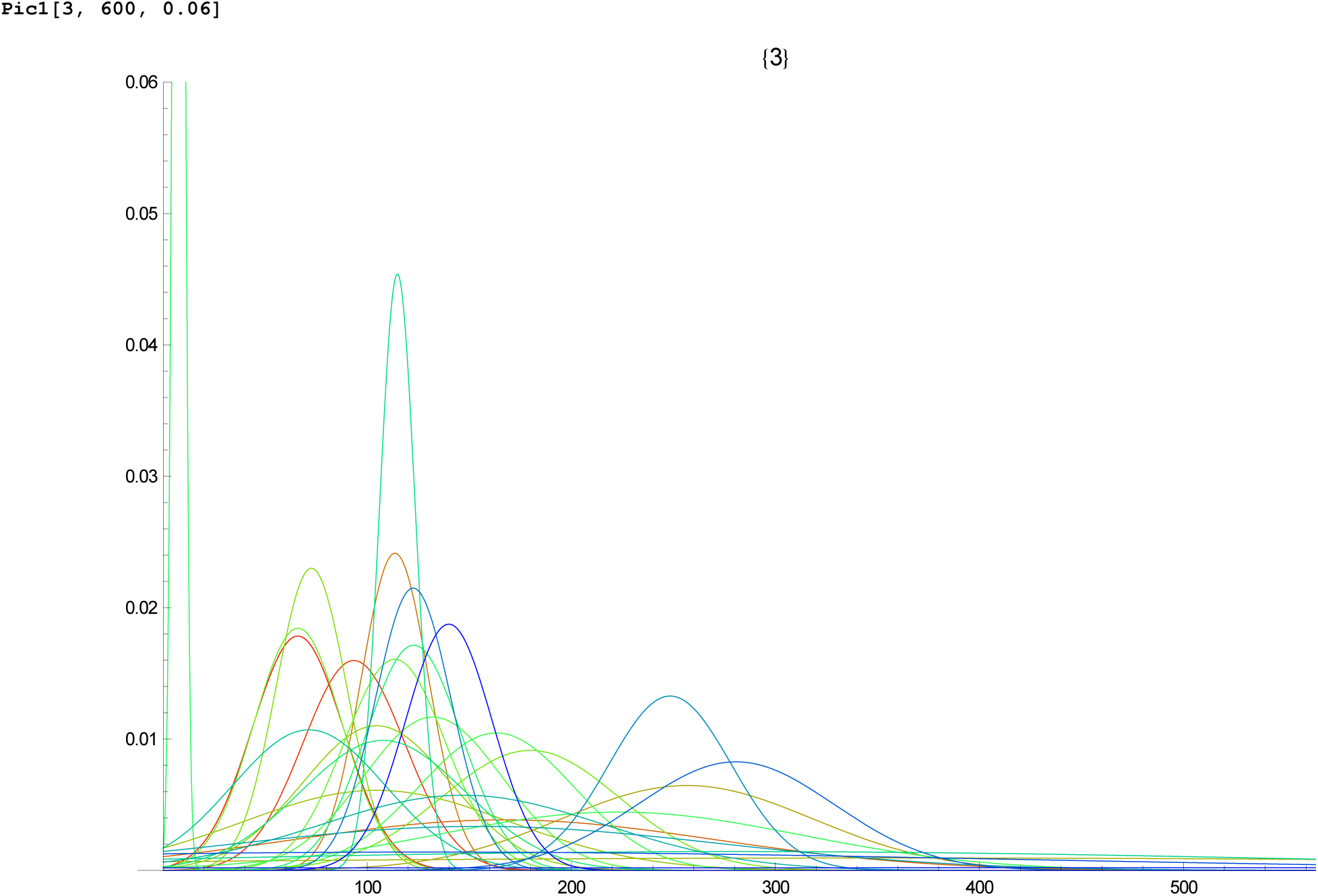

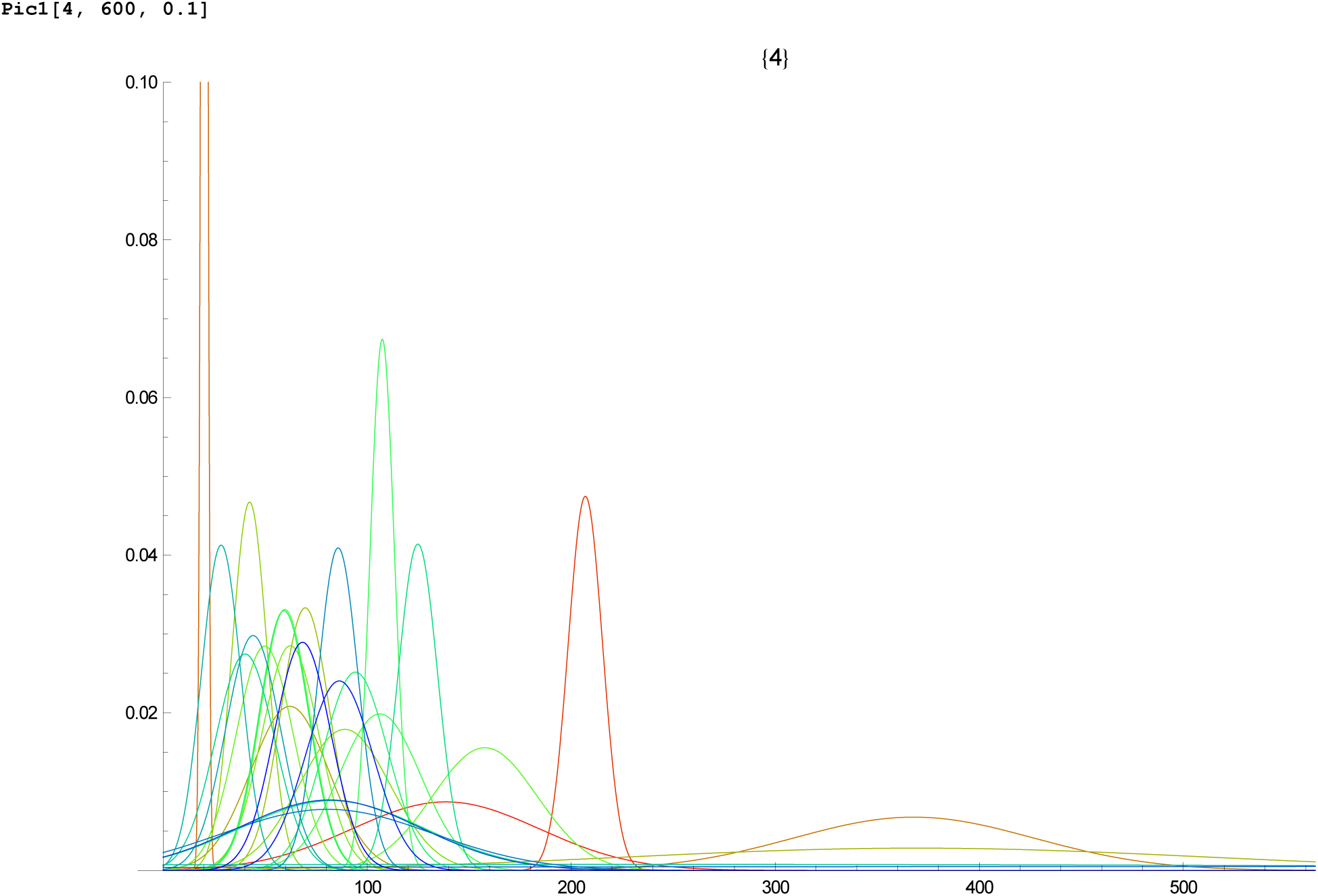

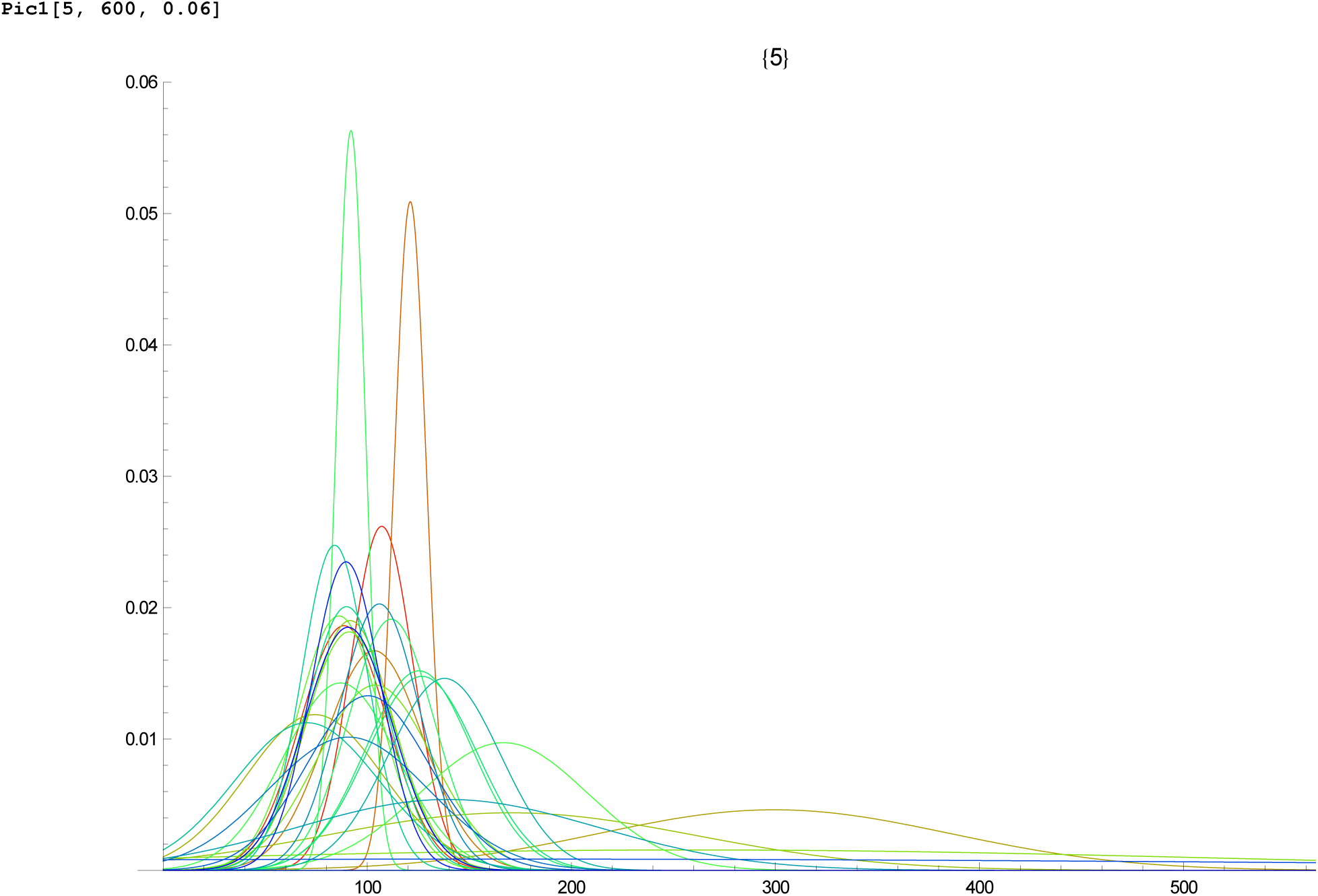

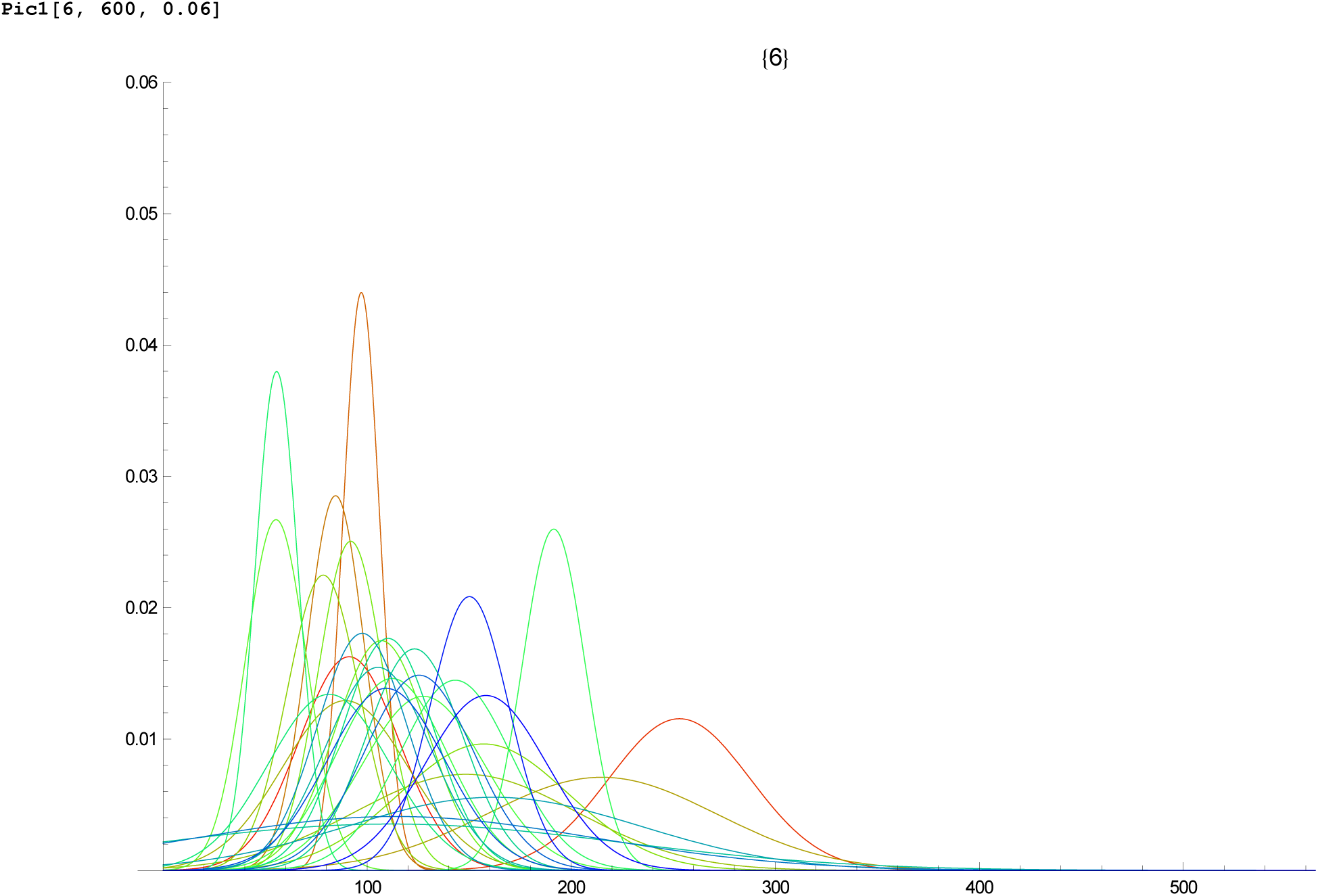

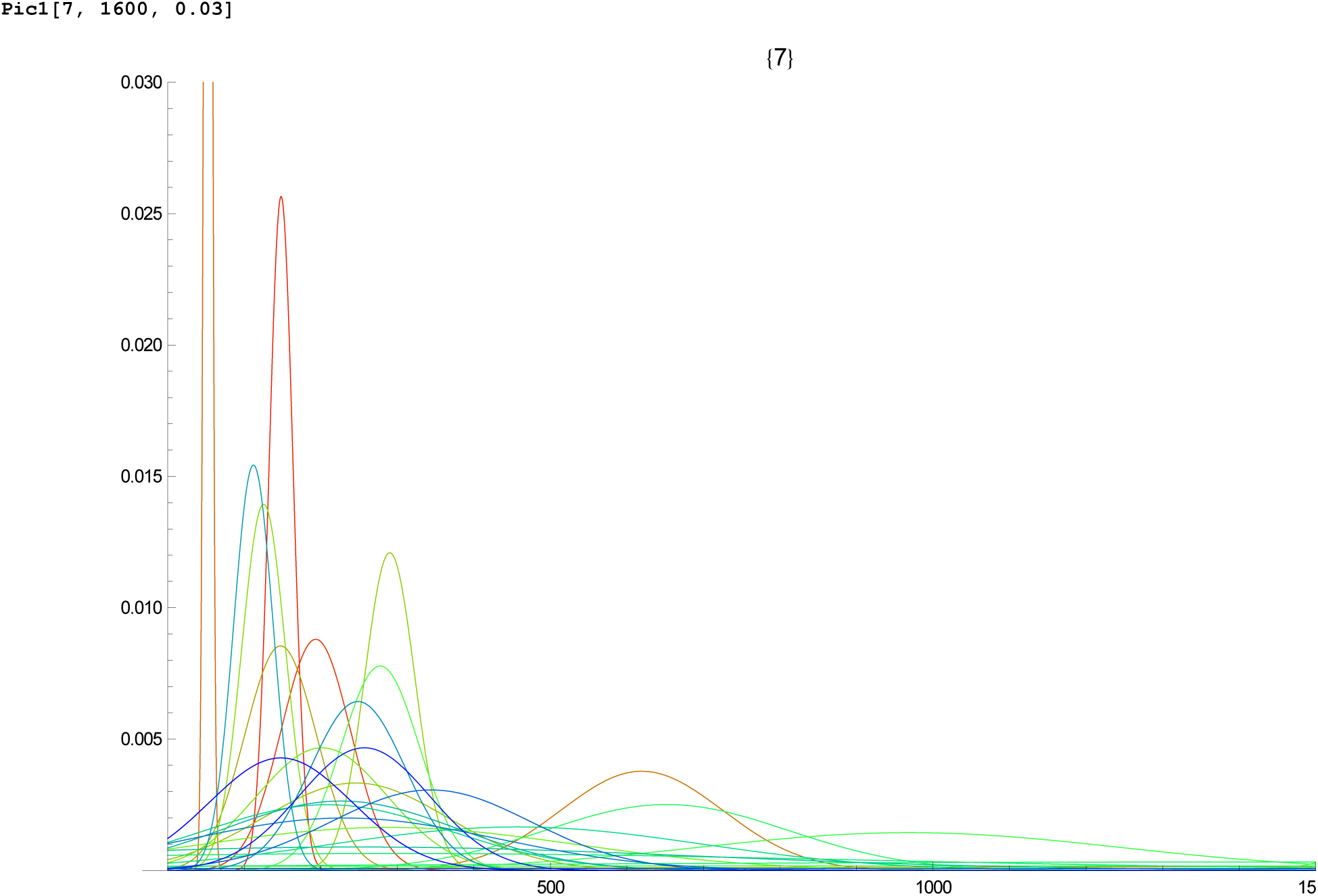

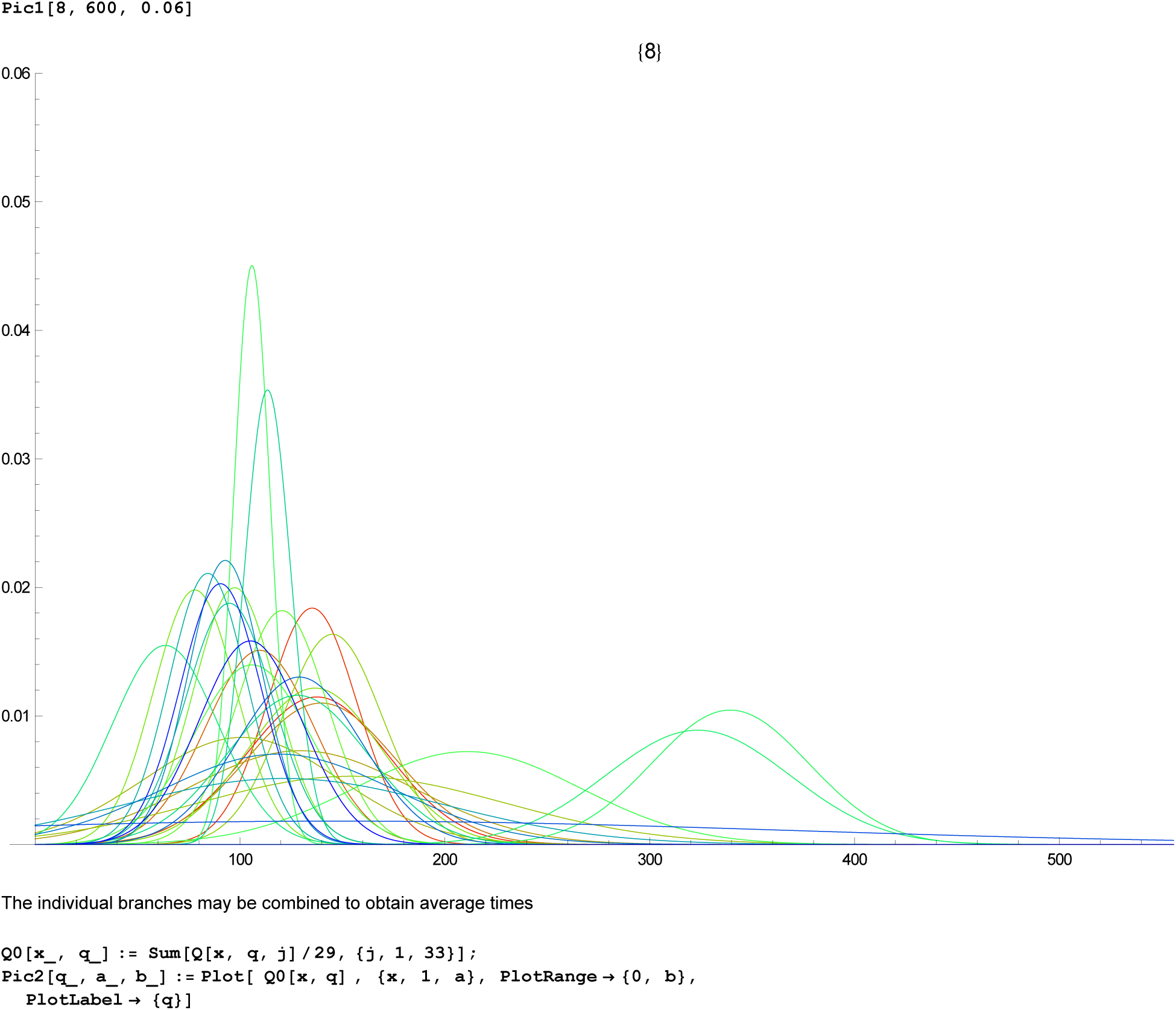

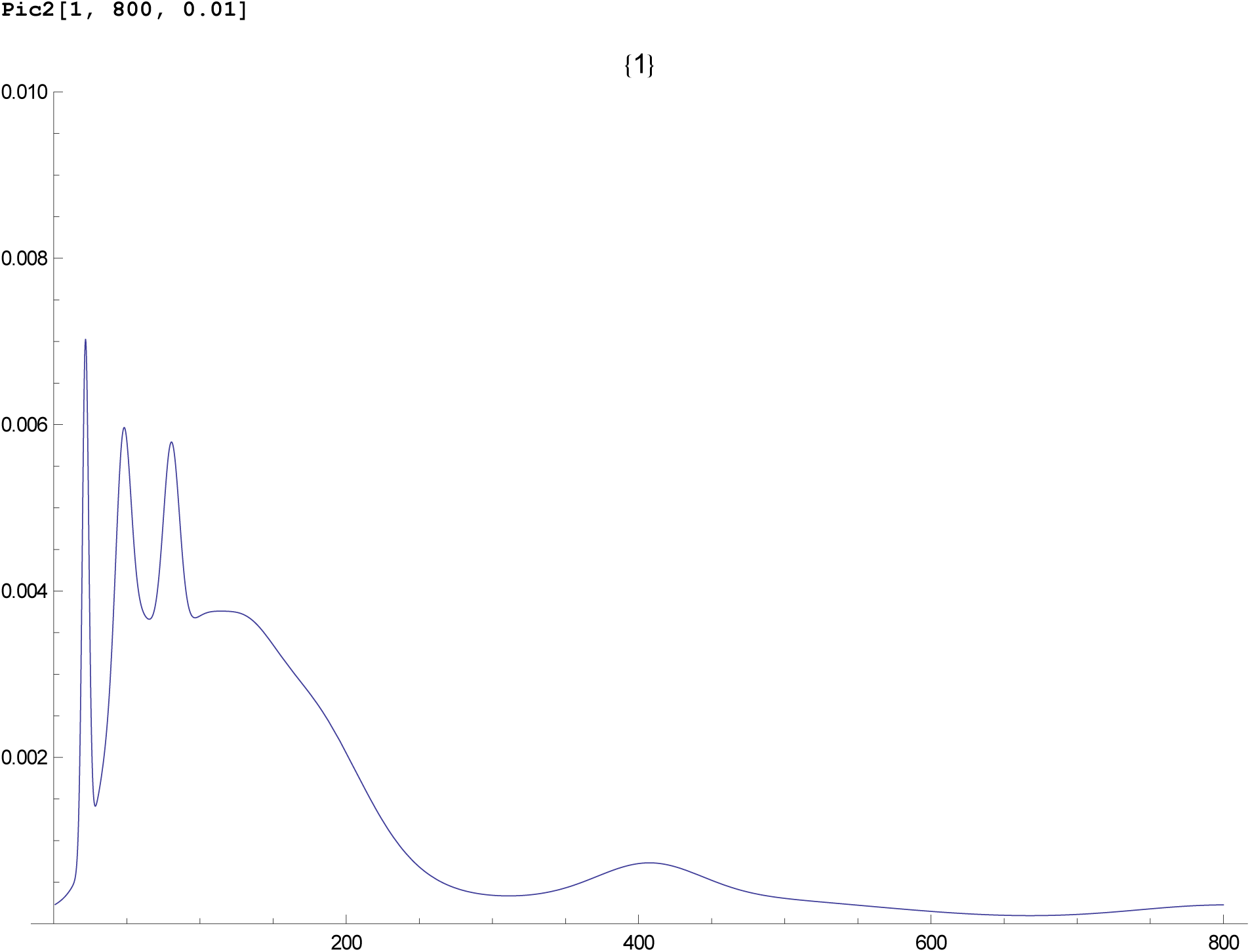

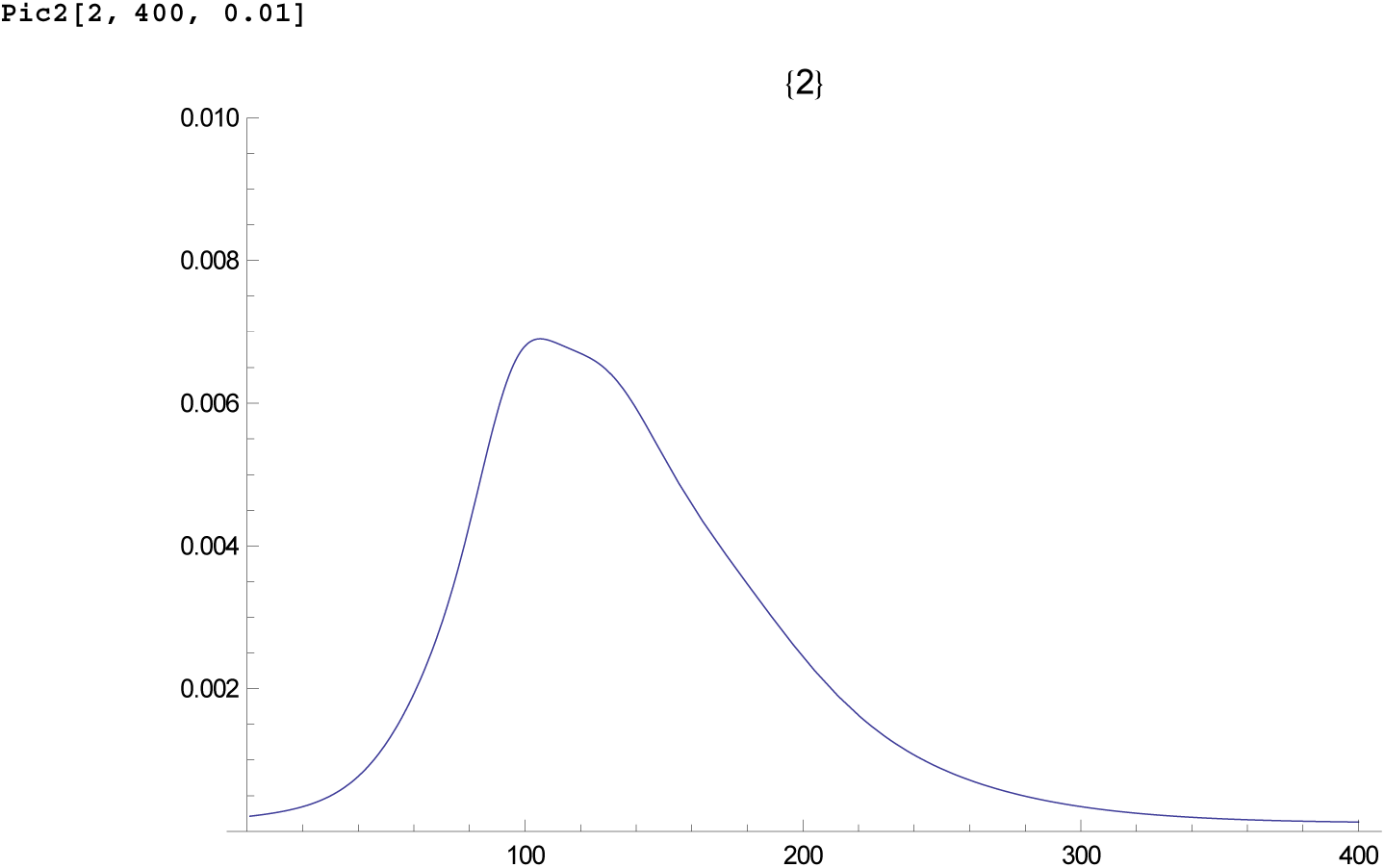

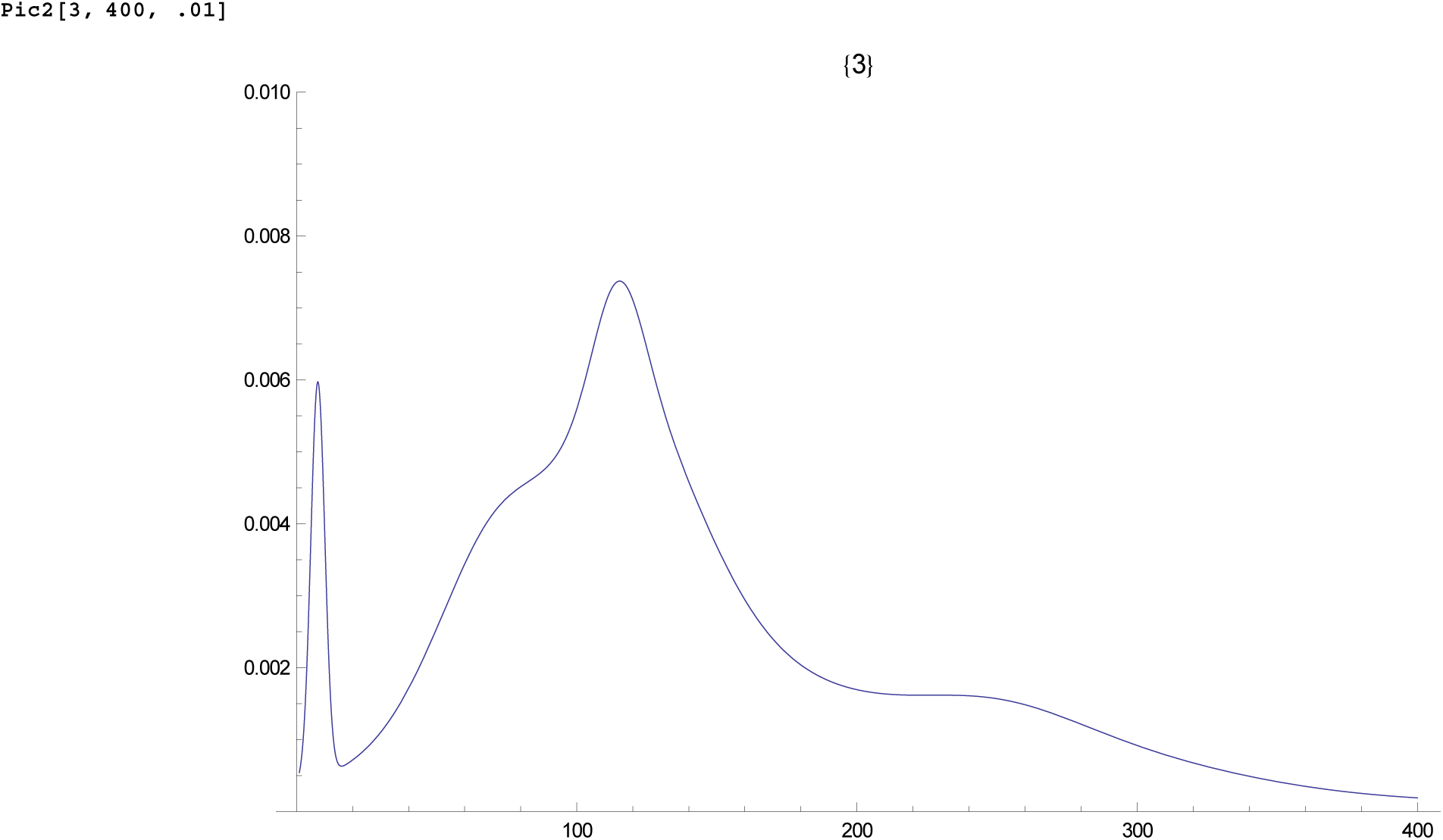

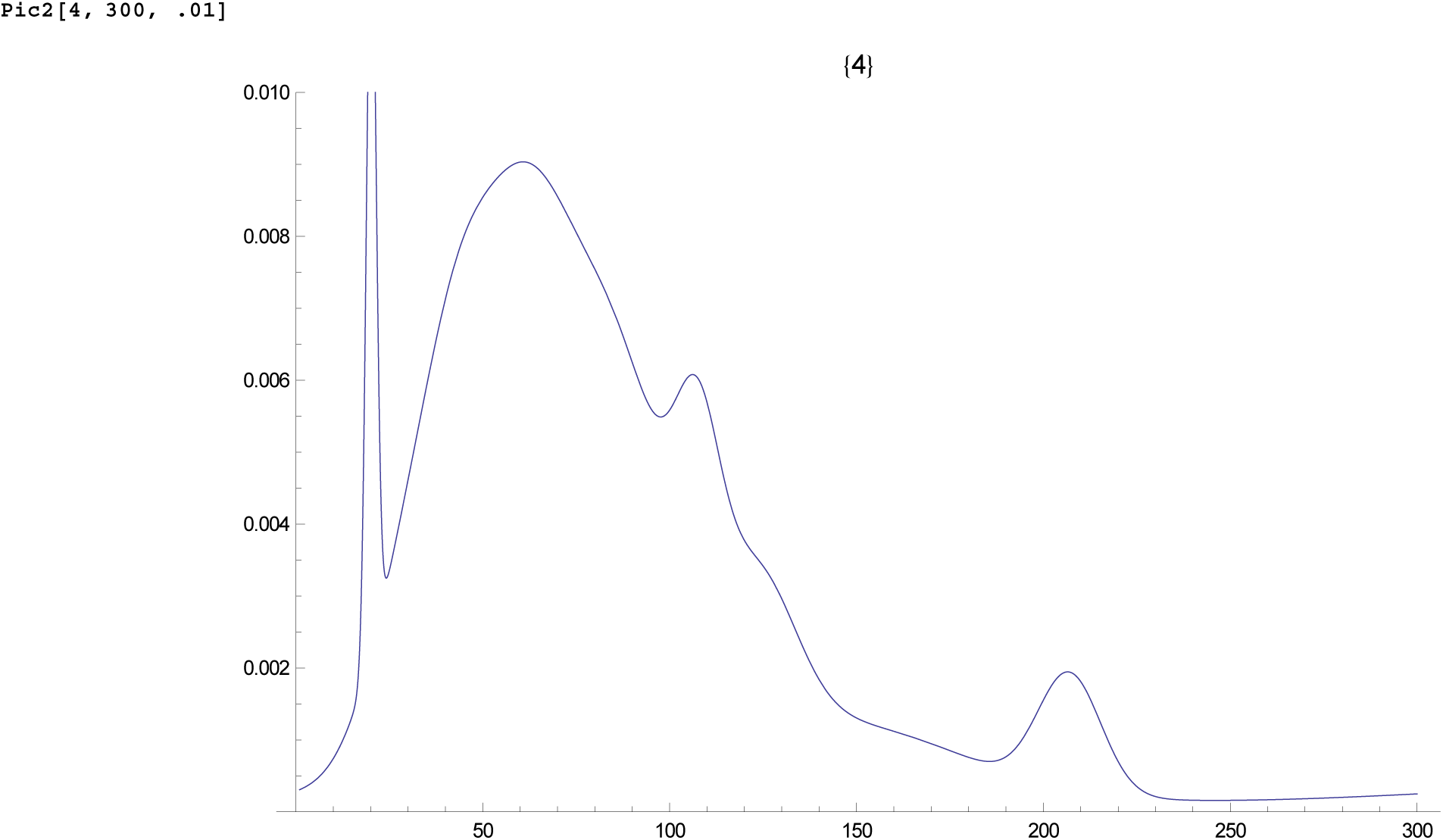

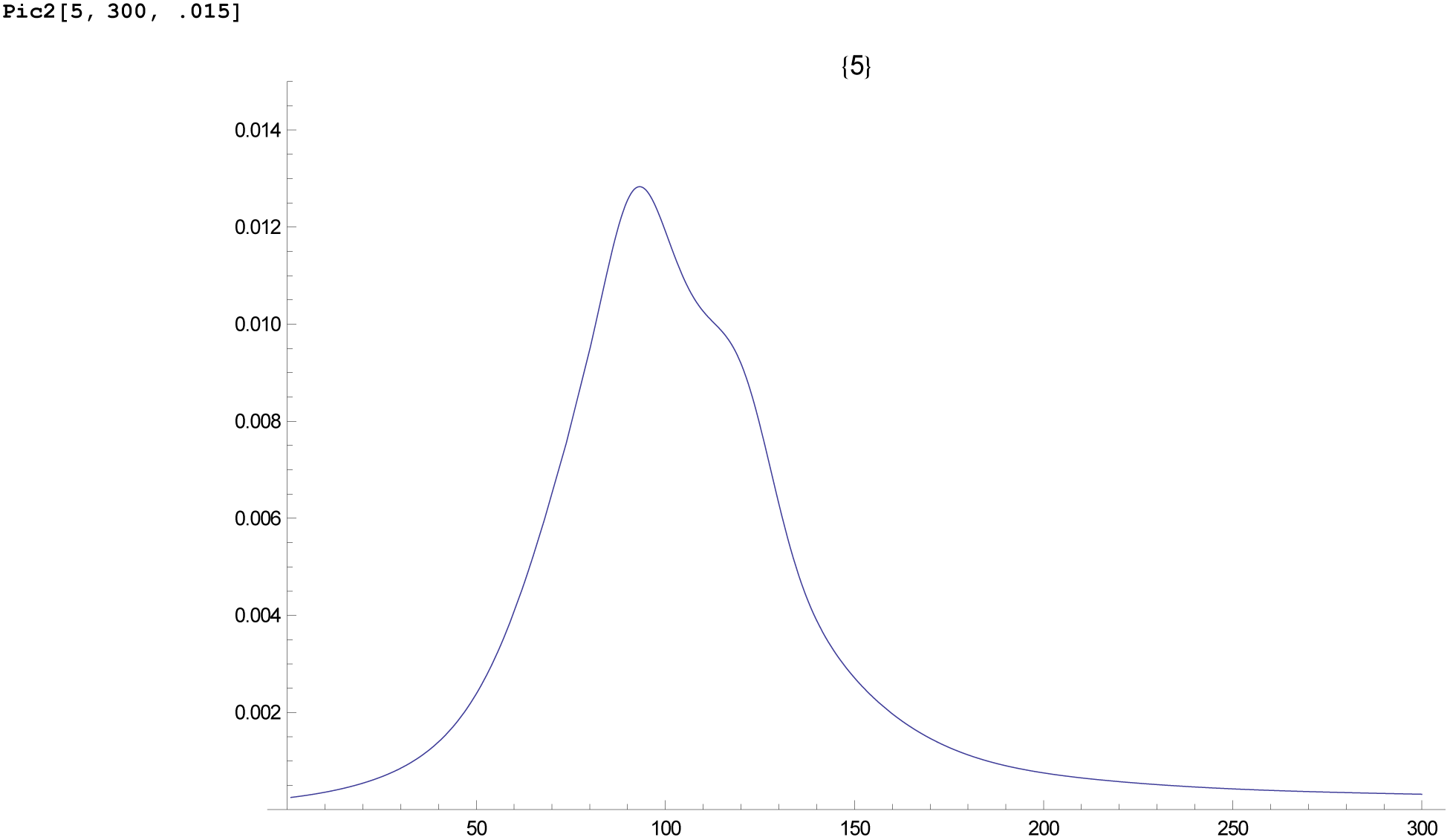

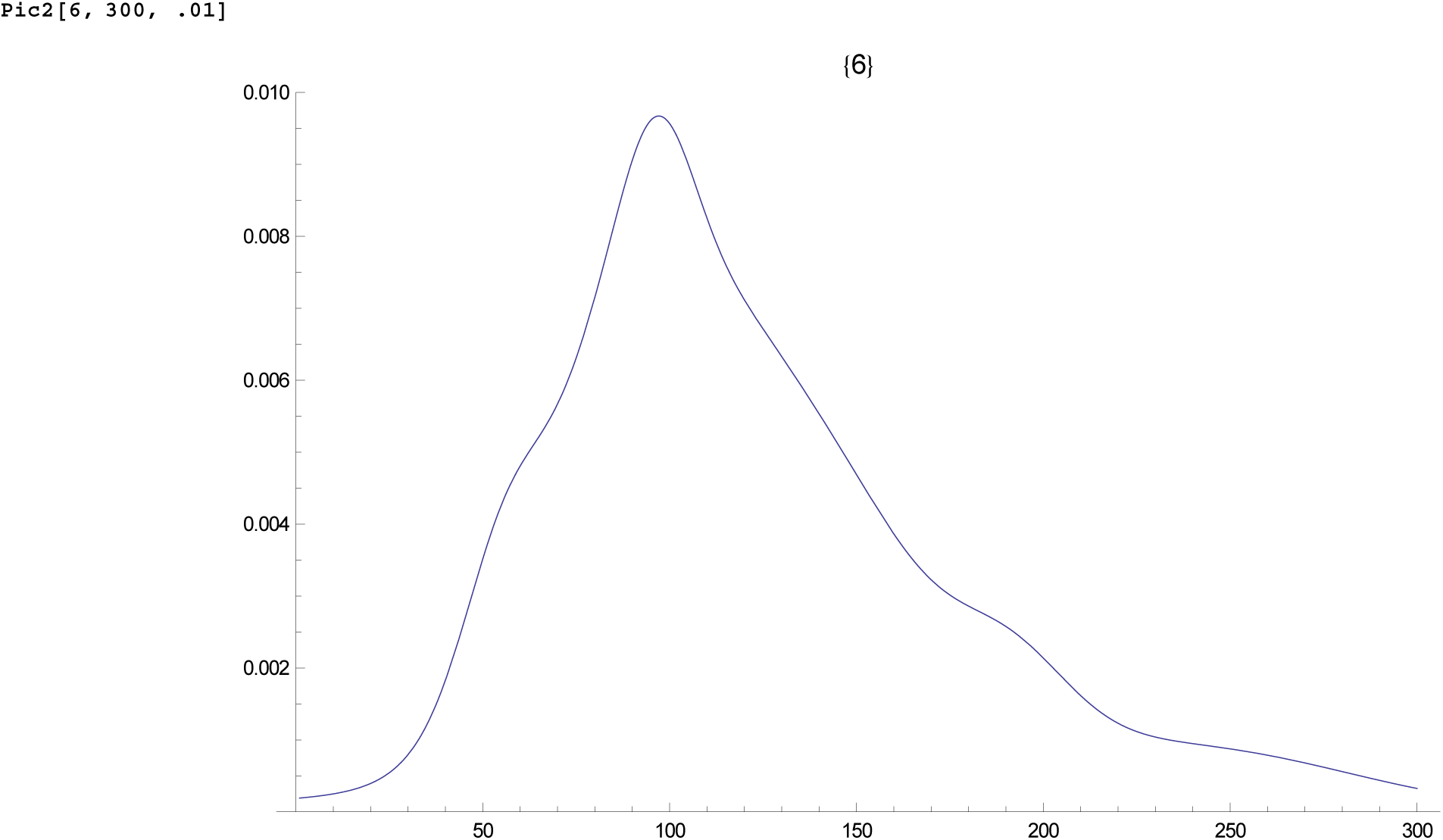

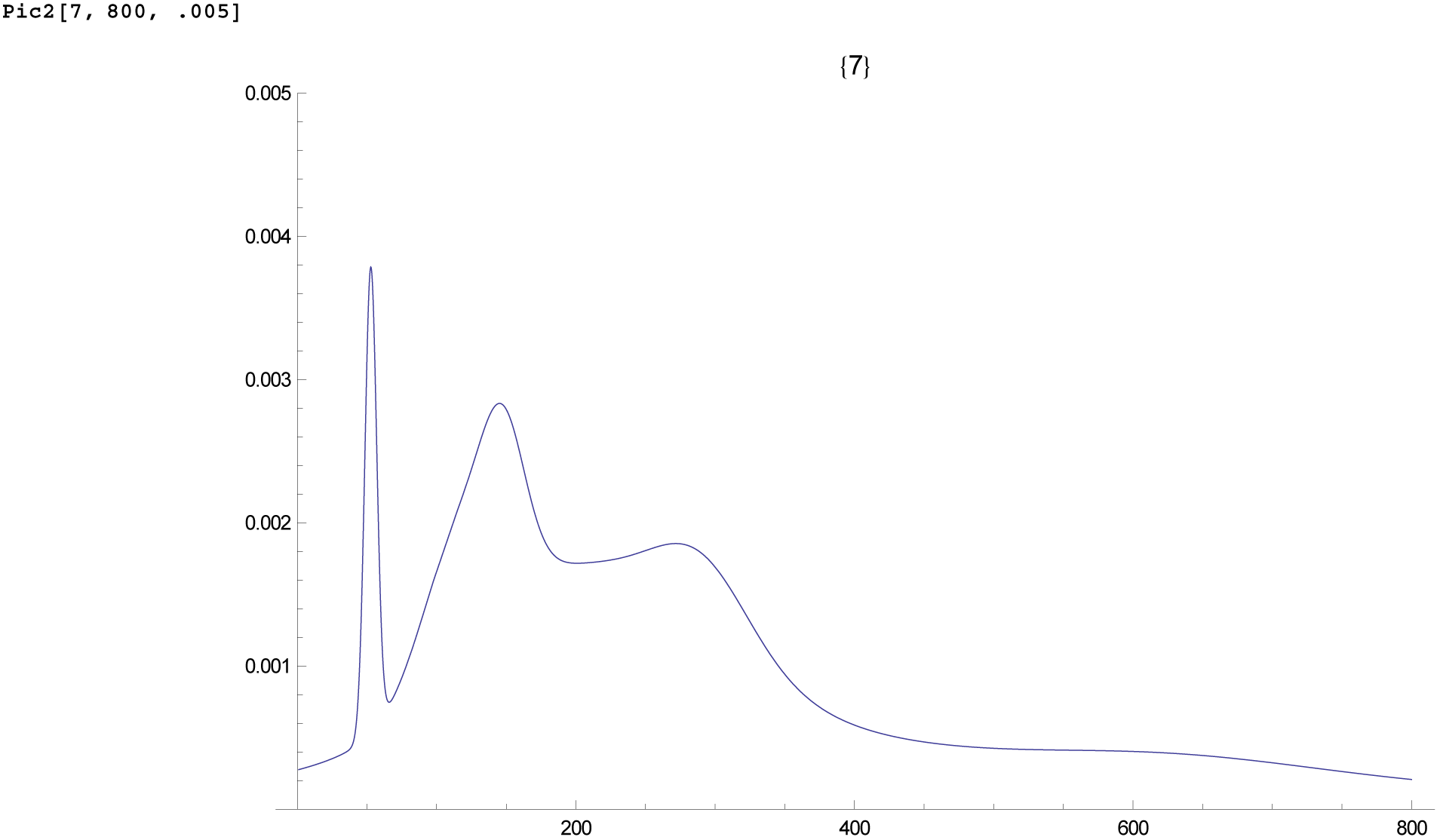

J2: Notice the spike at c50 generations ago, i.e. 650AD, the time of the Arab conquests.

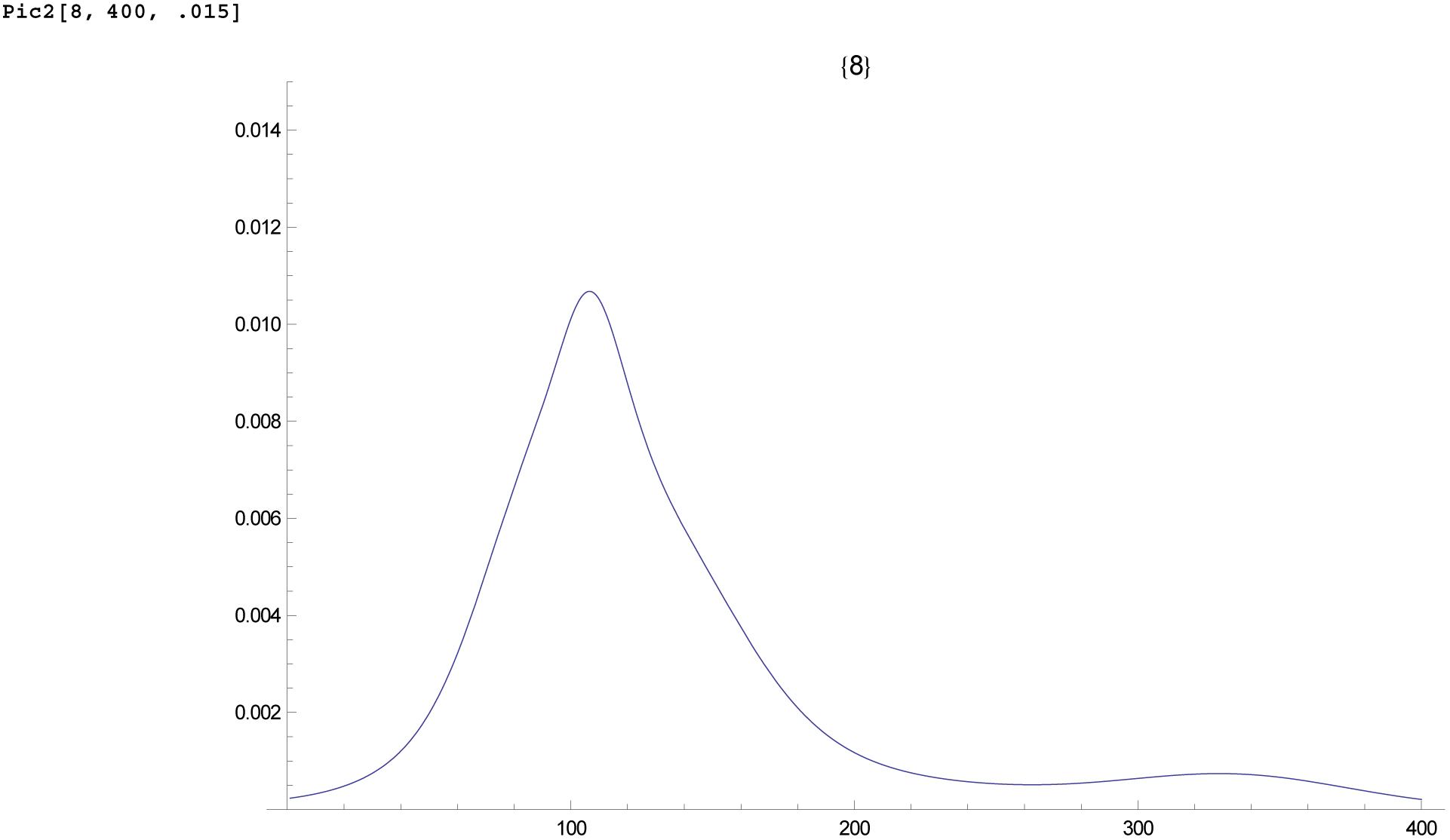

Some idea of the TMRCA is obtained by averaging, either the individual times or the overall, notice the discrepancy

~~~
m1 = Table[ Sum [BB[j] * ZZZ01[[ q, j]], {j, 1, 33}] / 29.0, {q, 1, 8}]
~~~

~~~
{267.84, 288.422, 166.504, 122.94, 119.412, 122.29, 501.842, 133.084}
~~~

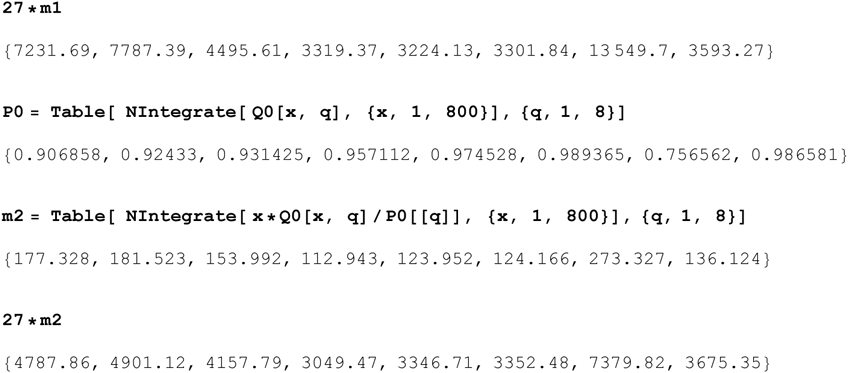

As the averages are not stable we use robust statistics, and stochastic simultations to get the MLE for the TMRCA

For later use we generate random quintiles at roughly 30,40,50,60,70%

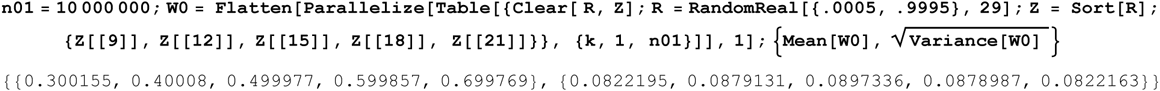

Using the distribution of times ZZZ01 we use bootstrap to generate the same quintiles for our data, inc SD

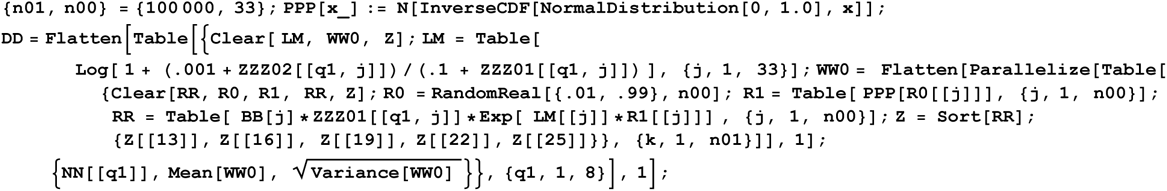

Thus for each file q our data used is the qunitile and its SD

~~~
MatrixForm[DD]
~~~

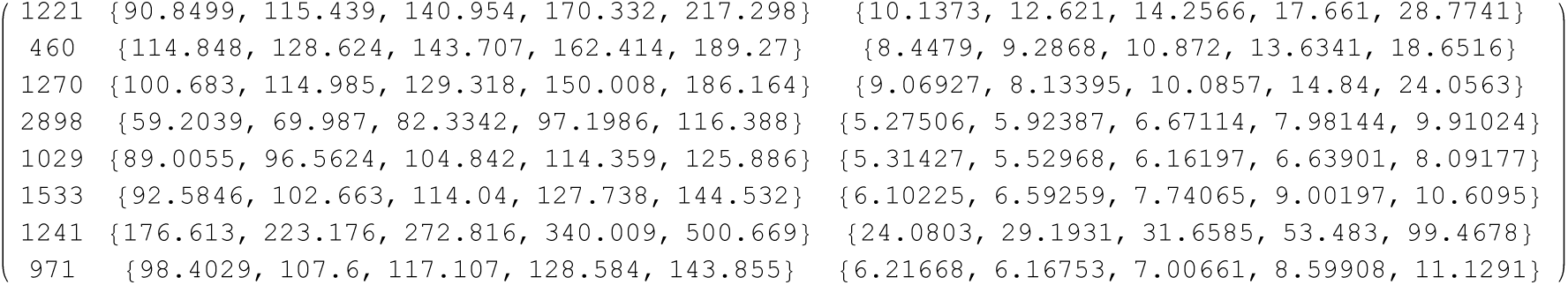

This is now applied to each case, begiining with

We do G2a3 using 5 quintiles

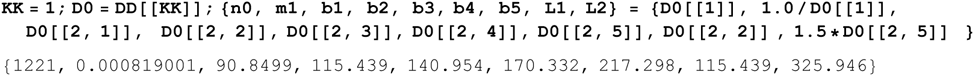

The stochastic simulation uses the branching distribution τo to generate random quintiles. These are filtered by requiring they are within two SD of the data. To speed the process we compiled the branching distribution in files 29ComFun together with interpolation file W29ComFun which must be loaded. This information is contained in the function F4 used. This are the slowest part of the routine, taking about an hour.

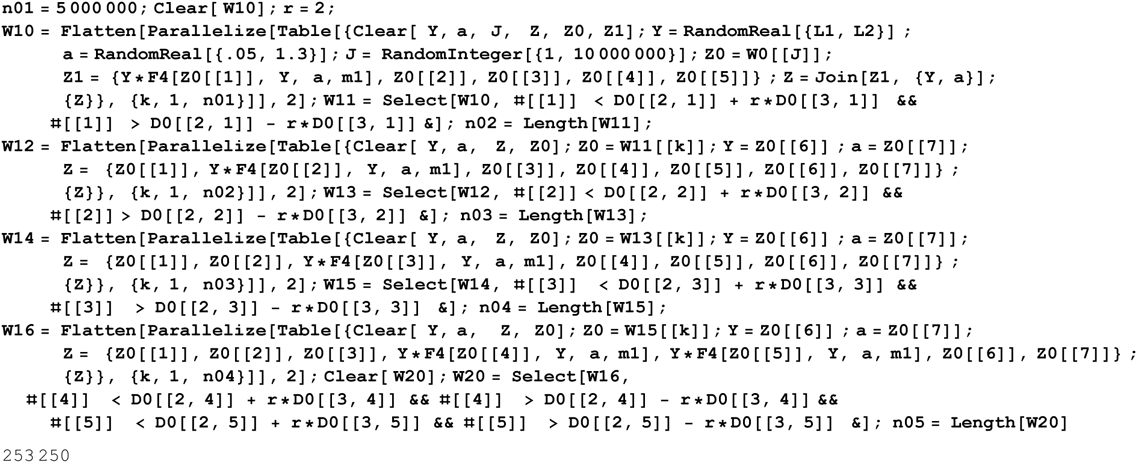

Thus by filtering we now have 253,250 random quintiles within 2 SD of the experimental data. The mean for these is

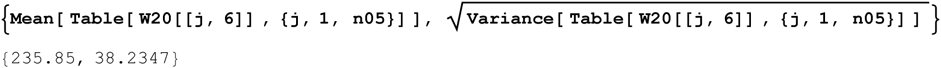

We now use least squares to find a quasilinear estimator which gives the best estimate of the TMRCA for all the random quintiles.

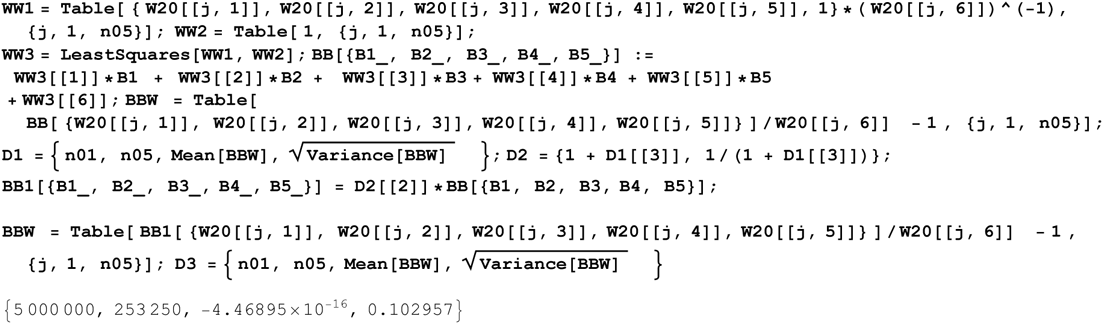

We see the QL estimates the TMRCA with average accuracy −4.46895 × 10^−16^ and overall SD 0.102957. Thus we estimate the TMRCA from our experimental data in generations

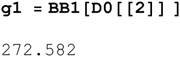

Also we estimate the SD given the variance in the experimental data

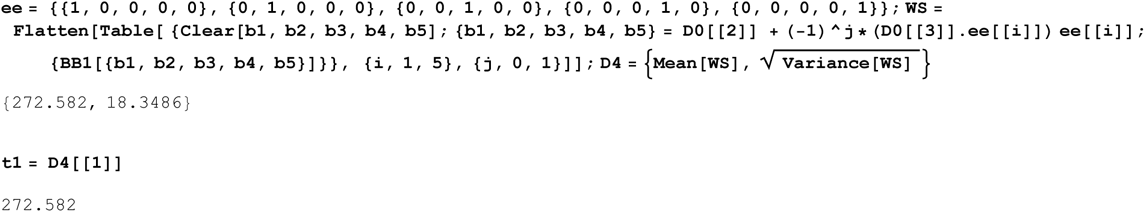

Thus our estimate for the TMRCA in ybp is

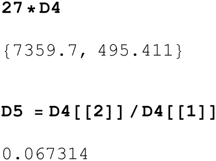

One should not forget that the overall variance is the sum of the variance from sample error and the intrinsic error of the stochastic simulation

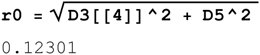

This % error gives SD in years:

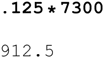

ie for G2a3 we have 5359BC ± 912(1824 at 95% CI)

This is now repeated for each file, but this time without explanation

Next we do R1b1a2 using 5 quintiles

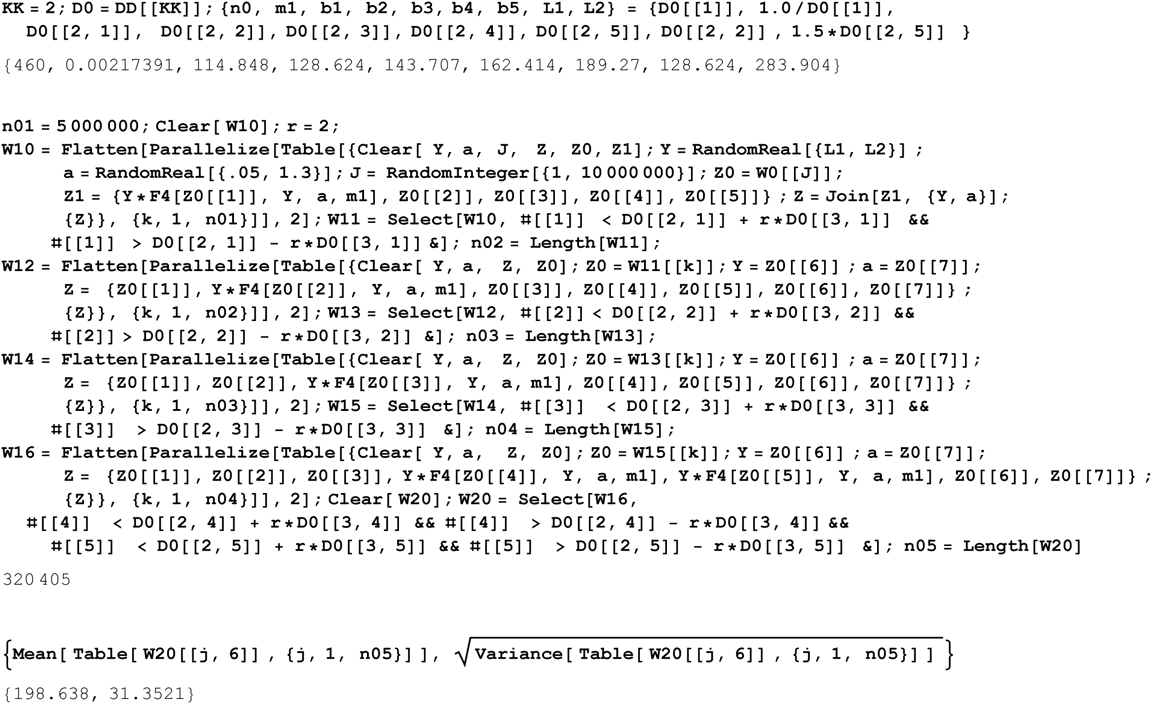

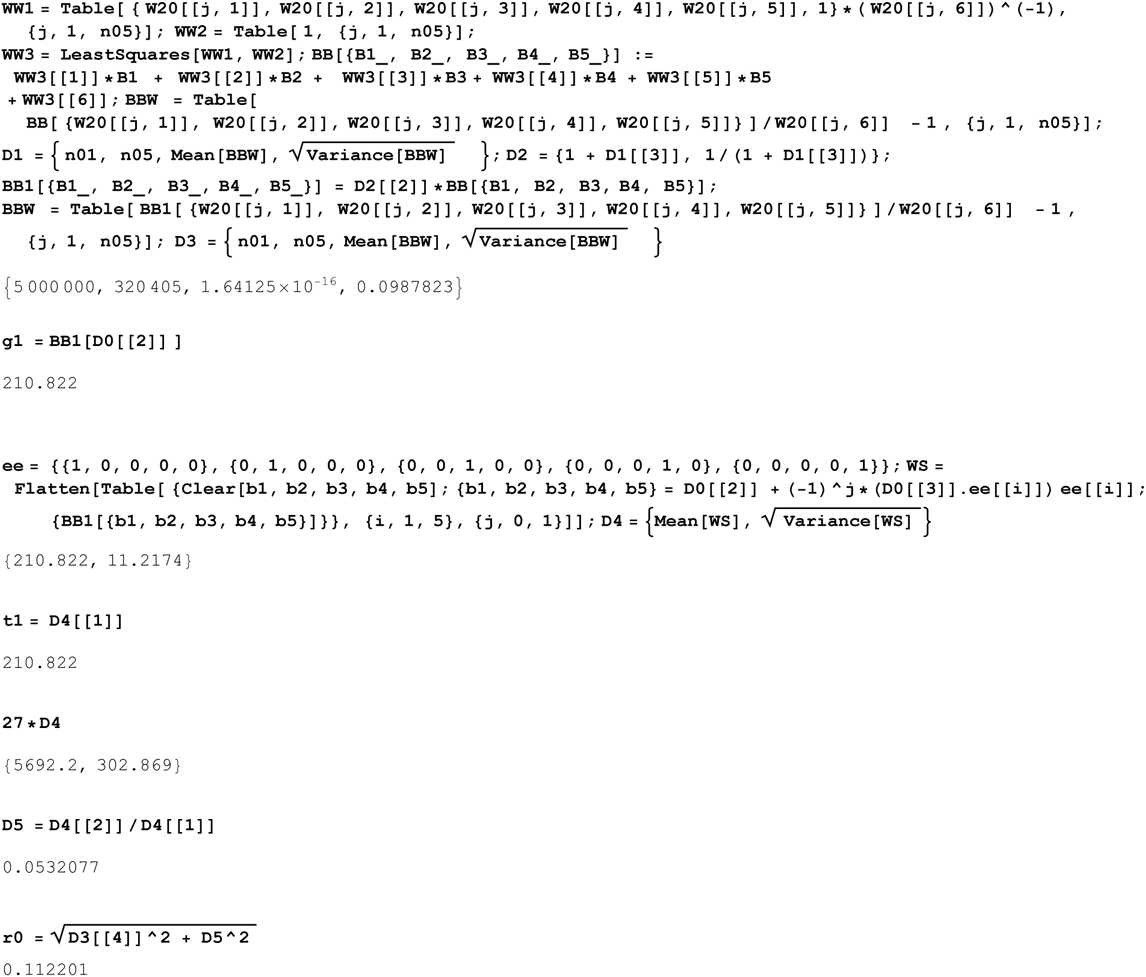

ie for R1b1a2 we have 3700BC ± 625(1250 at 95% CI)

Next we do R1a1 using 5 quintiles

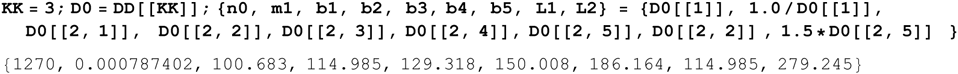

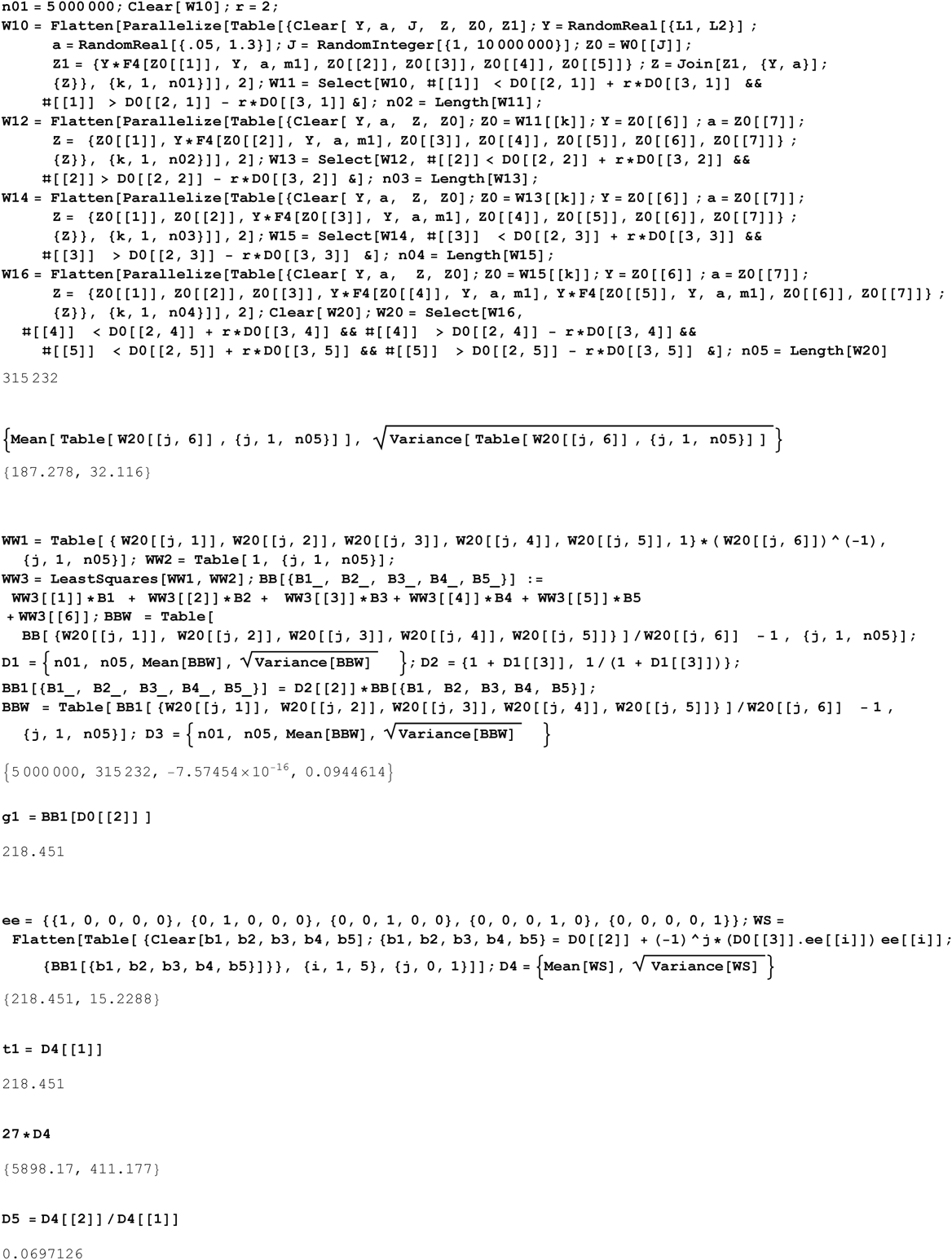

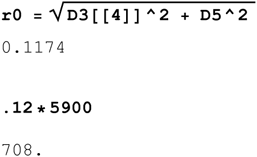

ie for R1a1 we have 4000BC ± 700(1400 at 95% CI)

Next we do I1 using 5 quintiles

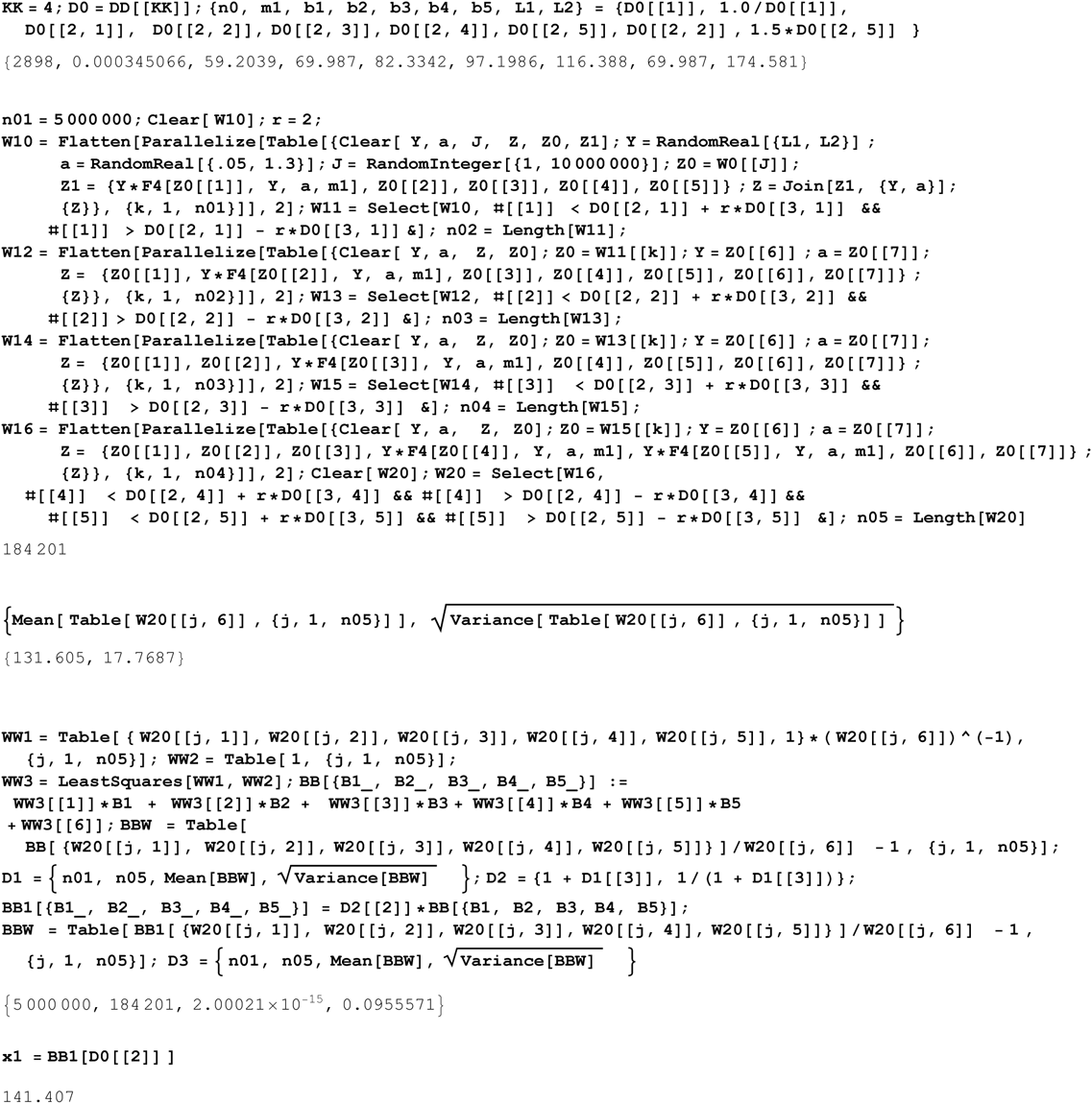

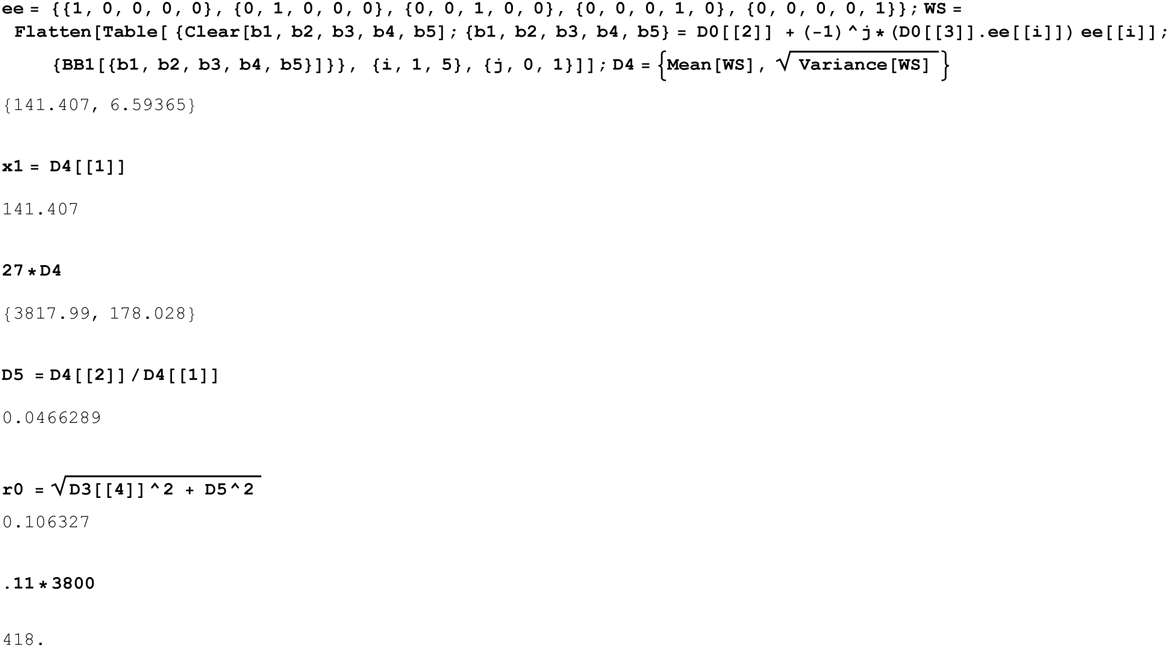

ie for I1 we have 1800BC ± 400(800 at 95% CI)

Next we do L21 using 5 quintiles

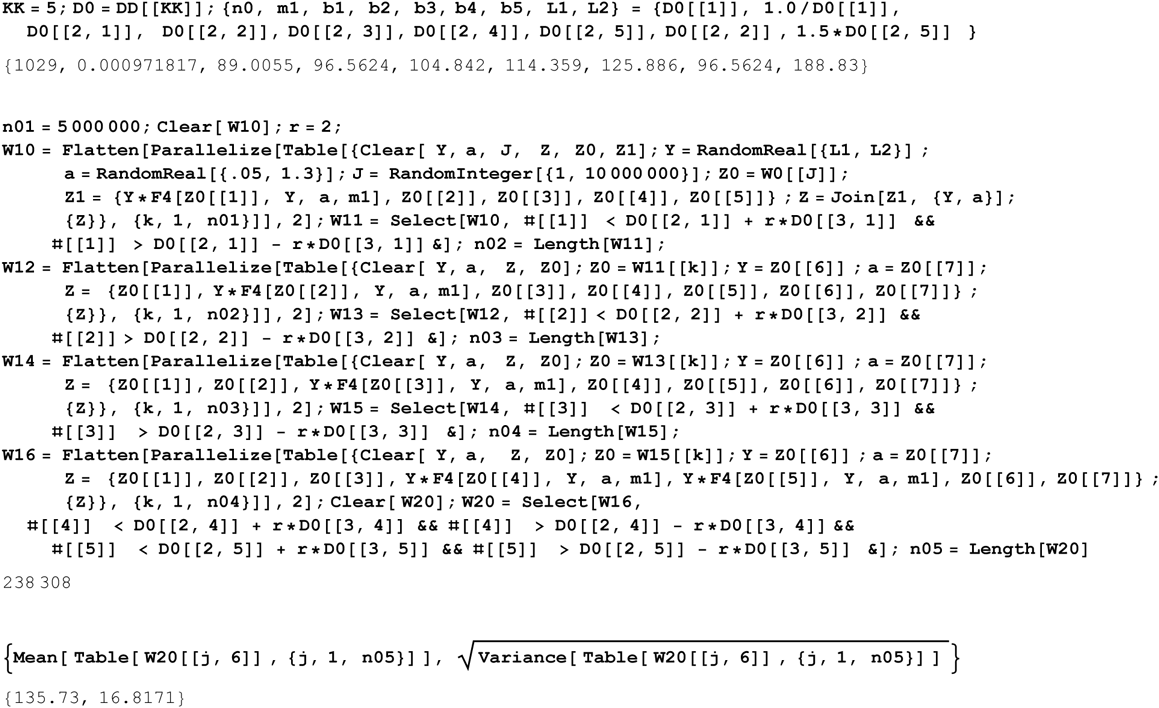

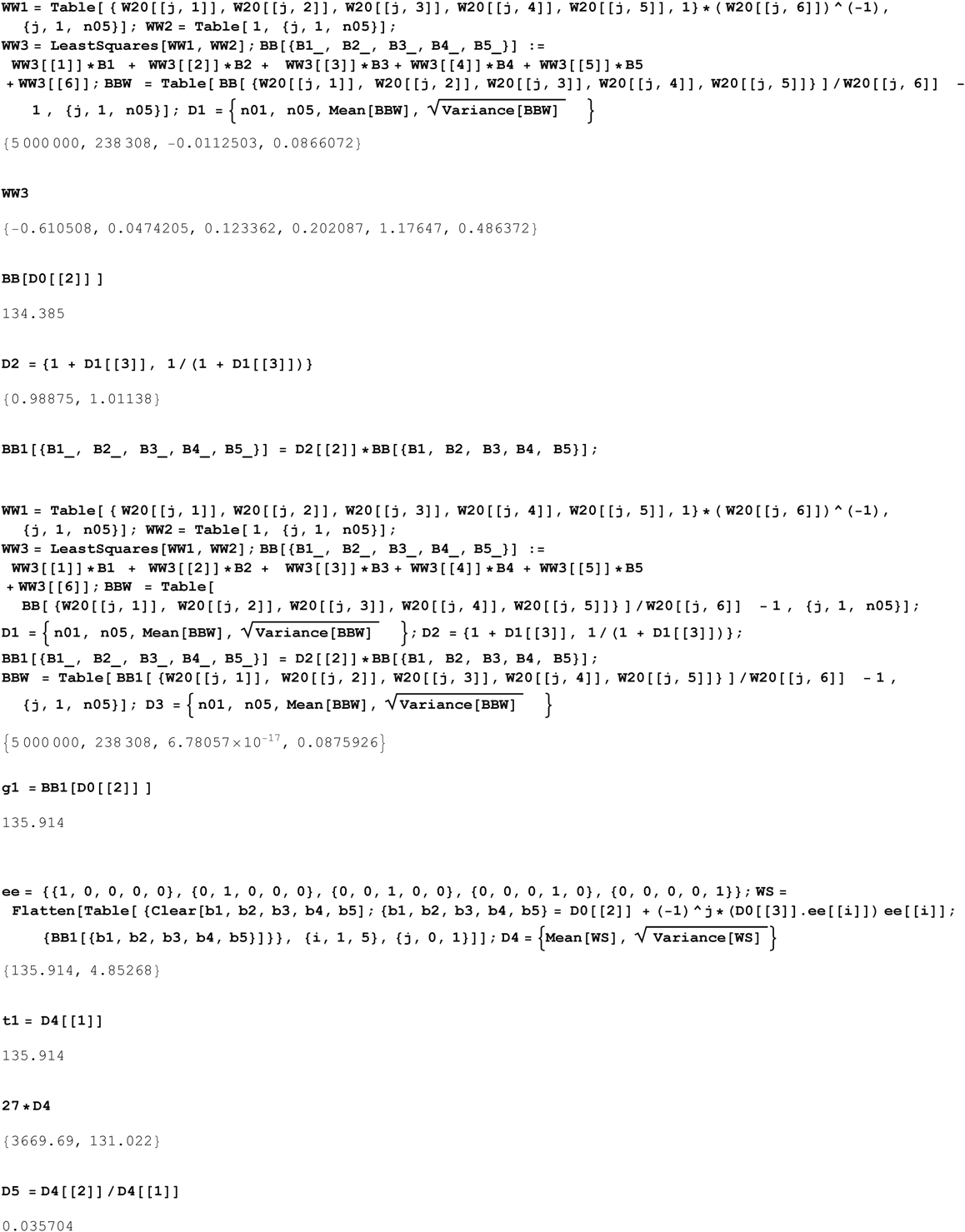

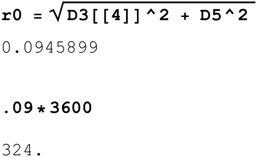

ie for L21 we have 1600BC ± 325(650 at 95% CI)

Next we do U106 using 5 quintiles

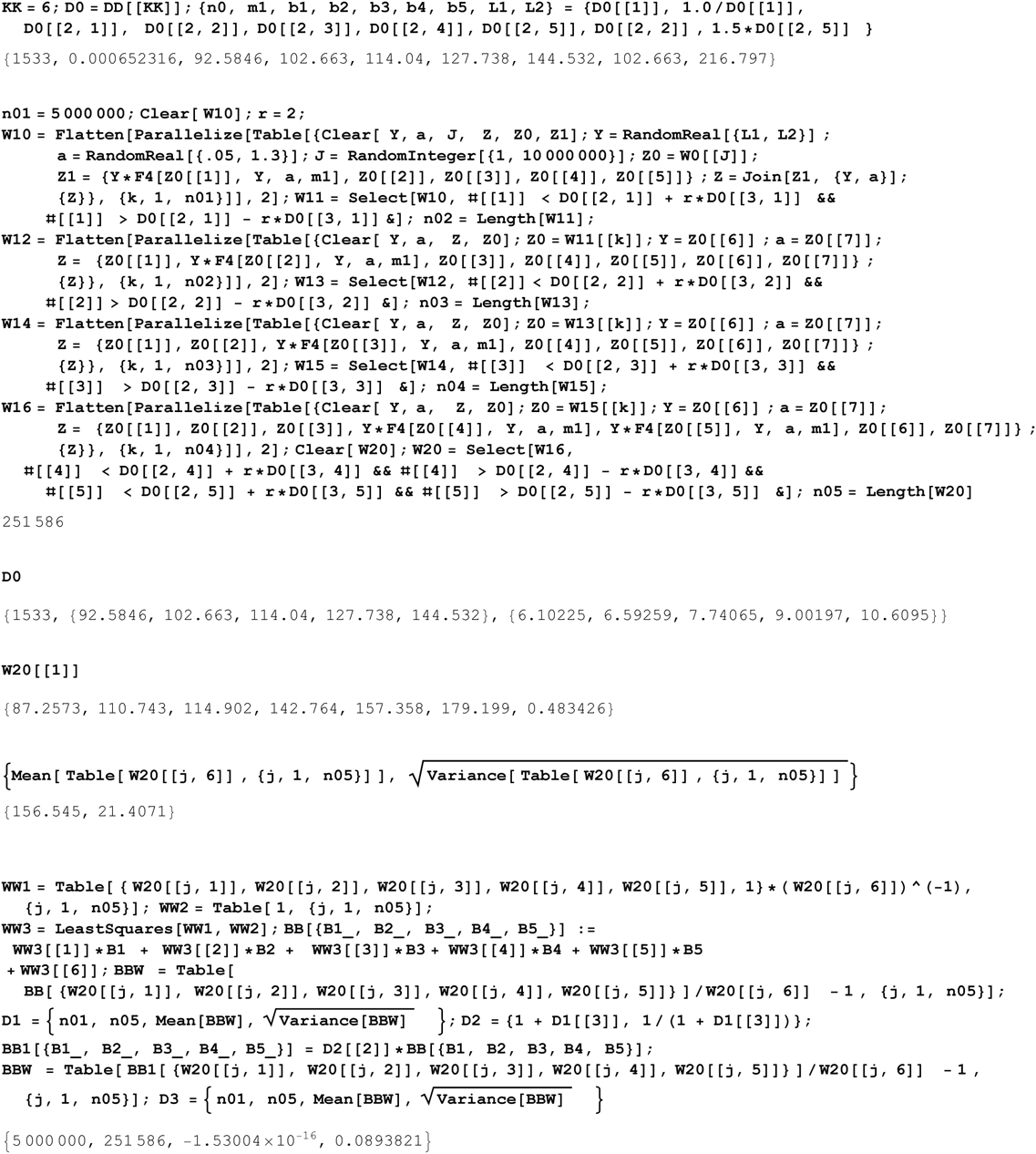

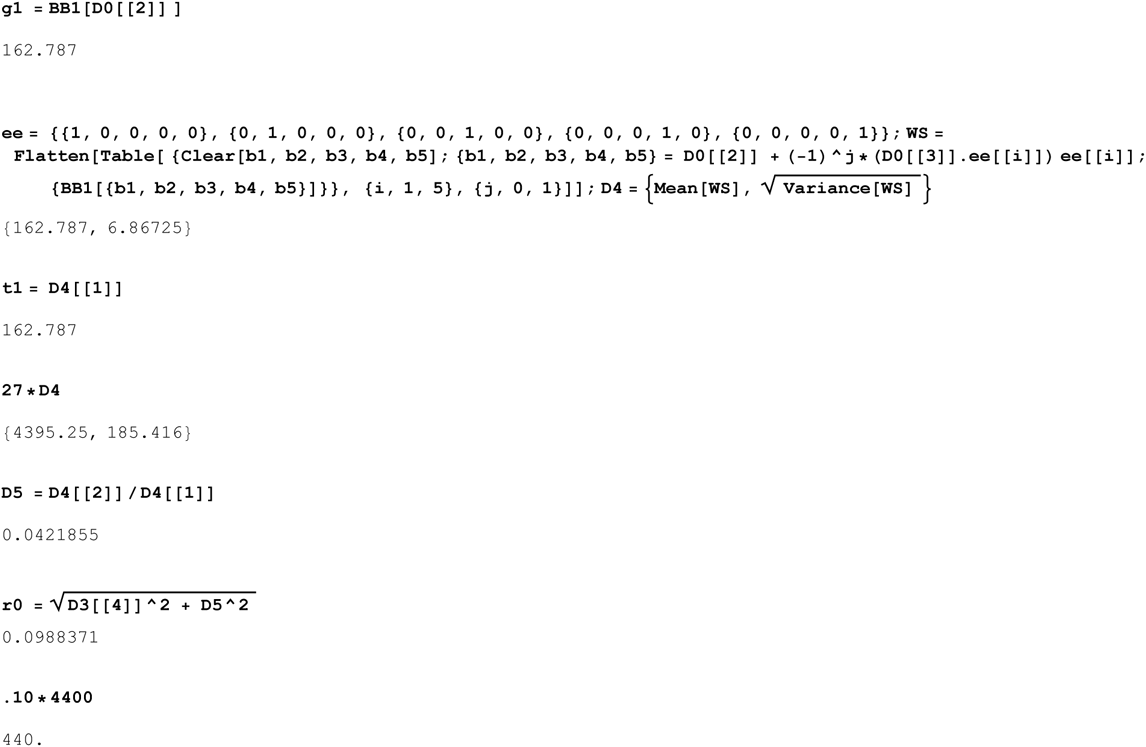

ie for U106 we have 2400BC ± 440(880 at 95% CI)

Next we do J2 using 5 quintiles

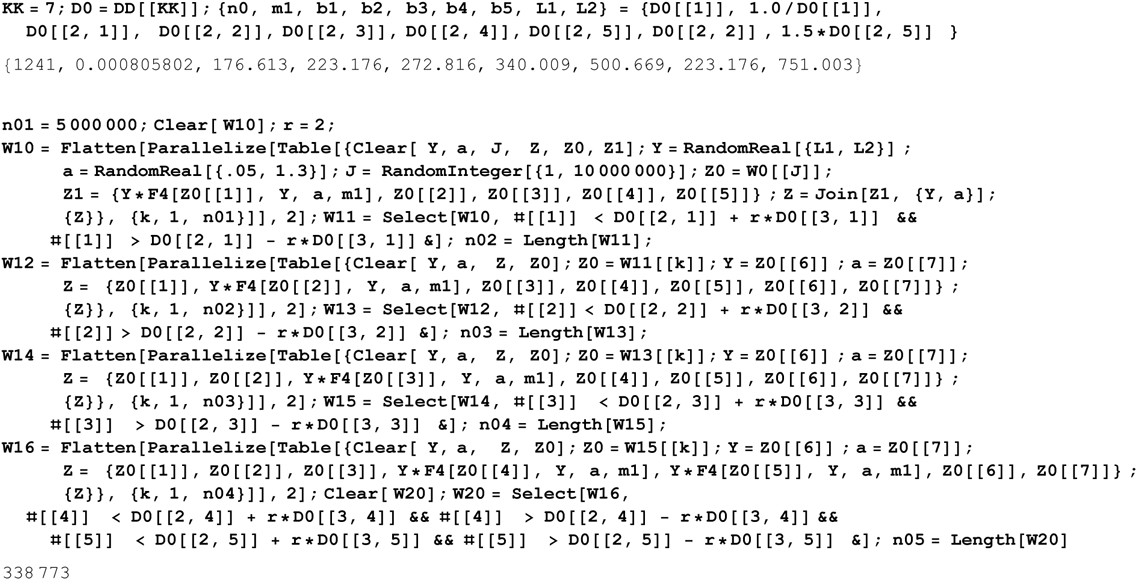

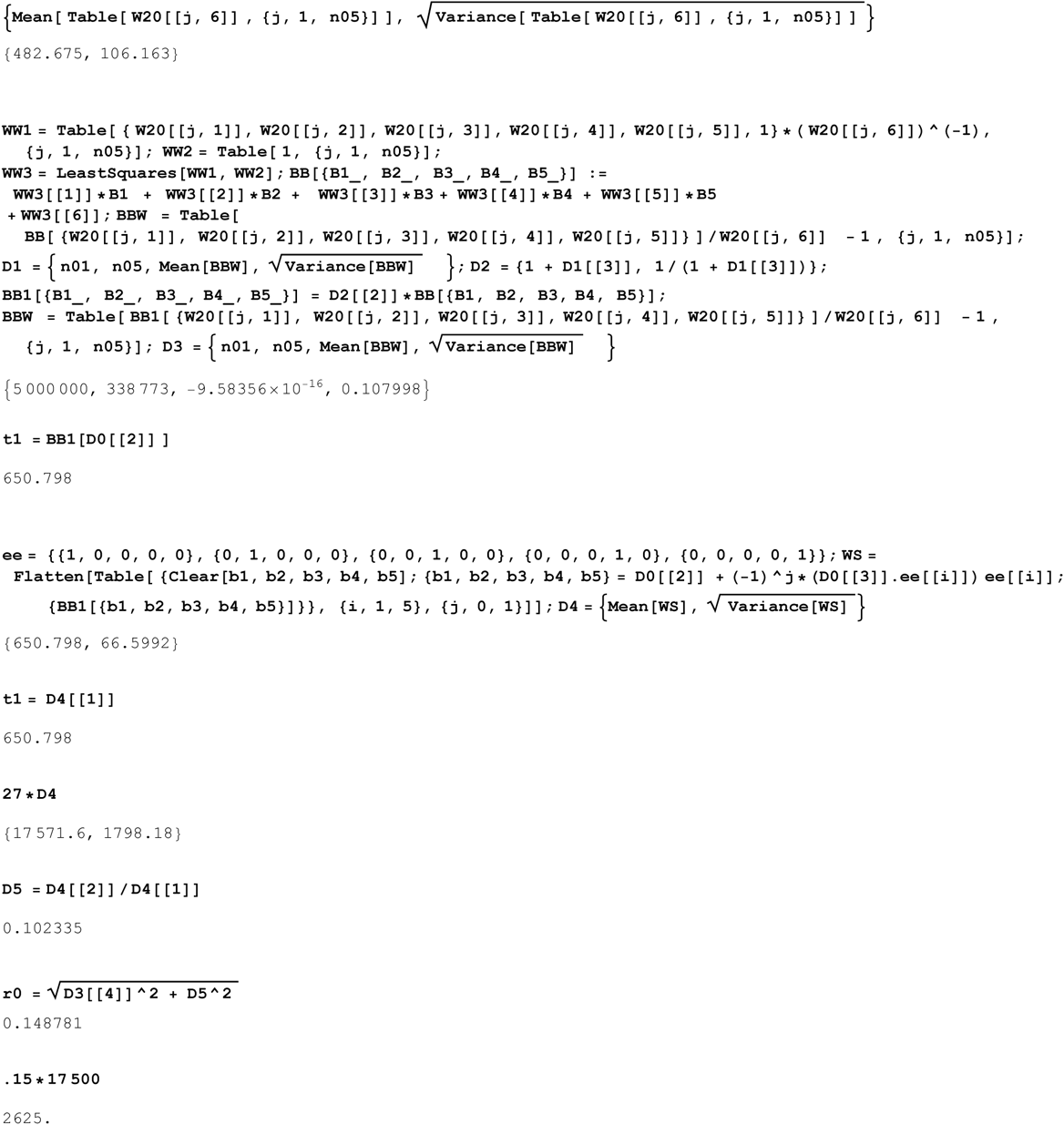

ie for J2 we have 15500BC ± 2600(5200 at 95% CI), this is definitely Paleolithic.

Next we do P312 using 5 quintiles

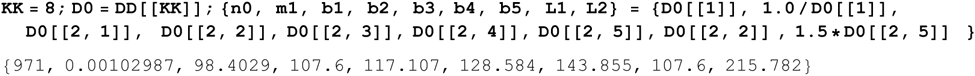

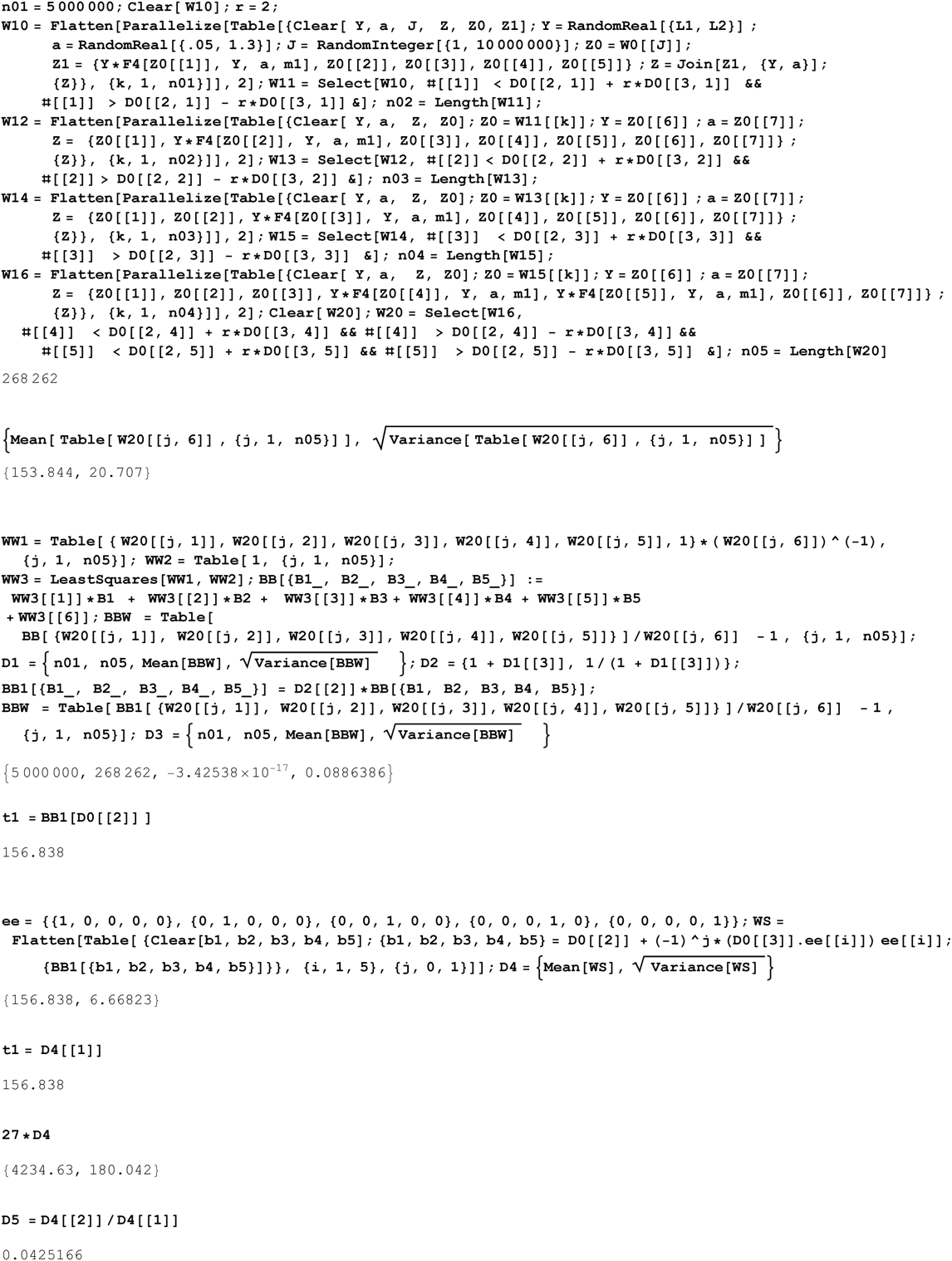

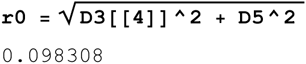

ie for P312 we have 2240BC ± 420(820 at 95% CI)

This information can be summarized by following showing the two means compared with our calc TMRCA and SD

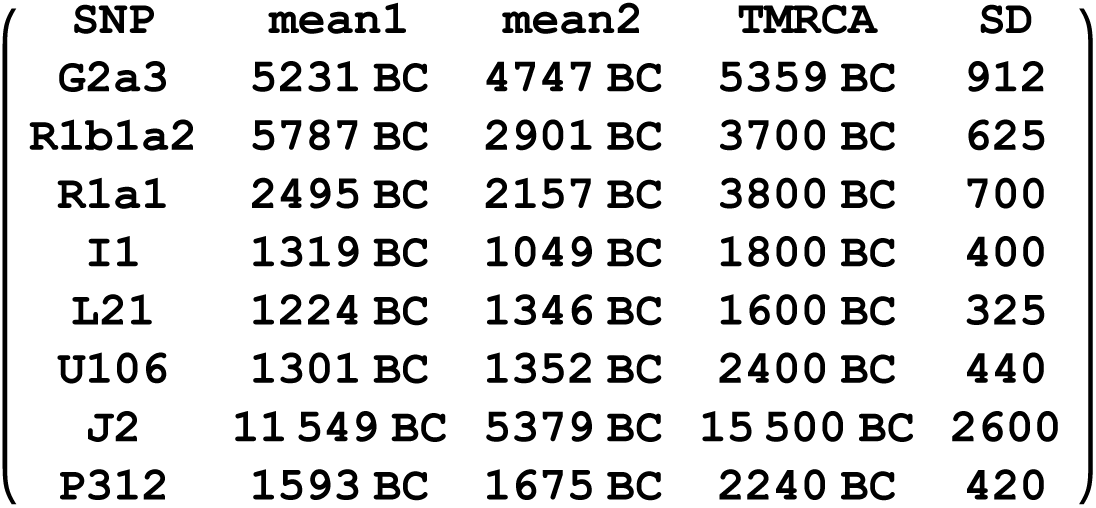

The means only give the right ballpark estimate, usually more than a SD less than the true TMRCA.

